# The Molecular Architecture of Variable Lifespan in Diversity Outbred Mice

**DOI:** 10.1101/2023.10.26.564069

**Authors:** Mohamed Sean R Hackett, Majed Mohamed Magzoub, Tobias M Maile, Ngoc Vu, Kevin M Wright, Eugene Melamud, Wilhelm Haas, Fiona E McAllister, Gary A Churchill, Bryson D Bennett

**Affiliations:** Calico Life Sciences LLC; Krantz Family Center for Cancer Research, Massachusetts General Hospital, and Department of Medicine, Harvard Medical School; The Jackson Laboratory

## Abstract

To unravel the causes and effects of aging we can monitor the time-evolution of the aging process and learn how it is structured by genetic and environmental variation before ultimately testing theories about the causal drivers of aging. Diverse Outbred (DO) mice provide widespread, yet controlled, genetic variation generating considerable variation in mouse lifespan - here, we explore the relationship between DO mouse aging and lifespan. We profiled the plasma multiome of 110 DO mice at three ages using liquid chromatography - mass spectrometry (LC-MS)-based metabolomics and lipidomics and proteomics. Individual mice varied more than two-fold in natural lifespan. The combination of known age and resulting lifespan allows us to evaluate alternative models of how molecules were related to chronological age and lifespan. The majority of the aging multiome shifts with chronological age highlighting the accelerating chemical stress of aging. In contrast, proteomic pathways encompassing both well-appreciated aspects of aging biology, such as dysregulation of proteostasis and inflammation, as well as lesser appreciated changes such as through toll-like receptor signaling, shift primarily with fraction of life lived (the ratio of chronological age to lifespan). This measure, which approximates biological age, varies greatly across DO mice creating a global disconnect between chronological and biological age. By sampling mice near their natural death we were able to detect loss-of-homeostasis signatures involving focal dysregulation of proteolysis and the secreted phosphoproteome which may be points-of-failure in DO aging. These events are succeeded by massive changes in the multiome in mice’s final three weeks as widespread cell death reshapes the plasma of near-death mice.

## Introduction

At a population level, age is the greatest risk factor for all-cause-mortality. This can be directly seen from the phenomenological Gompertz equation whereby hazard rises exponentially, doubling the risk of human death every eight years ^1–3^. Many changes are associated with aging and extreme changes in these hallmarks often manifest in severe disease ^4^. Yet, we still know relatively little about how natural age-associated shifts in hallmarks, either individually or in concert, contribute to the precipitous increase in hazard. In particular, it is not clear which natural age-associated shifts in hallmarks drive aging - to better understand and treat aging we must begin distinguishing the causal drivers of the aging process from its correlated symptoms.

Understanding the relationship between aging and lifespan requires us to measure lifespan, monitor the aging process via direct or indirect (biomarker) readouts of aging processes, and to relate age-associated changes to mortality.

It is generally appreciated that aging is not an ordered regulatory process, but rather the accumulation of physical and chemical defects that organisms combat with both adaptive and maladaptive regulatory responses ^4,5^. Mass spectrometry-based multiomics is a strong platform for probing such a system - proteomics captures realized regulation integrating transcriptional, and post-transcriptional control, while metabolomics and/or lipidomics reveal an organism’s physicochemical state ^6–9^.

Most studies characterizing molecular changes with age utilize model organisms and focus on tissues, which requires culling animals, thereby precluding the measurements of natural lifespan ^6,7,9^. To mitigate this issue, inbred strains can be characterized for both aging and lifespan in genetically identical individuals. In this way a subset of animals can be profiled for natural lifespan, while another set is used for terminal assays at various ages. The results can then be combined to summarize the aging and lifespan of their common strain. This is the prevailing scenario when comparing long- and short-lived strains that allows us to discover molecular changes associated with differential lifespan ^10,11^. Alternatively, a set of strains can be characterized for both age and lifespan to identify consistent associations. This design was used by Williams et al. 2022 to relate the hepatic transcriptomic and proteomic changes across 89 recombinant inbred mouse lines to aging, diet, and strain-specific lifespan ^7^. In this work the authors observed age-associated increases in extracellular matrix transcripts and proteins, and dysregulation of lysosomal proteostasis, particularly cathepsin D (Ctsd), but revealed few associations between lifespan and molecular abundances across strains.

Rather than profiling aging as an endpoint in tissues, measurements of blood cells and/or plasma can be collected with minimal invasiveness. This allows repeated measurements of molecules on an individual basis in a way that can be coupled to natural lifespan ^12^. Rather than exploring a single tissue, relevant biomarkers may be derived from multiple tissues, with changes in circulating proteins generally agreeing with changes in tissues ^9^. Beyond this, the primary way that physically separated tissues interact is through the circulation; thus we have the potential to capture molecules which are mediating non-tissue autonomous aging ^13–15^. This provides the potential for both biomarkers, and possible targets which can be readily profiled in either mice or humans ^16,17^. Ongoing efforts which are part of the UK BioBank, and other epidemiological consortia, are now profiling plasma at the scale of tens of thousands of individuals to relate circulating proteins and metabolites changes to relevant intermediate phenotypes and disease ^18,19^. While such efforts are of great value, human-derived hypotheses derived from observational studies should ultimately be tested with controlled interventions to disentangle causality. To wit, developing models which are representative of human aging physiology but amenable to diverse experimental interventions and deep profiling is a key enabler of translational medicine.

Diversity Outbred (DO) mice are outbred mice derived from eight commonly used strains of inbred mice ^20^. These founder strains vary in median lifespan by almost two-fold with short-lived strains dying from various cancers and long-lived strains exhibiting a pronounced anticorrelation of circulating Igf1 levels and lifespan ^21–23^. Cohorts of DO mice are particularly valuable for understanding how genetic variation shapes intermediate phenotypes which may cascade into functional outcomes such as lifespan. In DO mice environmental variation can be controlled, and genetics can be used as a tool for identifying the molecular drivers of complex traits ^20,24^. This potential for systems genetics has been realized by several recent studies looking at how dietary intervention’s influence is contextualized by DO genetics to impact tissue ‘omics and functions. This approach was first adopted by profiling 192 DO mice’s hepatic transcriptome and proteome revealing that both local and distant molecular Quantitative Trait Loci (QTLs) could be discovered, and that many protein QTLs’s effects on protein abundance are mediated through their cognate transcript ^25^. Establishing a more direct association between molecular changes and tissue function, in brown adipose tissue, measurements of 163 DO mice’s proteome identified regulators modulating thermogenic potential ^26^. Similarly, measurements of the skeletal muscle proteome of 215 DO mice were paired with metabolic measures to identify proteins associated with variable insulin resistance ^27^. Studies such as this which explore how tissue functions vary within DO mice are complemented by studies understanding how aging may interact with this underlying functional heterogeneity to drive gross organismal decline. The impact of aging on the DO mouse kidney transcriptome and proteome was explored in Takemon et al. 2021 ^6^ where they noted specific changes in podocyte cytoskeleton remodeling to maintain glomerular filtration rate as well as a global disconnect between age-associated transcriptional and protein changes reflecting altered proteostasis.

Here, we build upon these amassing atlas’ of DO physiology by profiling the plasma of 110 DO mice (55 males and 55 females) longitudinally at three ages (8, 14, & 20 months), roughly corresponding to adolescence, early adulthood and late adulthood, using liquid chromatography - mass spectrometry (LC-MS) metabolomics and lipidomics and tandem mass tag (TMT) proteomics (Figure 1). Following each mouse’s final blood-draw, their natural lifespan was determined, with profiled mice living between an additional two and 821 days. Near death, mice exhibited a widespread “death’s door” signature while the broader aging process is best described by three interacting aging archetypes following distinct kinetics. Most changes in the metabolome and lipidome track with chronological age with subtle shifts from 8 to 14 months and more pronounced changes progressing to 20 months. In contrast, most changes in the proteome, particularly changes in proteostasis and inflammation which are canonical features of the aging proteome ^4,28–30^, shift with “fraction of life lived”. This measure, the ratio of chronological age to lifespan, may be capturing the biological age of DO mice which is only loosely correlated with chronological age. A third archetype, “age x lifespan” interactions, are best described as a loss-of-homeostasis. These failures lead to punctuated accumulation of proteosomal components and targets of the Golgi kinase Fam20c, which may be key points-of-failure in the aging process leading to a broader death’s door signature and ensuing death.

**Figure 1:**
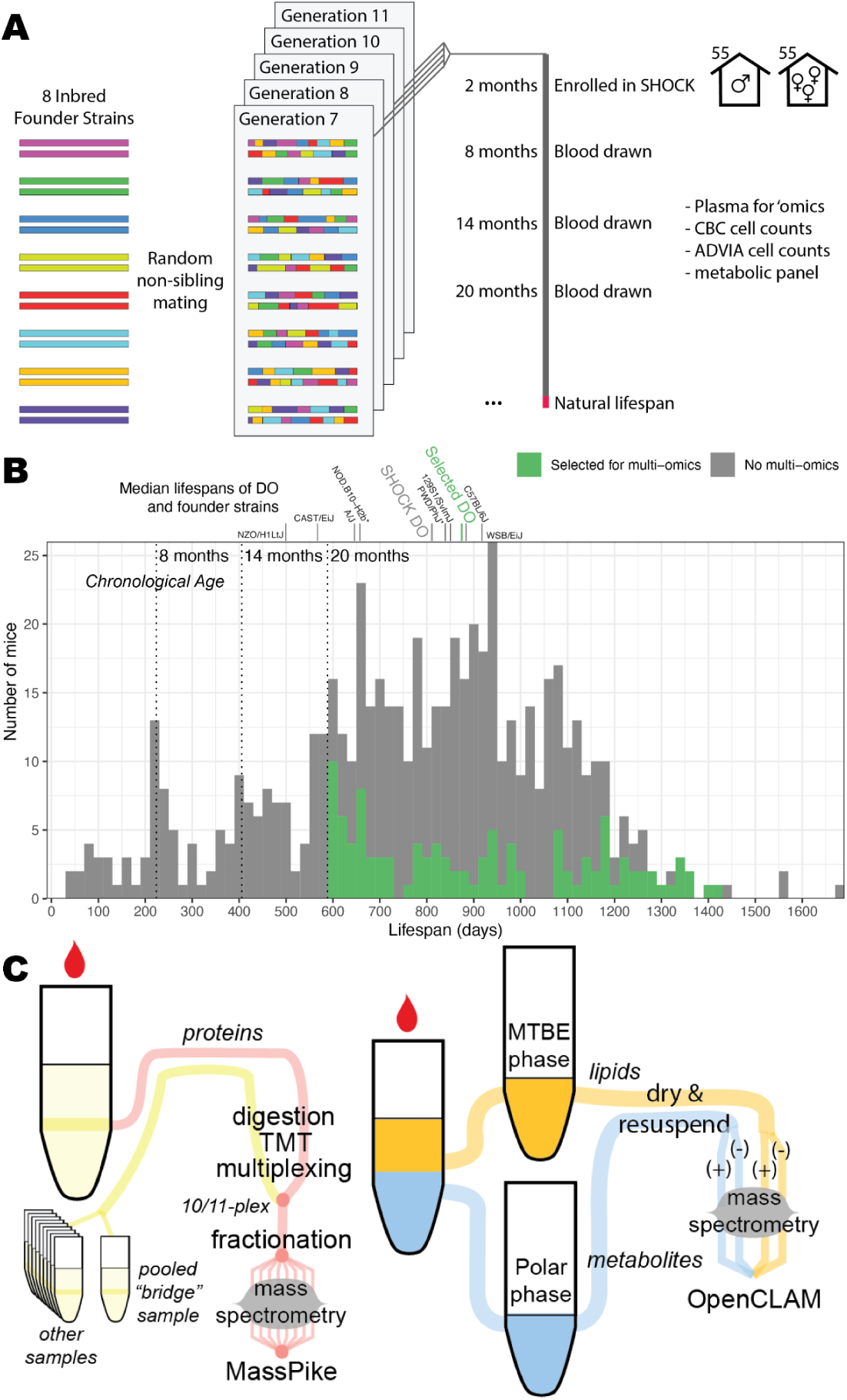
Profiling the plasma multiome of aging Diversity outcross (DO) mice with known lifespans. **(A)** DO mice were generated from multiple generations of random mating of inbred lines derived from common mouse strains. Mice from five DO generations were enrolled in the SHOCK cohort at 2 months, blood was drawn at 8, 14, and 20 months, and mice were then housed for the remainder of their natural life. **(B)** DO mice vary greatly in their natural lifespan. Mice selected for this study were restricted to those with three blood draws and further enriched for short- and long-lived mice. The DO cohort’s eight founder strains’ median lifespan are shown. Two strains, labelled with asterisks, are not an exact match to the founder strains. **(C)** Plasma samples were separately processed for proteomics and metabolomics/lipidomics. 9-10 experimental proteomics samples and one bridge sample which was shared across batches were labelled with distinct isobaric TMT tags and combined into a single physical 10/11-plex. Each plex was fractionated and injected into the mass spectrometer where abundances of individual samples were detected by their distinct MS3 reporter ions. Metabolites and lipids were extracted from a common plasma sample through a biphasic extraction. Following drying and resuspension, metabolites and lipids were separately analyzed in positive- and negative-mode by mass spectrometry.

## Results

### Dataset generation

The SHOCK study conducted at The Jackson Laboratory aimed to identify the genetic and physiological determinants of lifespan in an outbred population of mice. 600 mice, drawn from generations 7 to 11 of the broader DO breeding project, were enrolled in the SHOCK study (Figure 1A). These mice were raised in standardized housing and fed a chow diet from birth until their natural death. The median lifespan of the eight inbred founder strains of the DO cohort vary by about 70% with DO strains showing average lifespans typical of long-lived mouse strains but with greater lifespan variability than inbred strains (Figure 1B) ^21^. Each mouse was intermittently phenotyped including by drawing blood at 8, 14, and 20 months of age. Blood samples were immediately profiled using a panel of assays including complete blood count (CBC), which assesses the composition of cells in the blood. Plasma was then prepared from each blood sample and frozen for subsequent multiomic profiling.

To explore molecular readouts of aging and lifespan, we selected 110 mice with three available blood draws for deeper analysis using liquid-chromatography mass-spectrometry (LC-MS) multiomics (Figure 1B). Selected mice were enriched for extreme lifespans to improve the power of detecting lifespan-associated molecules. By profiling the same mouse at multiple ages using a longitudinal analysis, we are able to explore trends that occur within individual mice. This allows us to correct for mouse-specific variation when identifying molecules associated with chronological age, or age x lifespan interactions. Such interactions may occur if the change in a molecule between timepoints is more predictive of lifespan than the molecule’s baseline level. This design also allows us to define alternative aging-relevant measures: lifespan remaining (lifespan - age), and fraction of life lived (FLL; age / lifespan).

We profiled each samples’ biomolecules using LC-MS Tandem Mass Tag (TMT) proteomics, metabolomics (positive- and negative-mode) and lipidomics (positive- and negative-mode) (Figure 1C). Profiling small molecules in both ion-modes using separate LC-MS runs allows us to recover compounds that more readily form either positively- or negatively-charged ions, increasing the overall coverage of the dataset. We profiled samples across two tranches of 54 (set 1/2) and 56 mice (set 3). Despite using equivalent chromatography and mass spectrometry methods across both tranches, the considerable time between batches brought challenges in recovering a single coherent dataset. A detailed overview of the bioinformatics approach can be found in the materials and methods and the project’s [GitHub](https://github.com/calico/hackett-doomics) repository.

Briefly, we separately processed metabolomic and lipidomic positive-, and negative-mode datasets using OpenCLAM. Metabolite and lipids were manually curated using MAVEN to facilitate biological interpretation ^31^. Technical “injection” replicates of set1/2 metabolites were profiled and since replicates were highly correlated they were skipped for the remainder of the dataset (Figure <u>SF1</u>). Metabolomics datasets could be aggregated across tranches by aligning across unambiguous masses and shared fragmentation patterns. This facilitated subsequent analysis of metabolomic unknowns (i.e., metabolites with a known mass and retention time but unknown identity). The positive- and negative-mode lipidomics results could not be aligned between set1/2 and set3 due to non-monotonic shifts in retention time. So, each dataset was separately processed and manually identified lipids were merged across sets based on shared labels. This created a single coherent dataset but limited analysis of unknowns. We did manually retain 21 unknown lipids identified during preliminary untargeted analyses - these features were carefully treated to avoid ascertainment bias (see methods). Metabolite and lipid features were normalized by mean-centering by day (within a set) and were mean-centered again between set1/2 and set3. This batch centering approach, compared to other normalization approaches explored, greatly decreased variability between cross-day technical replicates while preserving variability between experimental samples (Figure <u>SF2</u>).

TMT proteomics combines multiple samples whose proteins have been labeled with different compatible TMT tags into a single physical sample called a “plex”. Once combined, each sample in a plex’s peptides will possess the same exact mass, retention time, and MS2-based fragmentation pattern but each TMT tag will fragment differently at the MS3 level to reveal reporter ions whose intensity is proportional to samples’ peptide concentrations ^25,32^. Since samples are pooled, technical variability covaries within a plex allowing for robust estimation of relative abundances between pairs of tags at the peptide or protein level. By including a shared “bridge” sample in each plex, we were able to aggregate relative abundances across multiple plexes with the same bridge channel ^26^. This is conceptually similar to using a common reference sample in a two-color microarray experiment ^33^. After aggregating peptides of the same protein ^34^, relative abundances within each plex were centered within each set to correct for systematic differences in each set’s bridge channel (Figure <u>SF3</u>).

### The multiome is shaped by age, sex, and looming death

Using measurements of ∼2,200 distinct features across 330 samples (Table 1, Table <u>ST1</u>), we first applied exploratory data analysis to identify the factors shaping the aggregated plasma multiome. These sources of sample-level variation may be due to the differences in age, sex, lifespan, or genetic variation which were intended aspects of our experimental design. Alternatively, they may be due to unintended sources of error.

**Table 1:** Measured features across data modalities.

| Data Modality | # of Identified Features | # of Unknowns |
| --- | --- | --- |
| Proteomics | 972 | 0 |
| Metabolomics | 160 | 611 |
| Lipidomics | 403 | 21 |

Applying principal components analysis, we identified and removed four data-modality or mode-specific (e.g., just affecting positive-mode metabolomics) outlier samples whose inclusion drove the leading principal components. Outlier samples at the data modality level could reflect problems with sample preparation, while mode-specific outliers generally reflect issues with chromatography (most of which were addressed with LC-MS re-runs). More problematic, we identified two patterns which are clearly shaped by blood draw date and mouse generation (Figure <u>SF4</u> and Figure <u>SF5</u>). One batch effect, the “early G8 effect”, impacts all of the 8 month blood draws from generation 8 and the first half of the 14 month blood draws (by draw date). The other effect, the “late blood draw (LBD) effect”, impacts all samples drawn after Nov 9th 2013 which includes the 20 months draws from G9, 10, and 11 and the latter half of the 14 month blood draws for G11. Both effects could be clearly captured as categorical covariates (Figures <u>SF6</u>, <u>SF7</u>) but due to their confounding with age, these batch effects were estimated during feature-level regressions rather than regressing them out up-front. For the remaining results, both the early G8 and LBD batch effects are regressed from the data but these two batch effects’ impact on individual features can be visualized in our [R Shiny application](http://connect.calicolabs.com/doresults). All results presented below have been spot-checked for consistency across generation, mass spec tranche (set1/2 vs. set3), and known batch effects. To identify other technical batch effects we explored the extent to which experimental batches structure the leading principal components in individual datasets and we verified that such effects were minor (Figure <u>SF8</u>).

After correcting for major undesirable sources of variation, a group of samples from near-death individuals with profoundly altered ‘omics stand out (Figure 2A, B). This near-death effect is observed across data modalities but most profoundly affects the metabolome hence we termed it the death’s door metabolite (DDM) signature. While mortality does increase following a blood draw, the number of animals in this group is similar to what we would expect from the overall increase in hazard independent of blood draw and thus we are likely monitoring the natural death of these animals. To convey how molecularly unusual these samples are, their Mahalanobis distance relative to the principal components’ ellipsoid, a multivariate generalization of a Z-statistic, was found and this measure clearly distinguishes these unusual samples (Figure 2C) ^35^. This analysis allowed us to define a cutoff for “DDM samples” as those where the mouse died less than 21 days after the final blood draw. This summary helps to both characterize the molecular makeup of the DDM and to exclude these samples when characterizing aging (they are exclusively 20 month blood draws) and lifespan (they are the shortest-lived mice) since they would drive ungeneralizable associations.

**Figure 2:**
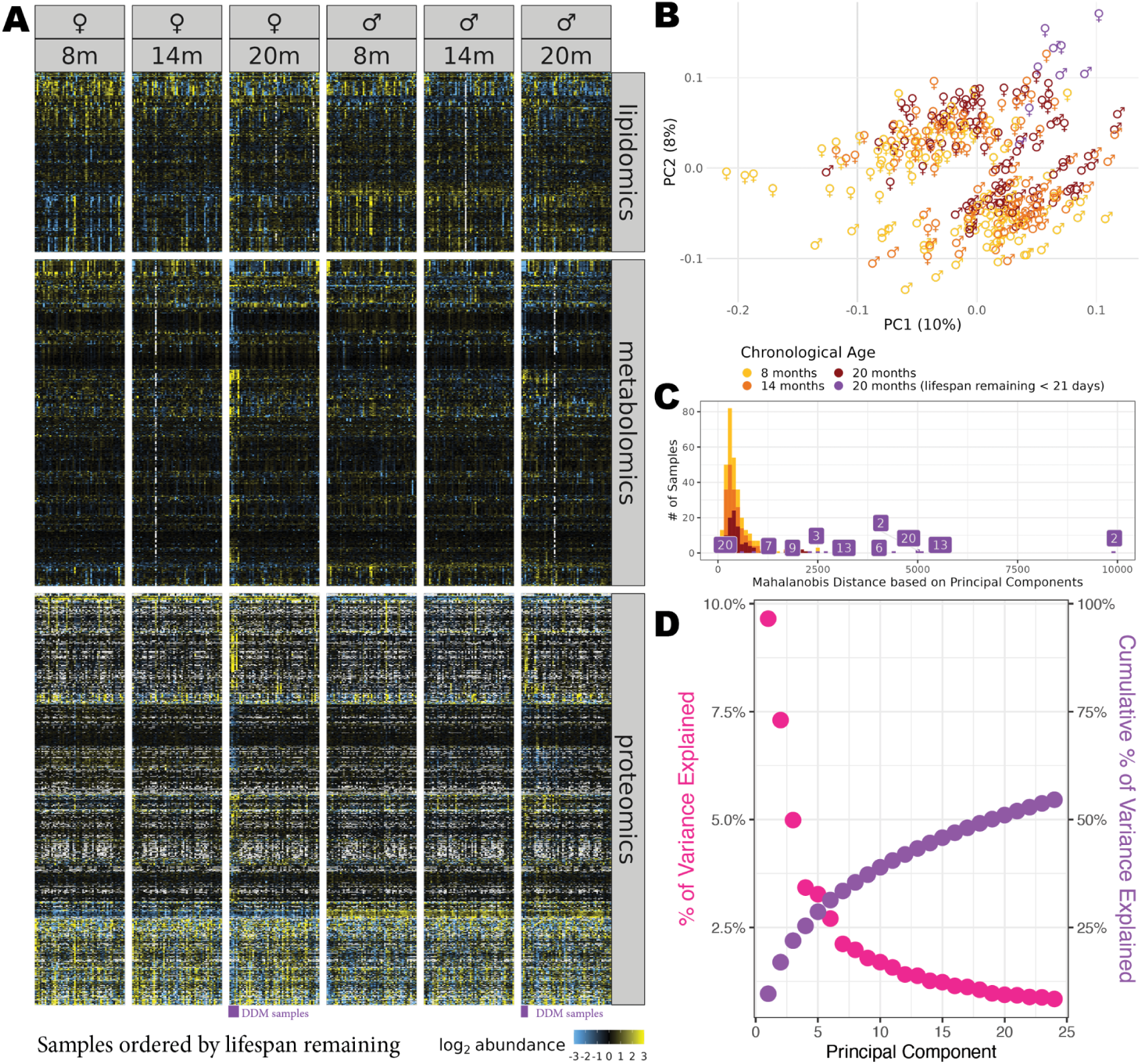
The multiome is shaped by age, sex, and looming death. **(A)** Heatmap of metabolites, lipids, and proteins with samples split by sex and age and ordered by lifespan remaining. 20 month observations with low lifespan remaining died soon after the blood draw (<21 days) resulting in “Death’s Door Metabolite” (DDM) signature. Missing observations are colored white. **(B)** PC plot showing the first two principal components relationship to sex, chronological age, and DDM samples. **(C)** samples’ distances from the centroid of all principal components weighted by their eigenvalues are used to calculate Mahalanobis distance. For DDM samples (purple), the number of days a mouse had left to live is shown. **(D)** Scree plot showing leading principal components contributions to the datasets variability. The slow drop-off in principal components’ % variance explained indicates that a range of factors shape structured variation in this dataset.

In addition to this DDM signature, age and sex are clear drivers of the multiome, with each shaping the structure of principal components 1 and 2 (Figure 2A, B). Beyond these leading principal components, there is a thick tail of trailing principal components whose covariance is likely shaped by other biological effects such as genetics, or lifespan-associated health (Figure 2D). To identify these interesting patterns which may be manifesting as subtle features of the dataset, the major effects highlighted by exploratory data analysis (age, sex, DDM, and the generation batch effects) need to be carefully accounted for.

### Hundreds of genetic loci shape the plasma multiome

The outbred nature of diverse outcross mice can be leveraged as a tool for causal inference if natural variation impacts specific biomolecules with a mediating effect on lifespan or other traits ^25,36^. While the modest number of mice in this study (110) limits our ability to apply such techniques, we did explore whether there are heritable signals and common genetic determinants of molecular traits. Based on similarity of mice’s measurements across three ages we can calculate the broad-sense heritability of each trait, a measure of the fraction of variability derived from additive and non-additive genetic variation ^37^. Heritability varies greatly across molecules but is generally low, reflecting that molecules’ abundances are primarily shaped by strain-independent biological (e.g., age) and technical effects (e.g., measurement noise) (Figure <u>SF9A</u>). There are some highly heritable molecules showing continuous variation in abundance consistent with a polygenic genetic architecture, and others, notably the GM2 and GM3 lipids, showing step-like jumps in abundance consistent with a simpler genetic architecture (Figure <u>SF9B</u>). Changes in GM2/GM3 lipids (due to a loss-of-function mutation in B4galnt1, the enzyme responsible for converting GM3 lipids into GM2 lipids), in NOD mice have been previously reported in the DO mouse lipidome ^38^. Consistent with previous reports, we find that mice that are NOD/NOD at the B4galnt1 (chr10:127Mb) locus accumulate the substrate of this reaction and deplete its products (Figure <u>SF10</u>).

We applied QTL mapping to each trait and cross-trait FDR control and found 258 QTLs associated with molecules’ abundances at a 10% FDR (Figure <u>SF11</u>, Table <u>ST2</u>). We similarly explored whether there was a genetic basis for fold-changes (which could capture a SNP which drives age-dependent accumulation of a molecule) but no associations could be retained at a 10% FDR. This suggests that genetics has a greater impact on shaping the baseline levels of molecules than how they shift with age. Consistent with previous reports, most protein QTLs are local QTLs which map to the protein’s coding locus ^26,27^.

### Identifying the molecular correlates of aging and lifespan

To identify relationships between biomarkers and variables of interest, we can test whether molecules’ concentrations are predicted in a statistically significant fashion based on the variable, accounting for major covariates. This allows us to test each molecule independently, applying standardized statistical tests across all features using a common regression framework. Beyond confirming the specific molecules driving the structure of the leading principal components by sex, chronological age and the DDM (Figure 2B), we can frame regression models to evaluate alternative models of how chronological age and lifespan shape the multiome, which we term “aging archetypes” (Figure 3A, Table 2).

**Figure 3:**
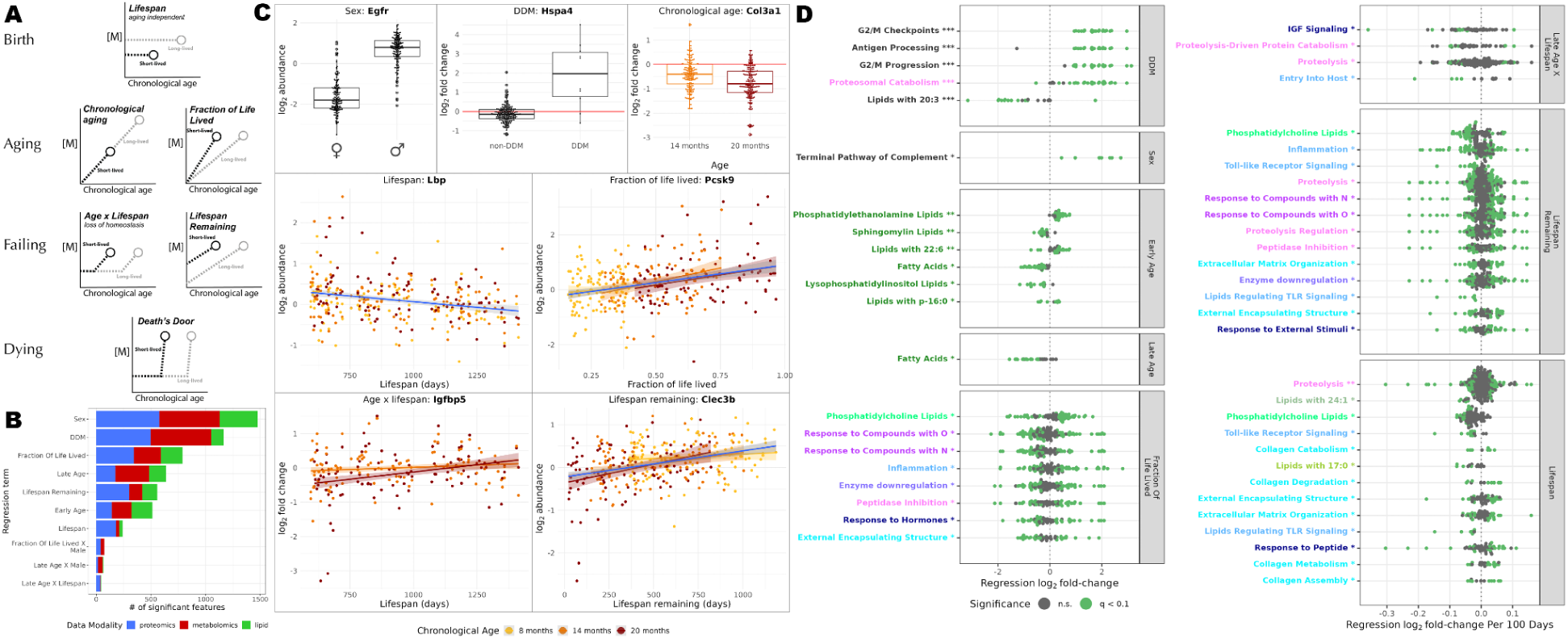
Identifying statistical associations between molecules’ abundances and age or lifespan. **(A)** Cartoon models showing how a molecule’s concentration, [M], would change for a short- and long-lived individual across chronological age if the molecule was an exemplar of each aging archetype. **(B)** P-value histogram of major biological effects with the number of significant associations listed for each term. **(C)** Examples of features which are strongly associated with each of the major effects. **(D)** Beehive plots summarizing gene sets and lipid categories which shift with sex, DDM status, and longevity modes. Individual features within a category are shown as single points summarized by their regression effect size for the coefficient of interest.

**Table 2:** Descriptions of Aging Archetypes: Aging archetypes are tested using one or more regression models. Regression terms labeled with a “$” are used as the primary readout of an archetype while alternative models using abundances and/or treating chronological age as a continuous variable are used for specific analyses.

| Aging Archetype | Definition | Regression Models |
| --- | --- | --- |
| Lifespan<br>*Aging independent* | Date of death - date of birth | $\log_2\text{abund} \sim \text{lifespan}^{\$} + \text{chronological age [categorical]} + \text{sex} + \text{batch}$ |
| Chronological Age | Data of sample collection - date of births | $\log_2\text{fold-change} \sim \text{chronological age [categorical]}^{\$} + \text{batch}$<br>$\log_2\text{abund} \sim \text{chronological age [categorical]} + \text{sex} + \text{batch}$<br>$\log_2\text{abund} \sim \text{lifespan} + \text{chronological age [days]} + \text{sex} + \text{batch}$ |
| Fraction of Life Lived (FLL) | Chronological age / lifespan | $\log_2\text{abund} \sim \text{fraction of life lived}^{\$} + \text{sex} + \text{batch}$ |
| Age x Lifespan<br>*Loss of homeostasis* | Interaction terms between chronological age and lifespan in regression models | $\log_2\text{fold-change} \sim \text{chronological age [categorical]} \times \text{lifespan}^{\$} + \text{batch}$<br>$\log_2\text{abund} \sim \text{chronological age [categorical]} \times \text{lifespan} + \text{sex} + \text{batch}$<br>$\log_2\text{abund} \sim \text{lifespan} + \text{chronological age [days]} \times \text{lifespan} + \text{sex} + \text{batch}$ |
| Lifespan remaining | Lifespan - chronological age | $\log_2\text{abund} \sim \text{lifespan remaining}^{\$} + \text{sex} + \text{batch}$ |

Aging archetypes capture molecules set at birth (i.e., inborn errors detected as aging-independent lifespan associations), those changing with age (chronological aging and FLL) and those defining a failure of homeostasis (age x lifespan interactions and lifespan remaining). We exclude the DDM signature from this list because while it is kinetically similar to age x lifespan interactions, DDM hits could be clearly distinguished from all of the aging archetypes and impacted independent sets of molecules (see below). In contrast, while aging archetypes can be separately tested, they are not independent. Features which vary with any of these archetypes would also be detected, albeit to a weaker extent by a subset of other archetypes (Figure <u>SF12</u>). For example, a molecule that changes with chronological age would also vary somewhat with FLL and lifespan remaining.

When exploring aspects of chronological aging, fold-changes of a mouse with respect to its earliest blood draw are used to identify early age (i.e., between 14 and 8 months) and late age changes (i.e., between 20 and 8 months). Working with fold-changes removes mouse-specific variation to increase power and decrease bias but this solution cannot be generally applied. For example, when mouse-specific attributes are of primary interest (i.e., lifespan or sex) fold-changes would remove the variation of interest. Instead most models are fit using abundances ignoring the fact that mice have repeated measures. So that we can compare multiple aging archetypes fit to identical data, chronological age and age x lifespan models are also fit using abundances (in addition to the primary model using fold-changes). Similarly, we fit models where chronological age was a numeric rather than a categorical variable so that we could estimate the biological age of individual samples based on molecular pathways (see below).

To ensure that we are able to detect biologically feasible effect sizes we can use a power analysis for chronological age-, lifespan-, age x lifespan-dependent changes (Figure <u>SF13</u>). While the level of noise varies feature-to-feature, a typical lipid, our noisiest data modality, possesses a log_2_ standard deviation of ∼0.75 (Figure <u>SF2</u>). At this noise level we have a 90% power to detect chronological age fold-changes of > 0.5, an oldest-youngest lifespan difference of > 0.5 (equivalent to 0.06 per 100 days) and an oldest-youngest late age x lifespan effect of > 0.58 (equivalent to > 0.073 per 100 days). This suggests that we are powered to detect moderately sized changes (well under 2-fold between extremes) even for noisy features.

Having defined statistical models for detecting molecules following each aging archetype, along with other effects such as sex, DDM, and age x sex models (see materials and methods) we fit each model to each of our ∼2,200 molecular features. To combat outlier-driven false positives, we use bootstrapped linear regressions that are robust to departures from residual normality ^39^. Such departures are common in mass spectrometry ^40^ and in cohorts due to non-additive genetic and/or environmental variation ^37^.

As expected from our exploratory data analysis, a large number of features vary by sex and are part of the DDM signature. 1,156 features are associated with at least one archetype (q < 0.1) with FLL, followed by late age (chronological aging), and lifespan remaining topping the list of archetypes that are statistically associated with the most features (Figure 3B, Figure <u>SF14</u>, Tables <u>ST3</u>, <u>ST4</u>). Associations’ frequencies are relatively consistent across data modalities and alternative regression models (Figure <u>SF15</u>, Figure <u>SF16</u>). Examples of molecules that are strongly associated with sex, DDM, and each aging archetype are shown in Figure 3C. Equivalent plots can be generated for any feature of interest in the study using our [R Shiny application](http://public-rstudio-connect.calicolabs.com/doomics_results_browser/).

Molecules changing with chronological age which also exhibit age x lifespan interactions such as Pcsk9, Clec3b, Nid1, and Igfbp5 show a strong anticorrelation of aging and age x lifespan effects (Figure <u>SF17</u>). Molecules elevated with age are associated with a negative age x lifespan interaction where an increase in abundance from early to late age predicts short lifespan. Intuitively, this pattern can be interpreted as age x lifespan interactions reflecting aging patterns which are specific to short-lived animals; or alternatively, markers which are faithfully maintained in long-lived animals. This analysis can be extended to compare all pairs of aging archetypes revealing the correlations among aging archetypes, but also a significant, albeit modest, overlap between features exhibiting aging effects and those with batch effects (Figure <u>SF18</u>). Overlapping signals are also apparent when comparing hits which are significant for just one term or a pair of terms (Figure <u>SF19</u>)

To better understand the biological processes enriched in each process we can apply gene set enrichment analysis to significantly changing proteins and lipid set enrichment analysis to lipids. We chose to not look for metabolite pathway enrichments using a systematic geneset-like approach since cellular pathways are a poor representation of the plasma due to low coverage (i.e., most pathway intermediates are not released into circulation) and undefined regulatory mechanisms.

### Pathway enrichments of aging archetypes

To identify functional enrichments we determined whether each gene set or lipid category was enriched for a regression term’s significant associations (q < 0.1) using Fisher Exact tests. Following FDR control and de-duplication of terms with highly overlapping membership we identified pathways that are significantly altered among each term’s associations (Figure 3D, Table <u>ST5</u>). Functional categories were grouped into related categories (such as those related to proteostasis, or the extracellular matrix). Despite more than ⅔ of features changing with sex, only a single pathway was enriched among sex hits - with complement signaling being elevated in male mice. In contrast, the DDM signature is linked to 35 non-degenerate gene sets, including cell cycle machinery and antigen presentation, that can be best summarized as intracellular components which should not be observed in the plasma unless there is widespread apoptosis or necrosis. This supports the idea that the DDM is capturing the acute and irreversible breakdown of organism-level homeostasis irrespective of the upstream driver of this failure.

Functional associations with aging archetypes are widespread and highly overlapping. For example, phosphatidylcholine (PC) lipids are enriched among associations for lifespan, FLL and lifespan remaining. This is unsurprising because most archetypes are strongly (anti)correlated by definition and thus their associated features overlap (Figures <u>SF12</u>, <u>SF18</u>, <u>SF19</u>). The lack of independence complicates the interpretation of signatures like PC lipids changing. Are they set at birth and independent of chronological age (i.e., associated with lifespan), a clock counting down to a mouse’s death (i.e., lifespan remaining), or something closer to a mouse’s biological age (i.e., FLL)? To determine which aging archetype best describes each aging feature and their higher-level grouping into pathways, we compared archetypes head-to-head.

### Molecular archetypes of aging

Strong associations with one aging archetype necessarily show up as weaker associations with other archetypes resulting in overlapping hits (Figure 4A). To attribute aging changes either at the individual feature level or pathway level to specific archetypes we adopted a model comparison approach. Model comparison methods evaluate a set of plausible models fit to the same data and determine the best supported model balancing how well each model fits the data (its log-likelihood) with the number of degrees of freedom they fit. Here, we performed model comparison using Akaike Information Criteria (AIC) with a correction for finite sample size (AICc)^41^. AIC can be thought of as an extension of the Likelihood Ratio Test to non-nested data treating the relative likelihood of a set of models fit to the same data as exp((AIC_min_ − AIC*_i_*)/2)^42^. This calculation can be extended across multiple features by summing the AICc of individual features for a given archetype (Figure 4B). Using this approach we calculated the relative support for each of the five aging archetypes for each pathway (and an “other” category of proteins, lipids, and metabolites (split into knowns and unknowns)) in the union of all aging archetypes’ functional enrichments (Figure <u>SF20</u>). Each of the five archetypes is associated with at least one functional category. Lipids containing 17:0 are associated with lifespan, while lipids regulating toll-like receptor signaling are associated with lifespan remaining. Both of these associations are borderline improvements over other archetypes for categories with a small number of molecular features so it is unlikely that these are legitimate. In contrast, chronological aging, FLL, and age x lifespan interactions are each associated with a handful of functional categories with proteomic categories primarily varying with FLL or age x lifespan interactions, and metabolomic/lipidomic categories generally shifting with chronological age.

**Figure 4:**
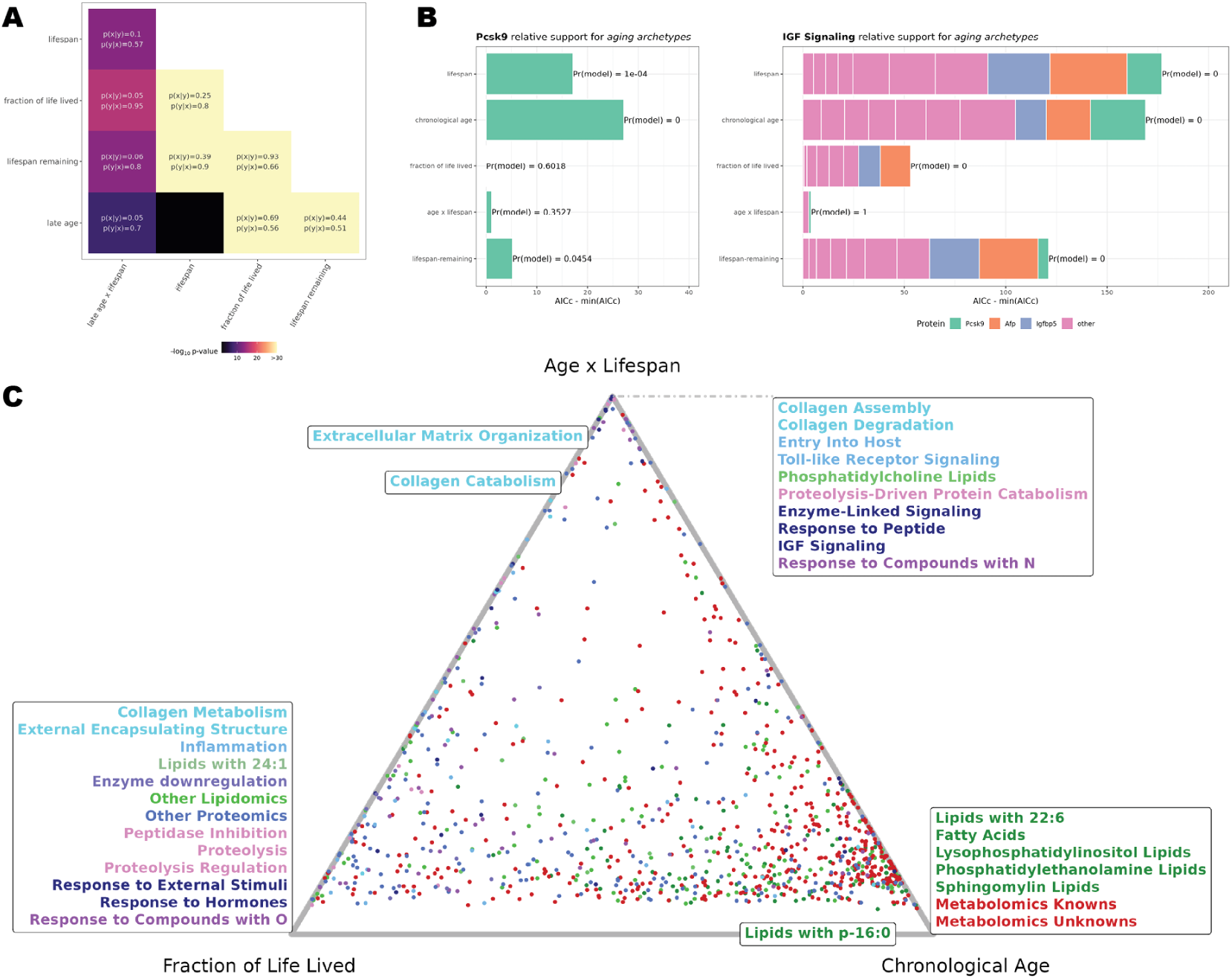
Assigning molecules and pathways to aging archetypes. **(A)** The molecules which are significantly associated with most pairs of aging archetypes strongly overlap. Compared pairs are colored based on the significance of a Fisher Exact test. Cells contain p(x|y) and p(y|x) where p(y|x) would be the fraction of significant molecules in the category on the x-axis which are also significantly associated with the term of the y-axis. p(x|y) is the converse. **(B)** Summarizing individual molecules (using Pcks9 as an example) and pathways using (IGF signaling as an example) based on their relative support across multiple aging archetypes based on AICc. **(C)** Since relative model supports sum to one over the aging archetypes they form a simplex, and three components of the simplex (fraction of life lived, chronological age, and age x lifespan interactions) can be visualized as a Ternary diagram where each points proximity to the three vertices is proportional to its support for that archetype. Both categories (labels) and individual molecules (points) are visualized on this simplex using a color-scheme which maps molecules onto their associated pathway.

The large number of features that are confidently associated with one of the principal archetypes and not others suggests that all three archetypes operate simultaneously to drive variation in distinct subsets of biomolecules. To further explore this idea, we visualized the relative likelihood of these three principle aging archetypes using a Ternary diagram (Figure 4B) where as expected from <u>SF20</u>, functional categories’ archetypes tend to sit at the vertices of the diagram with strong support for a single archetype. This belies the incoherence at the level of individual features - most pathways especially age x lifespan interactions contain a large number of features which are better described by an alternative aging archetype. An example of this is the GO term for collagen assembly where collagens show clear shifts with chronological age (Figure 3C) but the consensus model is an age x lifespan interaction. This is because the age x lifespan interaction model is the only model which fits a subset of features showing loss-of-homeostasis but this model will also fit a feature changing with chronological age well (since chronological aging model is a simpler form of the the age x lifespan model, the main difference in their AICc would be due to the difference in their number of fitted parameters). This aging archetype heterogeneity suggests that aging effects could cascade through individual pathways with some elements reflecting aging stress, and others, aging-associated failure. To better understand the interplay of feature- and pathway-level changes across various archetypes we further explored the relationships between pathways, and between individual features independent of their associated pathways.

### The molecular architecture of variable lifespan

While many measured molecules are associated with aging archetypes (Table <u>ST6</u>), few will causally impact the aging process to the extent that directly modulating a single metabolite, lipid, or protein would be sufficient to extend lifespan. Rather the core drivers of aging may be unmeasured and will often be in cells and tissues that are remote to the plasma (Figure <u>SF21</u>). In such cases, longevity associations will be biomarkers of other aging-relevant processes. Related biomarkers may be similarly affected by a risk resulting in covariation across samples. Molecules may also covary due to common molecular biology upstream of or independent of aging-relevant effects.

In line with the notion that abundances of individual aging-associated molecules may be driven by a myriad of partially overlapping upstream sources of upstream variation, aside from several classes of lipids which are highly correlated, clustering lifespan associations into cliques of correlated molecules is non-trivial. To disentangle pathways, we approach this problem from two complementary perspectives - one approach explores correlations across samples, another identifies correlations across molecules.

To leverage correlations across samples, we can explore whether pathways reflect common or distinct aging patterns. To perform this analysis we took inspiration from Airoldi et al. 2009 where yeast growth rates were predicted based on gene expression results by finding a growth rate which is consistent with a large number of growth-associated genes’ expression ^43^. Analogously, we can estimate the age or delta age of individual samples based on all pathway-associated features where age could be any of the aging archetypes. We approached this by comparing each sample’s residuals (ε) for molecules fit by a linear model of an aging archetype to molecules’ (k ⊂ K) slope with respect to the aging archetype (β_K_):

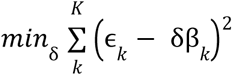

The measure δ can be thought of as delta age because it infers whether a sample’s molecular profile systematically under- or overshoots the direction of its true age (Figure <u>SF22</u>). Using this approach we can estimate each sample’s relative age for a given pathway assuming that the pathway follows FLL kinetics (Figure 5A) or based on each pathway’s best supported archetype (Figure <u>SF23</u>). The hierarchical clustering of these pathways’ relative ages highlights the tight correlations of some aging pathways and looser correlation of others suggesting that the aspects of aging which are most pronounced in individual mice differ. While we removed highly degenerate pathway enrichments, tight correlations may still be driven by overlapping pathway membership; this issue will be addressed below.

**Figure 5:**
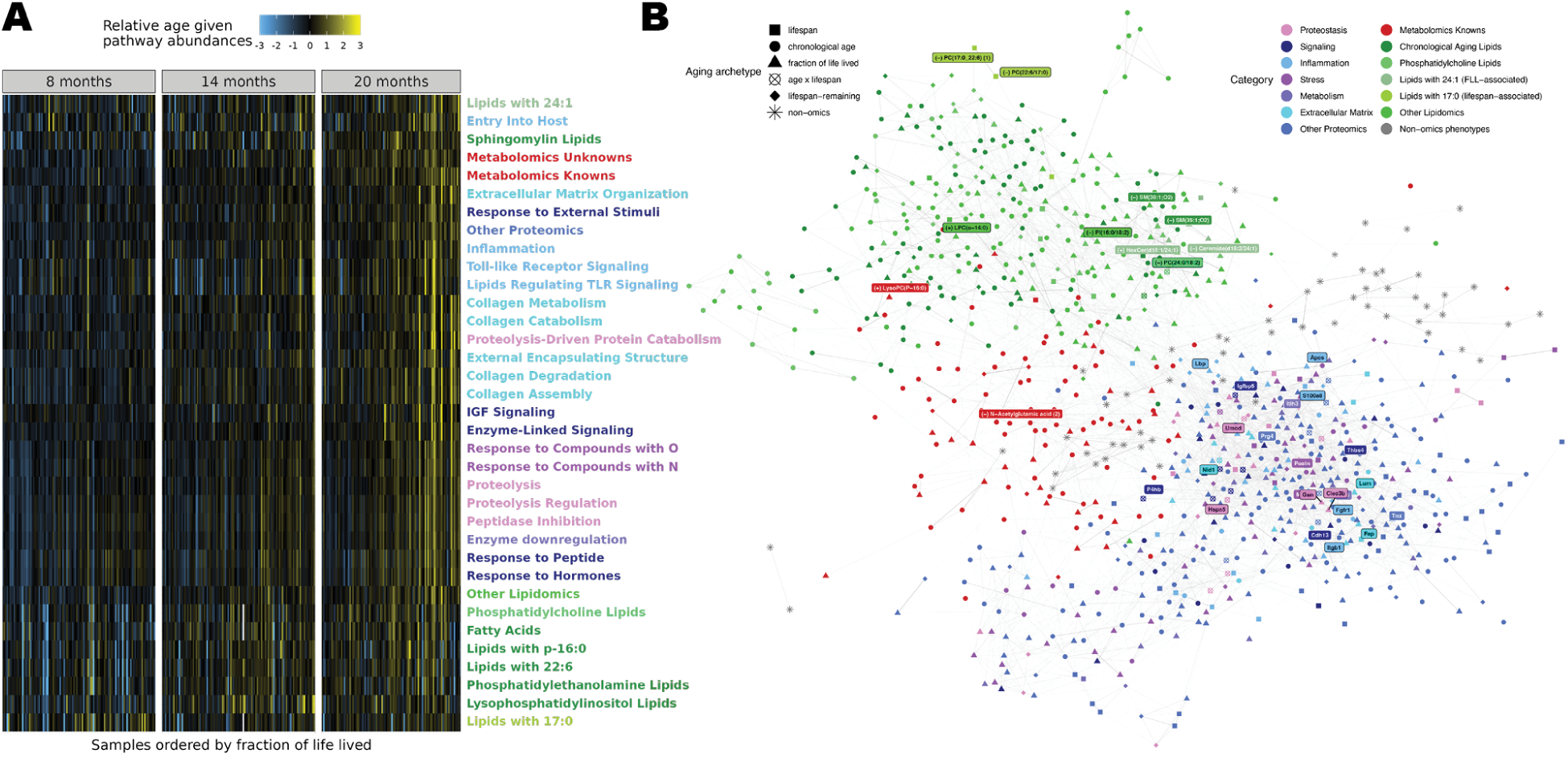
Cross-sample and cross-feature correlations highlight the interconnectedness of distinct functional enrichments. **(A)** Relative age estimates were defined by predicting pathway delta age based on all pathway member’s relationships to FLL, adding this delta age estimate to observed FLL and then standardizing each feature. Samples with a high relative age for a given pathway have an outsized change in the associated pathway relative to peers with a similar FLL. **(B)** Partial correlation network of associations between aging archetype-associated molecular features. Partial correlations were estimated using graphical LASSO and each feature’s strongest partial correlations were preserved to facilitate a layout. Features associated with an aging functional enrichment were colored according to the general category of the association, and strong hits within each of these categories are directly labelled. Features’ shapes were defined by the aging archetype that they best followed based on AICc. Additional non-omics phenotypes were included in this analysis; these includes features such as body weight and CBC measures.

To further understand how pathways covary during aging, while also allowing for the discovery of functional associations which may fall outside of established pathways, we can represent aging-associated molecules’ relationships as a partial correlation network. Partial correlations imply that two molecules are correlated conditioning on all others, thus pairs of molecules will tend to be connected if they privately share sources of variation. In such a network, some nodes may emerge as structuring their neighbors if they are either an accurate biomarker of an upstream health-relevant process or if they are actually mechanistically altering their neighbors (Figure <u>SF21</u>). Because we have more features than samples we cannot directly calculate partial correlations - rather we need to assume that partial correlations are sparse; an appropriate assumption for molecular networks ^44^. To construct this sparse web of partial correlations, we use graphical lasso to estimate a sparse representation of the precision matrix (inverse covariance) which we can rescale to sparse partial correlations ^45^. We find that an optimal model explains ∼70% the off-diagonal covariance in the sample covariance matrix (Figures <u>SF24</u>, <u>SF25</u>) in line with the ∼80% of variance explained by the top 10 principal components of the sample covariance matrix.

The resulting partial correlation network displays a complex architecture (Figures 5B, <u>SF26</u>). Strong partial correlations capture molecules such as fibrinogens which are tightly coregulated (Table <u>ST7</u>). More broadly, weak and strong associations form a single network with the clear separation between lipids and protein associations, with metabolite associations scattered across the graph. The strongest aging associations within pathways (labeled features) are spread across the graph and functional enrichments of the proteome cluster poorly. Indeed when comparing similarity within individual functional categories versus between them based on either correlation or distance on a network, many pairs of gene sets show similar coherence when lumped together versus split apart (Figure <u>SF27</u>). This may be due to the tight interaction between multiple different pathway or, functional categories may be defined with the wrong resolution with categories such as collagen degradation encompassing multiple ways in which age and genetics can shape a set of biomolecules (such as via the loss of collagen, or the deposition of fibrotic plaques). Similarly, there are surely unappreciated interactions between pathways where proteins which are part of the broad reaching signatures of inflammation and proteostatic dysregulation impact other pathways in unappreciated ways.

### Aging dynamics capture how physiology breaks down

The analyses we’ve presented thus far highlight three principles for understanding the molecular nature of aging:

1. Aging manifests in different ways captured by aging archetypes. Some processes seem to change with chronological age while others are better described by FLL. A small number of age x lifespan interactions appear to capture systems which fail in aging mice, while a death’s door signature captures the total systems-level breakdown of mice in their final weeks of life.
2. Aging changes are widespread but functionally coherent. Many pathways with aging signatures fit into the canon of aging biology such as changes in proteostasis and the extracellular matrix. Others are previously unappreciated aspects of the aging process, such as changes in the aging lipidome, and loss-of-homeostasis signatures.
3. While pathways are coherent, in many cases, correlations between pathways are similar to within pathways, making it challenging to disentangle processes from one-another.

Here, we will reconcile these principles into a working model of DO mouse aging which further describes failing subsystems and their potential interactions (Figure 6A). In doing so, we can roughly order events from upstream aging lesions (many of which track with chronological age), to physiological compensatory mechanisms (which often track with FLL), to tipping points where homeostatic mechanisms are insufficient to hold-back aging changes.

**Figure 6:**
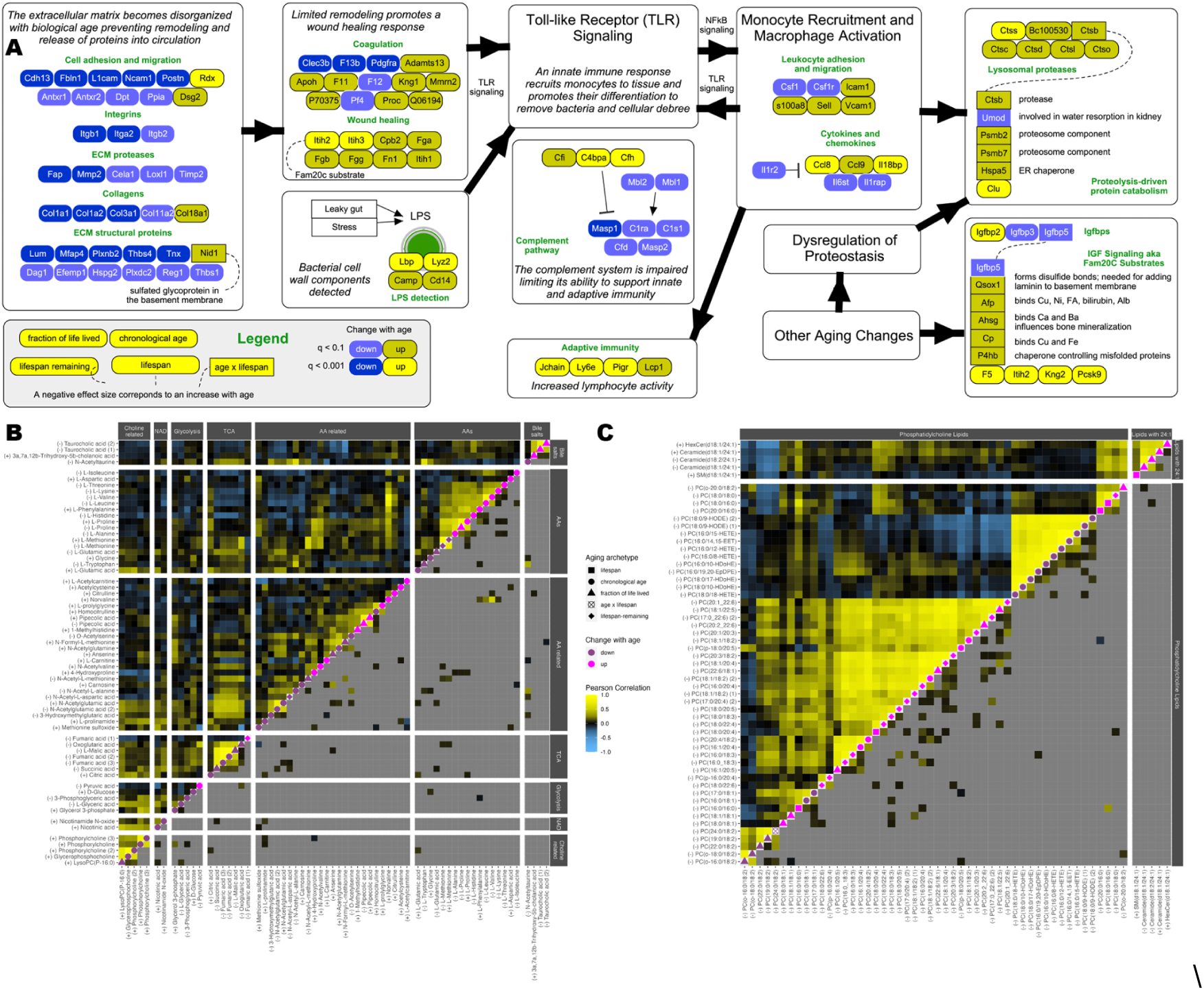
Functional changes in the DO circulating multiome. **(A)**: Cartoon model proposing mechanistic links between aging stresses and innate immunity ultimately leading to the incompletely-understood loss-of-homeostasis signatures. Individual proteins were manually summarized (Table ST8) and summarized with a shape based their aging archetype (pooling all aging archetypes besides age x lifespan) and colored according to whether they increase or decrease with age and the statistical significance of this change. **(B)** Aging patterns and correlations in the aging metabolome are visualized using a dynamics (p)corr plot. In this plot upper diagonal entries are colored by metabolites’ Pearson correlation, the diagonal indicates whether a metabolite increases or decreases with age, and lower diagonal entries include partial correlations between pairs of metabolites. **(C)** Dynamics (p)corr plot highlighting patterns in lipid categories which shift with FLL.

As there are 1,224 features significantly associated with one or more aging archetypes (Table <u>ST6</u>), we simplified the interpretation of aging mechanisms by focusing on known metabolites, and lipids and proteins which map onto aging-enriched functional categories, as well as including the top 20 other proteins with the strongest association for their aging archetype. When exploring the metabolome and lipidome we try to distinguish features associated with chronological age versus FLL. For the proteome, we generally interpret features which are best described by chronological age, lifespan or lifespan remaining aging archetype as actually tracking with FLL since we lack the power to reliably discriminate among these cases for individual features (Table <u>ST8</u>).

Both the metabolome and lipidome are strongly shaped by chronological age. Among the diverse changes in the aging metabolome, we observe a drop of nicotinamide with age, as well as in many metabolites of the TCA cycle consistent with reported drops in NAD and mitochondrial function with age (Figure 6B, Figure <u>SF27</u>) ^4,46^. Conversely, we observe age-associated increases in many amino acids, and taurocholic acid, a bile salt where taurine is conjugated to cholic acid, which is one of a small number of metabolites which appears to scale with FLL rather than chronological age (Figure <u>SF28</u>). Elevation of taurine-conjugated bile acids was previously observed in 15 month old rats, and the supplementation of taurine has recently been reported to extend lifespan ^47,48^.

Like the metabolome, the lipidome primarily covaries with chronological age, with notable exceptions. Lipids with phosphatidylethanolamine headgroups increase with chronological age, while free fatty acids, sphingomyelin lipids, and lipids with phosphatidylinositol headgroups decrease with chronological age (Figure <u>SF29</u>). 17:0, p-16:0 and 22:6 tail groups were each functionally enriched among chronological aging changes, but lipids with these tail groups tend to be tightly correlated with other examples of their headgroup, so changes in the composition of fatty acid changes in the aging lipidome are likely minor relative to shifts in the distribution of lipid classes. While most lipids change with chronological age, two categories of lipids, those with phosphatidylcholine (PC) headgroups or 24:1 fatty chains, followed the FLL aging archetype (Figure 6C). Most PC lipids increase with FLL, while a subset containing oxidized fatty acid chains (e.g., hydroxy-eicosatetraenoic acid (HETE)) decrease with age ^49^. Another clique of lipids were defined by their 24:1 fatty acids. These lipids are further unified by the fact that they all contain a second d18:1 or d18:2 chain and are either ceramides, hexosylceramides (HexCer) or sphingomyelins (SM). While HexCer and SM lipids are derived from ceramides lending a structural coherence to the overall group, it is unclear why these lipids shift with FLL in such a coherent manner.

While few studies have profiled the circulating aging metabolome and lipidome, many proteomic changes can be interpreted as canonical hallmarks of aging ^4,28–30,50^. Among them, chronic increases in inflammation, i.e., inflammaging, and alterations of proteostasis are clear features of our dataset ^28^. In addition, changes in extracellular-matrix associated proteins, i.e, the matrisome, which result in a more highly crosslinked, irregular, and protease-resistant matrix are appreciated as a key aspect of the aging process which may contribute to stem cell exhaustion, and recruitment of immune cells ^50–52^.

Broad decreases in circulating levels of extracellular matrix (ECM) components include fibril-associated collagens (type I, III, and XI), integrins, and additional ECM-associated structural proteins, proteases, and proteins mediating cell-cell communication and migration (Figure <u>SF30</u>) ^52,53^. This is consistent with an increasingly protease-resistant ECM where turnover of components is low and hence fewer proteins are liberated into circulation. This depletion of ECM-derived proteins in plasma contrasts with an elevation of proteins involved in wound healing and tissue remodeling which are deposited on the ECM, but generated in remote sites, particularly the liver ^54^. These proteins include the hyaluronic acid carrier proteins Itih1 and Itih3, fibrinogens, and numerous proteins involved in coagulation. Beyond their direct role in serving as the primary substrate for clotting, fibrinogens are sensed by Toll-like receptor (TLR) signaling pathways to promote tissue remodeling ^55^.

As one of the two main arms of activation of the innate immune system, TLR signaling is highly responsive to bacterial exposure, with the exposure to gram-negative bacteria being tracked primarily through the detection of the lipopolysaccharide (LPS) by LPS binding proteins (Lbp) and other secondary proteins ^55^. These signals of LPS exposure increase precipitously with age, likely due to either age-associated increases in gut leakiness or systemic stress (Figure <u>SF31</u>) ^56,57^. The dual stimulation of TLR signaling through fibrinogen and Lbp may activate the pathway, stimulating release of cytokines and type I interferon in order to recruit monocytes and promote their differentiation into tissue-resident macrophages ^55^. Lbp is often used as a biomarker of an LPS-driven innate immune response ^58^ and consistent with monocyte recruitment and macrophage activation through TLR signaling, Lbp is strongly correlated with both monocyte percentage and CD11b+ percentage read-out through FACS (Figure <u>SF32</u>). Downstream of TLR signaling, secreted cytokines will activate lysosomal cathepsins in macrophages and in other cell types, and we note that these lysosomal proteases are markedly elevated with age ^30^. Curiously, while innate immunity seems to increase with age through TLR signaling, the other major pathway driving innate immune responses, the complement pathway, appears to decrease its activity with age - pathway inhibitors (Cfi and Cfh) increase with age, while components of the classical pathway (e.g., C1, Masps) are depleted ^59^.

Decreased levels of ECM-associated proteases in circulation and elevated levels of cathepsins downstream of TLR signaling are specific instances of broad changes in proteostasis with age. In line with the collapse of proteolysis with aging through impairment of both lysosomal and proteosomal proteolysis ^4,30,29^ we observe a clear directional shift in proteins which are part of the “protease-mediated protein catabolism” geneset which includes both proteosomal and lysosomal proteins (<u>Figure SF33</u>). However, rather than being a progressive change with FLL, these proteins are stably maintained across age before spiking as a loss-of-homeostasis, age x lifespan interaction, signature. The robust increase in intracellular proteostasis proteins in plasma contrasts with extended patterns of directionally incoherent changes in both proteases and protease inhibitors, many of which operate in circulation.

The pronounced elevation of intracellular proteolysis components in plasma before the broader onset of the catastrophic DDM suggests that this signature may be a readout which is mechanistically proximal to the onset of total systems failure. Like this change in proteolysis, elevation of the “IGF signaling” pathway from Reactome yielded a clear age x lifespan signature. In this dataset, the pathway’s name is a misnomer, since aside from circulating IGF binding proteins (Igfbps), most members of this gene set are included because they are substrates for the Golgi-associated kinase Fam20c which is responsible for secreting plasma phosphoproteins ^60^. This raises the possibility that Fam20c activity may shift with age resulting in functional consequences - while Fam20c levels do not change in the plasma with age, its localization is modulated by Fam20a and hence its activity as a Golgi kinase may be poorly represented by its circulating concentration ^61^. Fam20c ligands are involved in diverse processes particularly proteostasis, Igf1 signaling and ion homeostasis and consequently dysregulation of Fam20c activity could broadly alter homeostasis (<u>Figure SF34</u>). As a ligand of Fam20c, Igfbp5 is a strong age x lifespan interaction, where it exhibits a pronounced drop in pre-DDM low lifespan remaining mice. Igfbp5 is correlated with circulating Igf1 levels and other Igfbps which change progressively with age. Another target of Fam20c which exhibits a pronounced age x lifespan interaction is Pcsk9, a key regulator of cholesterol homeostasis, which operates by binding to low-density lipid receptor family member proteins and promoting their intracellular breakdown ^62^. Severe mutations of Pcsk9 lead to familial hypercholesterolemia, while the protein is a drug targeted more generally due to its association with coronary artery disease and cardiovascular disease ^63^. Despite the established connection between Pcsk9 and cholesterol we find that Pcsk9 is uncorrelated with the six cholesterol esters in our dataset suggesting that functional variation in Pcsk9 levels likely does not cascade into organismal decline.

These changes are but a subset of wide changes in homeostasis and mouse-level physiology which are reflected in circulation. Other changes included alterations in lipid transport and deposition, growth factors, and oxygenation (<u>Figure SF34</u>). Many changes with age can be reconciled into a working model which describes the relationship between the aging ECM, innate immunity and proteostasis and highlights proteolysis and Fam20c as possible points of failure (Figure 6A). But this model remains incomplete, as there are numerous mechanistic connections which still need to be made to better understand how the thousands of detected changes in the aging metabolome, lipidome and proteome fit into a common framework of reporters and drivers of aging failures.

## Discussion

Here we release a high-quality multiomic dataset and an interactive [R Shiny Browser](http://public-rstudio-connect.calicolabs.com/doomics_results_browser/) which allows any researcher to visualize thousands of molecules’ associations with age in plasma. In carrying out this study we encountered a range of experimental and technical challenges which complicated our analysis and subsequent work will be needed to fully articulate the molecular causality of aging.

One challenge of creating a longitudinal cohort with multiple measurements per mouse is that mice’s age were confounded with draw date. To the extent that non-biological effects were not adequately controlled over time, e.g., switching reagents used in plasma preparation, experimental changes would be confounded with aging changes. By enrolling five staggered cohorts of mice we identified and corrected two such effects (the LBD and early G8 effects). Both effects primarily alter the metabolome and lipidome and features affected by these batch effects show some overlap with chronological aging associations. Still, with these effects regressed out, we see consistent changes with chronological age throughout the metabolome and lipidome. To make it easy to determine whether a batch effect may be contaminating a biological finding, any biological features’ changes can be stratified by biological or technical batch in our Shiny app.

A second challenge arose from generating mass spectrometry data in multiple tranches separated by around two years. From an informatics point-of-view, it was difficult to align metabolomics and lipidomics features across the full dataset. In the end, it was possible to align the metabolomics data by developing new alignment algorithms, but due to the non-monotonic nature of retention time shifts in lipidomics data, feature-level alignment could not be accomplished. This precluded the systematic analysis of lipidomic unknowns, with a few exceptions.

Beyond these limitations in accurately detecting a wide range of molecules’ abundances across our aging cohort, elements of our experimental design hampered data interpretation. The late blood draws of short-lived mice tend to be similar to the early blood draws of long-lived mice for features following a FLL trend, suggesting that the main differences between long- and short-lived mice is the rate that they are transitioning along a common aging cascade, but the absence of late blood draws for long-lived animals prevents us from concluding that this is the case. We expect that signatures of FLL and aging would blend somewhat at very large ages (when most mice have died) as chronological age approaches the limit of mouse lifespan and old-age becomes synonymous with an FLL of near one. The late life chronological changes may reflect either new changes, or an acceleration of aging-associated changes (as we have seen with the greater magnitude of late age changes than early age). It is also unclear whether new failure points (i.e., the loss-of-homeostasis age x lifespan interactions and subsequent DDM) manifest in long-lived mice, or if these mice just take longer to reach breaking points which are conserved across DO mice. It is possible that mechanisms of age-associated mortality between short- and long-lived individuals differ, as is the case in the human population ^64^. Short lived strains have high occurrences of cancer, while the occurrence is somewhat lower in long-lived strains (but still far more common than in humans) ^21–23^. While a mouse’s cause of death was not assessed in our study, most mice, particularly short-lived mice, die of cancer and hence the FLL, loss-of-homeostasis and DDM signatures are all likely either preludes to or concomitant features of cancer. FLL-associated chronic inflammation processes could drive oncogenesis ^65^, while the collapse of immunosurveillance and decreased competition for growth of an increasingly unhealthy pool of non-cancerous cells could allow for the rapid tumor growth ^66^, though there is likely already a high tumor burden in mice within the DDM window. It is not clear how proteasomal collapse, nor dysregulation of the circulating phosphoproteome could promote oncogenesis but this connection warrants further investigation.

The identification of discrete subsets of molecules following three distinct aging archetypes provides a useful framework for organizing molecules with distinct relationships to aging and lifespan, but also highlights how easy it is to confuse aging archetypes with one another. Indeed, within this study we lacked the power to discriminate among correlated aging archetypes for individual features, while we could resolve archetypes by aggregating results to the level of pathways.

Studies exploring shared patterns of aging across outbred cohorts have typically done so by profiling individuals with a shared chronological age ^6,7^. In model organisms this is often because terminal profiling of tissue physiology precludes the measurement of natural lifespan, while in humans, aging is often explored through cross-sectional studies avoiding more challenging longitudinal analyses requiring long-term follow-up ^12^. By studying cohorts of DO mice, genetic variability creates underlying differences in rates of aging, possibly partly through Igf1 signaling ^21^, while non-terminal phenotyping and measurement of lifespans allows us to decouple aging from lifespan. Ignoring these underlying differences in rates of aging, as we could have done by only focusing on chronological age, would lead to misinterpretation of aging dynamics leading to an over-appreciation of molecules associated with chronological age at the expense of molecules scaling with biological age. At the same time, associations with lifespan are defined by canonical changes in the aging proteome, due to the confounding anticorrelation of lifespan and FLL. Future studies using outbred individuals should take care in interpreting associations with chronological age or lifespan, as either measure may mis-represent underlying associations with biological age.

Most hallmarks of aging were defined in inbred strains where due to low animal-to-animal variability, biological age and chronological age were aligned ^67,68^. Still even in this context, there is a keen appreciation for the interplay between chronological, biological age and lifespan. In *C. elegans*, only a subset of the age-associated changes in the transcriptome and proteomes are slowed by lifespan-extending partial loss of function *daf-2* mutations indicating that different molecules track with biological and chronological age ^69,70^. Long-lived strains of models organisms, such as the *daf-2* strains of *C. elegans* or yeast strains with a low rDNA burden, display an extended healthspan and hence be thought of as having a lower biological age ^71,72^. This notion that we can capture summaries of the aging process and then profile individuals based on these markers to define their “biological age” and delta age (i.e., the difference between biological and chronological age) is central to how the longevity field is exploring interventions which can slow the aging process ^73^.

Following this strategy we estimated sample-specific delta ages based on individual pathways (Figure 5A) and while biological age estimates based on these delta ages are correlated across samples, individuals who would be considered “old” differ particularly when comparing delta age for pathways that follow a chronological aging versus a FLL aging archetype. This underscores that using chronological age alone to estimate biological age and delta age may be undesirable. If we were to select molecules that consistently predict chronological age then our “aging clock” would focus on metabolites and lipids which accurately track the increased exposure of physiology to the insults of damage, but do not reflect intertwined changes in inflammaging, ECM dysregulation, and proteolysis which are central to our understanding of biological aging. Some aging clocks such as PhenoAge and GrimAge focus on markers which change with age, and are associated with functional outcomes - we believe that such clocks will produce meaningful estimates of biological age, such that changes in age following interventions will be borne-out in commensurate changes in health and mortality ^74,75^.

While FLL is a valid measure of biological age because it approximates the relative risk of mortality of individuals in a population, it is flawed for the same reasons that limit the utility of directly studying variable lifespan in experiments ^12^. Namely, death is a discrete stochastic event and a healthy individual could die an “untimely” death which may be unrepresentative of the all-cause mortality risk implied by their biological age. In our data this may manifest as mice with non-fatal disorders which nevertheless require euthanasia such as dermatitis. Furthermore, any measure which requires measuring lifespan is of little utility when exploring aging through terminal experiments, or those where an intervention is applied. The estimates of sample-level FLL based on pathway information which we described (Figure 5A) could provide an instantaneous estimate of FLL avoiding these limitations.

Measures like this “FLL-pathway age” which define the state of individual aging components could dovetail with other functional summaries of intermediate phenotypes to describe the progression of individuals along distinct aging trajectories. We describe one such set of mechanistic transitions involving interlocking changes in the proteome transitioning with FLL, and ultimately culminating in two near death loss-of-homeostasis signatures. This signature is complemented by previously unappreciated alterations of the aging metabolome and lipidome reflecting both chronological aging dependent physiological insults, and key molecular classes which become entrained to variable rates of biological aging. One class of FLL-associated molecules are PC lipids which are the most abundant lipids class and are overrepresented in lipid droplets and many organelle’s membranes ^76^. While past reports have focused on a depletion of PC lipids leading to membrane stress in disease, we observe an accumulation of PC lipids with FLL ^77^. This could contribute to lipid stress, but it may also be a read-out of the increase in organellar membrane expected from the increased surface area to volume of fragmented organelles ^78^.

The most prominent proteomic features of our FLL signature are concerted changes in the extracellular matrix, innate immunity, and proteostasis. ECM-associated proteins deposited from nearby cells are depleted in circulation suggesting decreased remodeling of the ECM with age ^52^. In contrast, systemic factors deposited from remote sites to promote ECM remodeling and wound healing are enriched. Curiously, unlike much of the proteome, many collagens seem to decrease with chronological age rather than FLL in circulation suggesting that changes in the ECM may be a stress, which snowball into the physiological strain of FLL. These observations are consistent with changes in ECM cross-linking which in turn alter matrix stiffness contributing to both stem cell exhaustion and chronic inflammation ^52,79^. Inflammation is further driven by increased sensing of LPS where both wound-healing-promoting fibrinogens, and Lbp activate TLR signaling resulting in increased monocyte recruitment and macrophage polarization ^4^. TLR signaling also serves as an input into NF𝜅B signaling leading to secretion of pro-inflammatory cytokines and an inflammaging response. Inflammation in-turn promotes tissue remodeling, a strain of proteostasis which would be compounded by accumulation of mis-folded proteins ^80^.

The spike in proteosomal proteins in circulation suggests that there is an acute crisis in a subset of cell types which precedes wider cellular death as part of the DDM. The death of cells with high proteosomal factors could result from cells being overwhelmed by protein aggregates which interfere with proteosomal function ^81^. Alternatively, continued activation of the proteasome may induce apoptosis through neutralization of IAP (inhibitor of apoptosis) in a stressed cell population ^82^. While this mechanism may guard against sporadic failures of individual cells who can “pass the baton” to healthy cells, it could cascade into organism failure when a population of cells makes a concerted commitment to apoptosis.

A similar crisis seems to be reached when cells acutely accumulate a diverse set of proteins with the common feature that each is a target of the Golgi kinase Fam20c which is responsible for phosphorylating secreted proteins particularly those involved in biomineralization ^60^. Loss-of-function mutations in Fam20c lead to Raine syndrome, an autosomal recessive rare disease characterized by osteosclerosis and brain calcifications leading to microcephaly ^83^.

Understanding the causal role of these loss-of-homeostasis signatures in the aging process remains an open problem but one where future efforts can hope to disentangle the true drivers of aging’s etiology. Four strategies stand-out as promising approaches to clearly resolve the causal drivers of aging:

1. With more profiled individuals, correlations across individuals may emerge as a set of functional elements of the aging process and it is likely that these modules overlap with but are poorly represented by the types of gene sets we explored to define the complex biology at play. Modules can be further anchored using either directly measured or latent intermediate phenotypes ^12^. At this scale, hazard modeling will be possible where modules define separate signatures funneling into baseline and/or age-associated risk ^2^.
2. Causal assertions about how variation across individuals impacts health modules and in-turn hazard could be tested either by lifespan-extending perturbations or using natural variation. Lifespan-extending perturbations such as dietary restriction or molecule supplementation read-out through molecules with aging x lifespan interactions may decelerate aging’s impact downstream of the lifespan-extending intervention’s core effects ^73,84^.
3. Taking advantage of the bounded genetic heterogeneity of the DO cohort could provide further insights into whether mendelian perturbations affect individual molecular components in a way that tracks with downstream causal consequences ^25,36^. Here, we demonstrated that genetic associations can be identified and that molecules are heritable but 110 mice limits our ability to identify molecules’ genetic determinants and to in-turn use this information for causal inference.
4. One of the greatest limitations of studying plasma is also one of its greatest strengths - molecules from plasma could come from anywhere in the body. Without a reference of where individual molecules come from, nor where they go, it is difficult to anchor interpretation in physiology. With the emergence of high quality genome-scale datasets studying aging DO mice in many major tissues, this is beginning to change ^6,25–27^. Connecting our understanding of tissues and their connection in plasma to gross physiology is a problem suited for the nascent field of systems physiology ^9,85^.

Understanding how aging’s diverse changes lead to systems level failure in outbred mice through the combination of genetic exposures, molecular mediators, physiologically-relevant intermediate phenotypes, and measures or surrogates of longevity creates a blueprint for implementing similar strategies in humans. To the extent that aging’s etiology is conserved between mouse and humans we can learn more - directly mapping aging stresses (i.e., changes with chronological age), strains (i.e., processes scaling with FLL), and failures (i.e., processes associated with age x lifespan interactions) onto human physiology. The use of diverse mice exposes a greater diversity of aging trajectories than would be present in any inbred strain. This necessarily increases the variance of any aging pattern but also reduces the bias that we would over-focus on an element of aging biology which is not broadly applicable within, yet alone beyond, mice ^20^. Overlaps between human and murine aging mechanisms are likely common, as many of the processes we discussed in depth such as changes in ECM, inflammation, and proteostasis are appreciated as conserved aspects of the aging process conserved across mammals, while some of the unappreciated aspects of aging physiology which we highlight particularly those reflected in the metabolome and lipidome will need to be corroborated in humans.

## Supplemental Tables

Table ST1 – Abundances

.xlsx summary of molecule abundances summarized as features, samples, and measurements tables.

Table ST2 – QTLs

Summaries of 258 molecular QTLs. Column descriptions: A - A - feature_name: a unique name for the feature (see ST1). B - genename: gene symbol if “feature_name” is a protein. C - gene_id: entrez gene ID if “feature_name” is a protein. D - chr: chromosome of QTL. E - pos: locus maximizing the LOD score. F - lod: LOD score for association to locus. G/H - ci_lo/ci_hi: confidence interval for QTL. I - start_location: location of “gene_id” in base pairs and signed to represent strand. J - Chromosome - chromosome where “gene_id” is encoded. K - Pos_Mb - absolute value of “start_location” * 1e6. L - strand - strand encoding ORF {-1,1}. M - qtl_type - summary of whether QTL is a local QTL where the gene’s locus matches the QTL’s locus, a trans QTL where it does not, or other types of pQTL or mQTLs where such an assessment was not made.

Table ST3 - Differential Abundances

Differential abundance results for primary terms across all features. Column descriptions: A - data_modality: proteomics, metabolomics, lipidomics. B - data_type: finer-grain summary of “data_modality” differentiating the mode of metabolomics and lipidomics features. C - feature_name: a unique name for the feature (see ST1). D - model_name: name of regression model that was fit. E - term: regression term. F - estimate: effects size of “term”. G - statistic: t-statistic for term’s prediction. H - pvalue_ols: ordinary least squares p-value based on “statistic”. I - pvalue_bs: bootstrapped p-value for term’s regression. J - q.value: “pvalue_bs” corrected for multiple testing.

Table ST4 - Differential Abundances (Extended)

Differential abundance results for alternative regression models across all features. Column descriptions: A - data_modality: proteomics, metabolomics, lipidomics. B - data_type: finer-grain summary of “data_modality” differentiating the mode of metabolomics and lipidomics features. C - feature_name: a unique name for the feature (see ST1). D - model_name: name of regression model that was fit. E - term: regression term. F - estimate: effects size of “term”. G - statistic: t-statistic for term’s prediction. H - pvalue_ols: ordinary least squares p-value based on “statistic”. I - pvalue_bs: bootstrapped p-value for term’s regression. J - q.value: “pvalue_bs” corrected for multiple testing.

Table ST5 - Functional Enrichments

.xlsx summary of pathways which are enriched among differential abundance associations summarized as summaries of distinct pathways, enrichments of pathways for specific terms, and members of molecules associated with each pathway/term.

Table ST6 - Aging Features

1,224 features which are statistically associated with one or more aging archetypes. Column descriptions: A - data_modality: proteomics, metabolomics, lipidomics. B - feature_label: a molecule’s name or protein symbol which may not be unique. C - feature_name: a unique name for the feature (see ST1). D - category_label: a readable name for a functional enrichment (or other {data_modality} for a molecule which changes with age but didn’t fall into a functionally enriched pathway). These names and term associations are defined in ST5. For molecules that are members of multiple enriched pathways; the pathway with the strongest statistical association is selected. E - category_general_label: “category_label”s high-level category defined in ST5. F - feature_aging_archetype: archetype minimizing the feature’s AICc; G - change_w_age: does the feature increase or decrease with age based on the regression effect size. The sign of effects are flipped for lifespan, age x lifespan, and lifespan remaining. H - term: the regression term for the archetype in the best-fitting model. I - estimate: regression effect size. J - pvalue_ols: ordinary least squares p-value from regression. K - pvalue_bs: bootstrapped p-value from regression. L - q.value: fdr-controlled pvalue_bs; lipidomics unknowns do not possess a q-value because they were selected due to strong nominal significance. M - go_aging_archetype: aging archetype of “category_label” pathway. N - gsea_term: specific regression model which minimized the patwhay’s AICc leading to selection of “go_aging_archetype”. O - qvalue_aging_archetype: q-value of “go_aging_archetype” hits enriched in “category_label” pathway. This value can be an NA if either a feature is part of a other {data_modality} pathway or if a pathway is assigned to a gsea_term based on AICc even though it was not significantly enriched based on the Fisher Exact test. P - qvalue_min_overal - minimum q-value for “category_label” being associated with any aging term.

Table ST7 - Partial Correlations

5,040 partial correlations selected through graphical LASSO. This table includes three variables: feature_1, feature_2, and partial_corr. Since partial correlations are symmetric only one combination of a pair of features is included.

Table ST8 - Biological curation of the aging proteome

233 ing-associated proteins whose functions were manually curated to identify additional aging patterns. Proteins from ST6 were selected if their value for “category_general_label” was “Other Proteomics” or if they were one of the top-20 “Other Proteomics” associated based on “pvalue_bs”. Column descriptions: A - feature_name: a unique name for the feature (see ST1). B - feature_label: a molecule’s name or protein symbol which may not be unique. C - category_general_label: see ST5. D - manual_category: manually curated category of protein. E - manual_general_category: high-level grouping of “manual_category”. F-H - feature_aging_archetype/change_w_age/q.value: corresponding to values in ST6. I - Notes from curating protein.

## Supplemental Figures

**Figure SF1.**
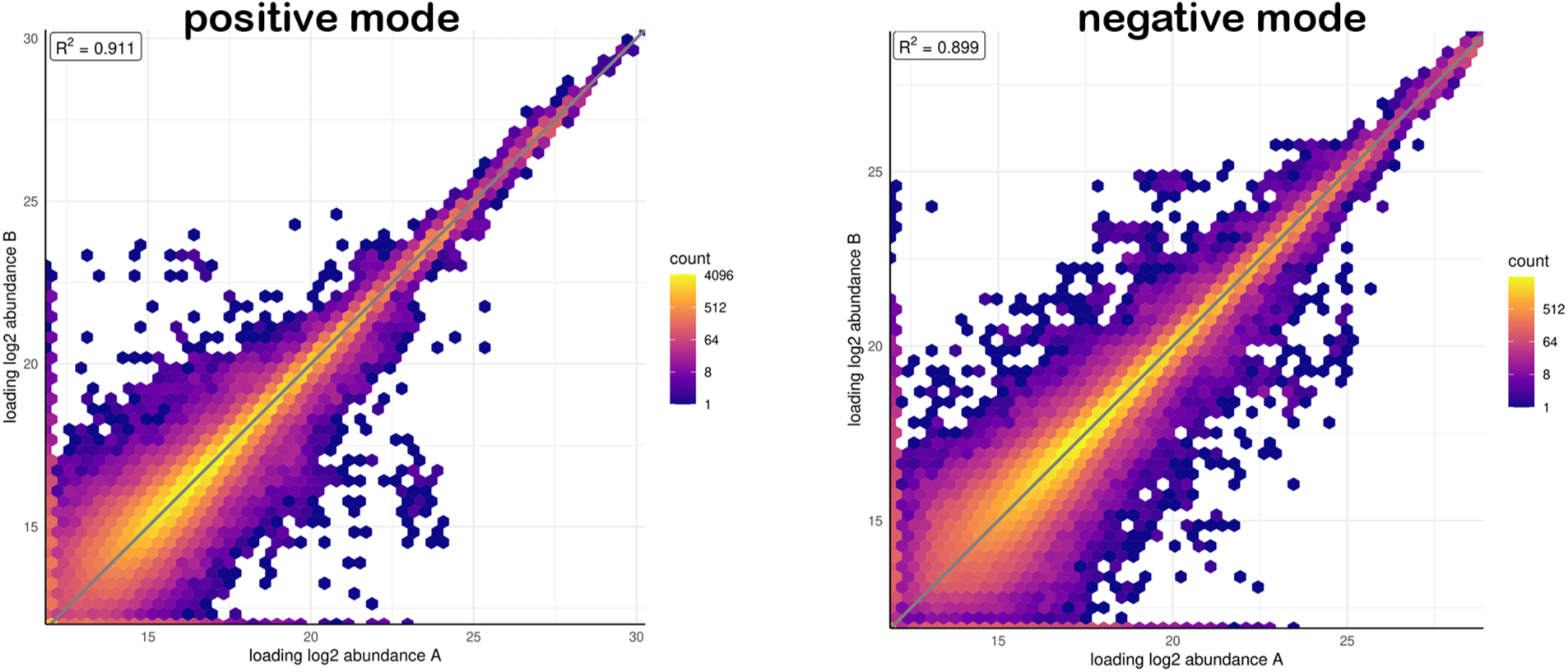
Technical replicates where the same sample was injected into the mass spectrometer in duplicate are shown for set1/2 metabolomics.

**Figure SF2.**
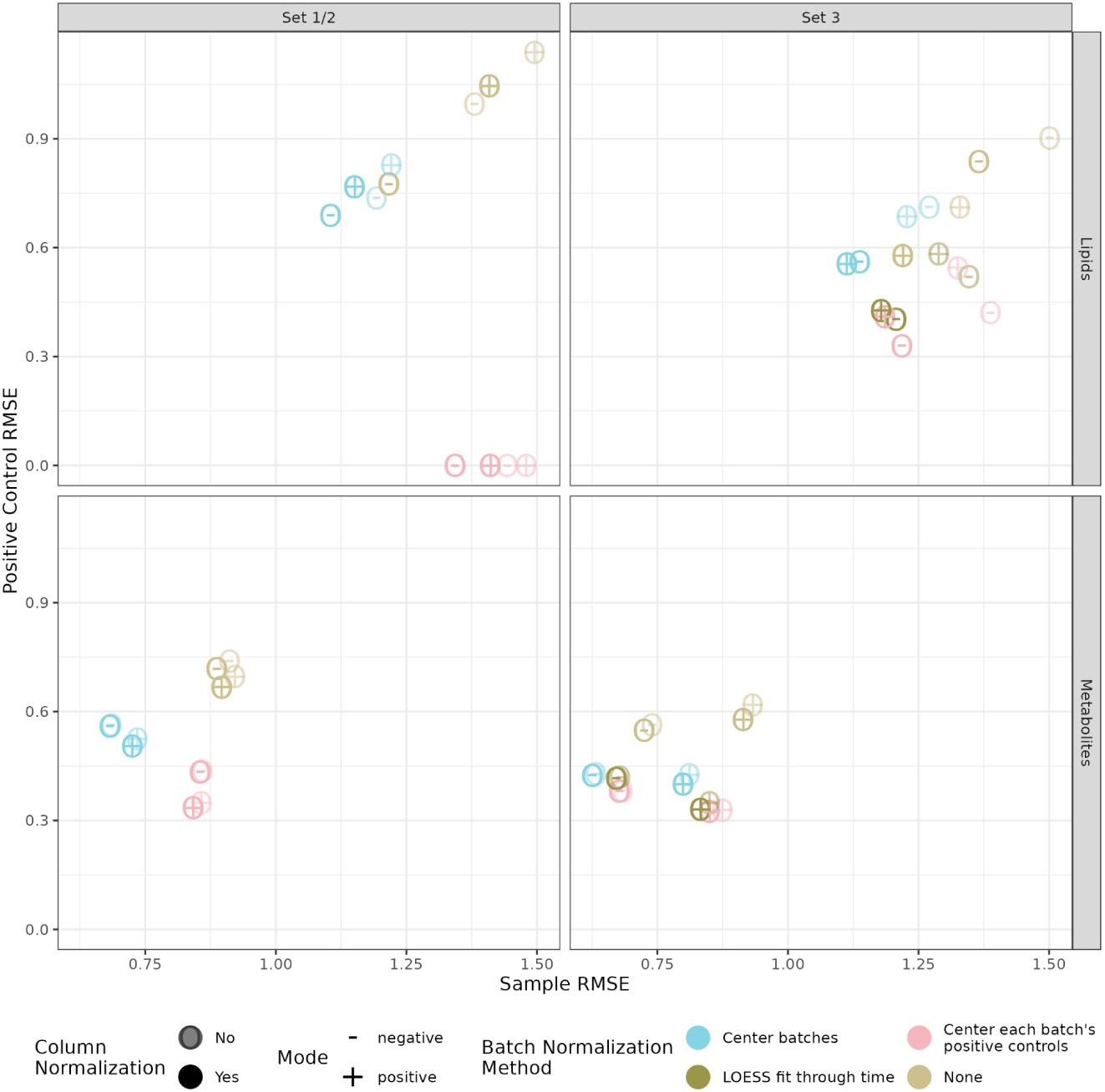
Metabolomics and lipidomics normalization methods are compared across both tranches of samples (set1/2 and set3) based on consistency of positive controls (same sample in different batches) and samples. Feature-wise batch normalization methods explored to remove “day effects” are (1) None: no adjustment, (2) Center batches: the mean value of all samples and positive controls is subtracted from each batch; (3) Center each batch’s positive controls: the mean value of all positive controls is subtracted from each batch; (4) LOESS fit through time: a LOESS curve is fit through intensity ∼ time upweighting positive controls and this fit is subtracted from each sample. Column/sample normalization, adjusting each sample across all features by a constant value, is evaluated for each batch normalization method.

**Figure SF3.**
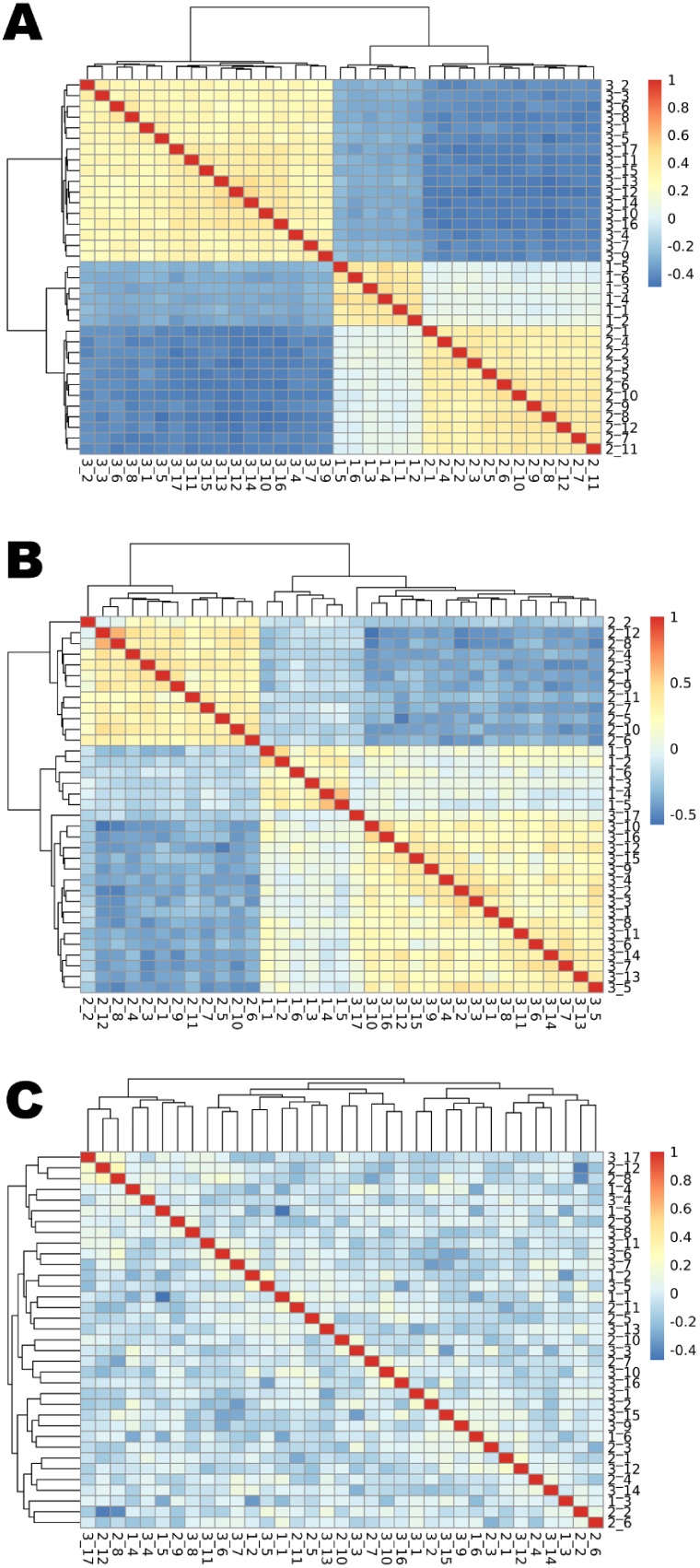
Aggregating multiple TMT plexes into a single dataset. **(A)** Correlating each plex’s “bridge” channel shows strong clustering by set due to each set being a different biological sample (a mixture of all of the experimental samples in the set). **(B)** Taking the ratio of experimental samples to their bridge removes batch effects which are shared by samples in the same plex. The mean of the sample relative abundances in each plex is correlated across plexes revealing lingering within set correlation. **(C)** Each protein is mean centered on a set-by-set basis and within the correlations between each plexes mean intensity is shown.

**Figure SF4.**
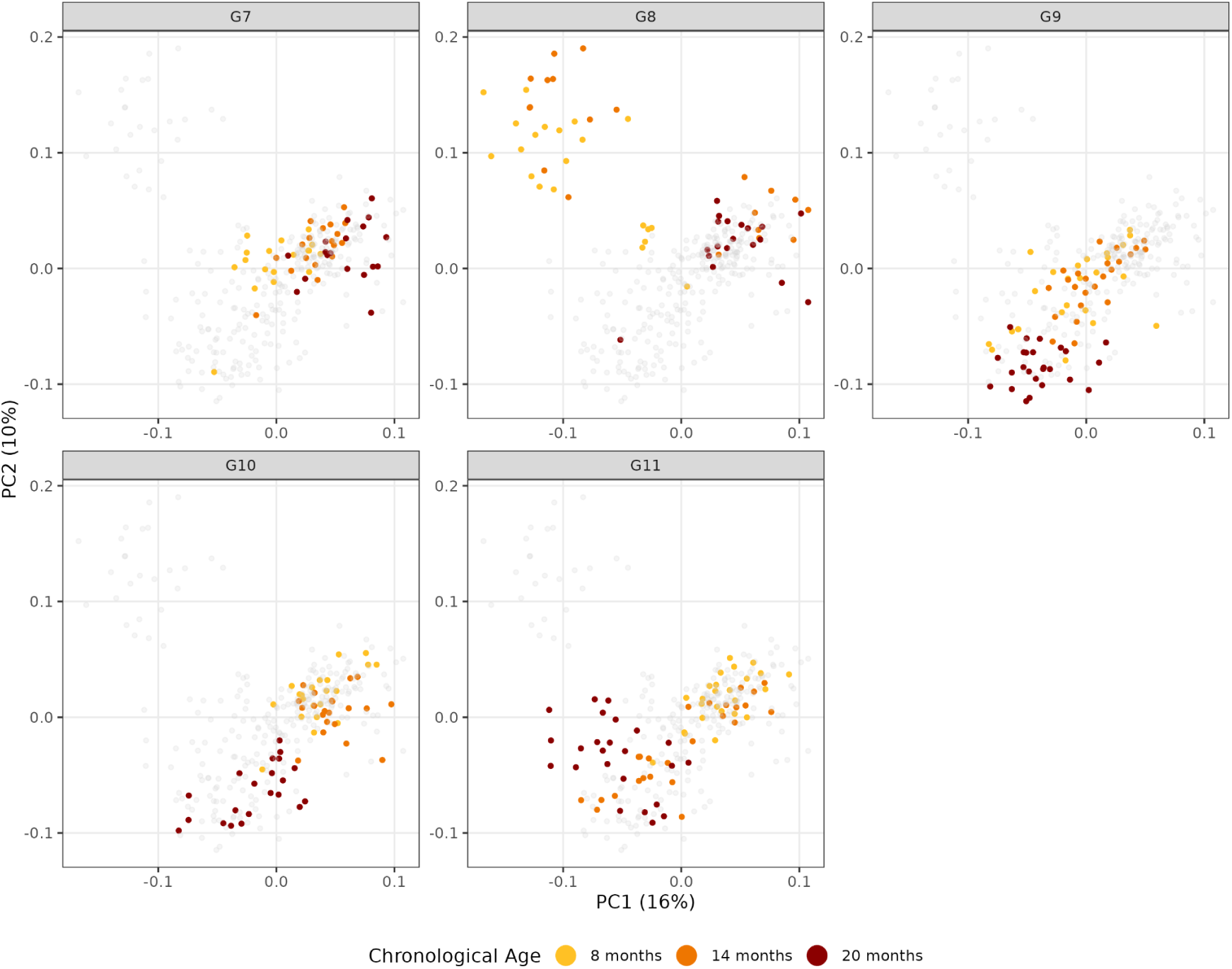
The leading principal components are overlaid with chronological age separately for each generation prior to removal of biological batch effects. These batch effects vary by mouse generation and are confounded with age. Gray dots represent mice which are not part of the highlighted generation. For instance, the G7 pane shows colored points for G7 mice and gray points for mice from all other generations.

**Figure SF5.**
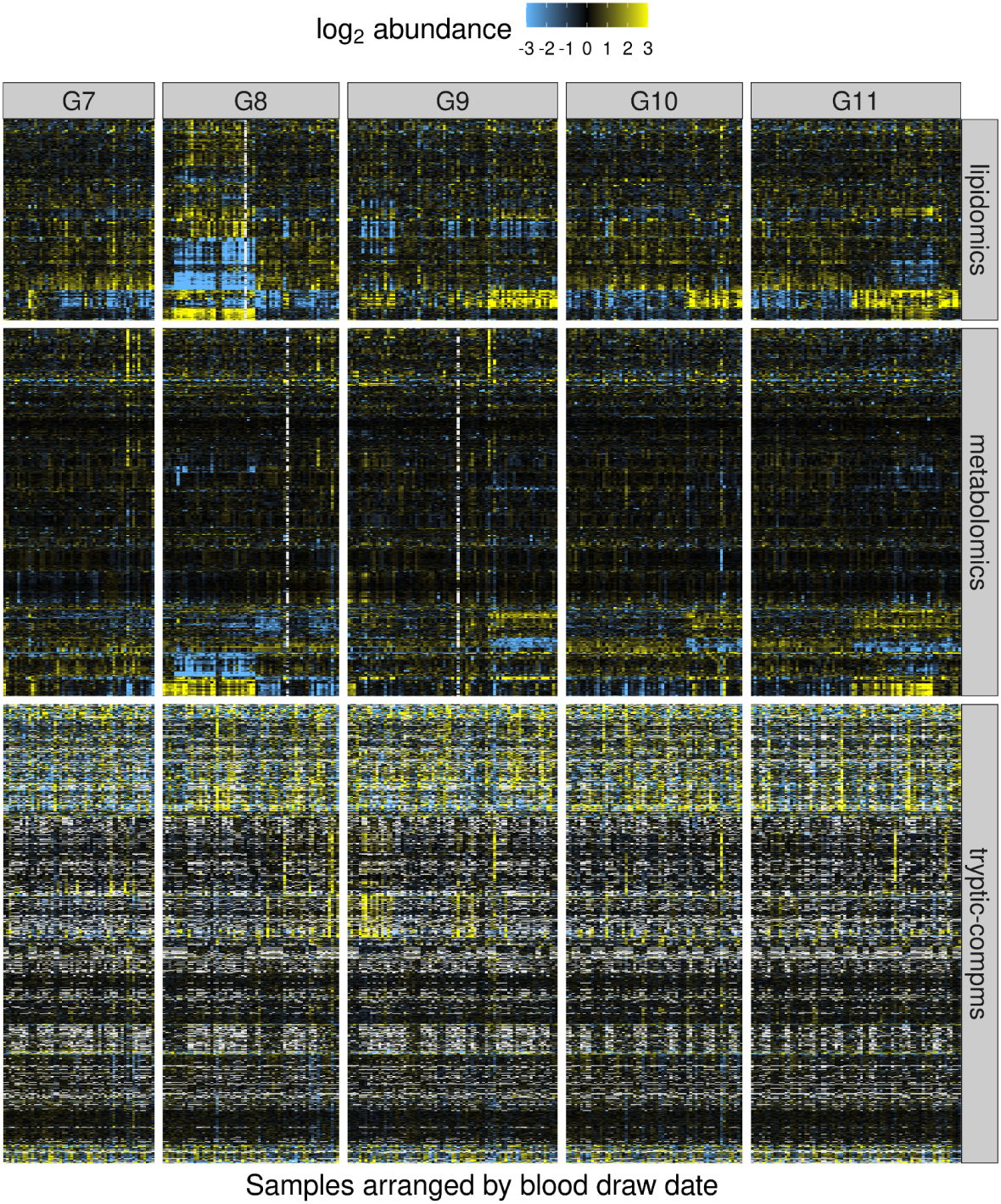
All lipids, metabolites and proteins with samples split by mouse generation and ordered by sample blood draw date prior to removal of the biological batch effects. This reveals prominent changes in the multiome of early generation 8 mice, and for late blood draw samples in generations 9, 10, and 11.

**Figure SF6.**
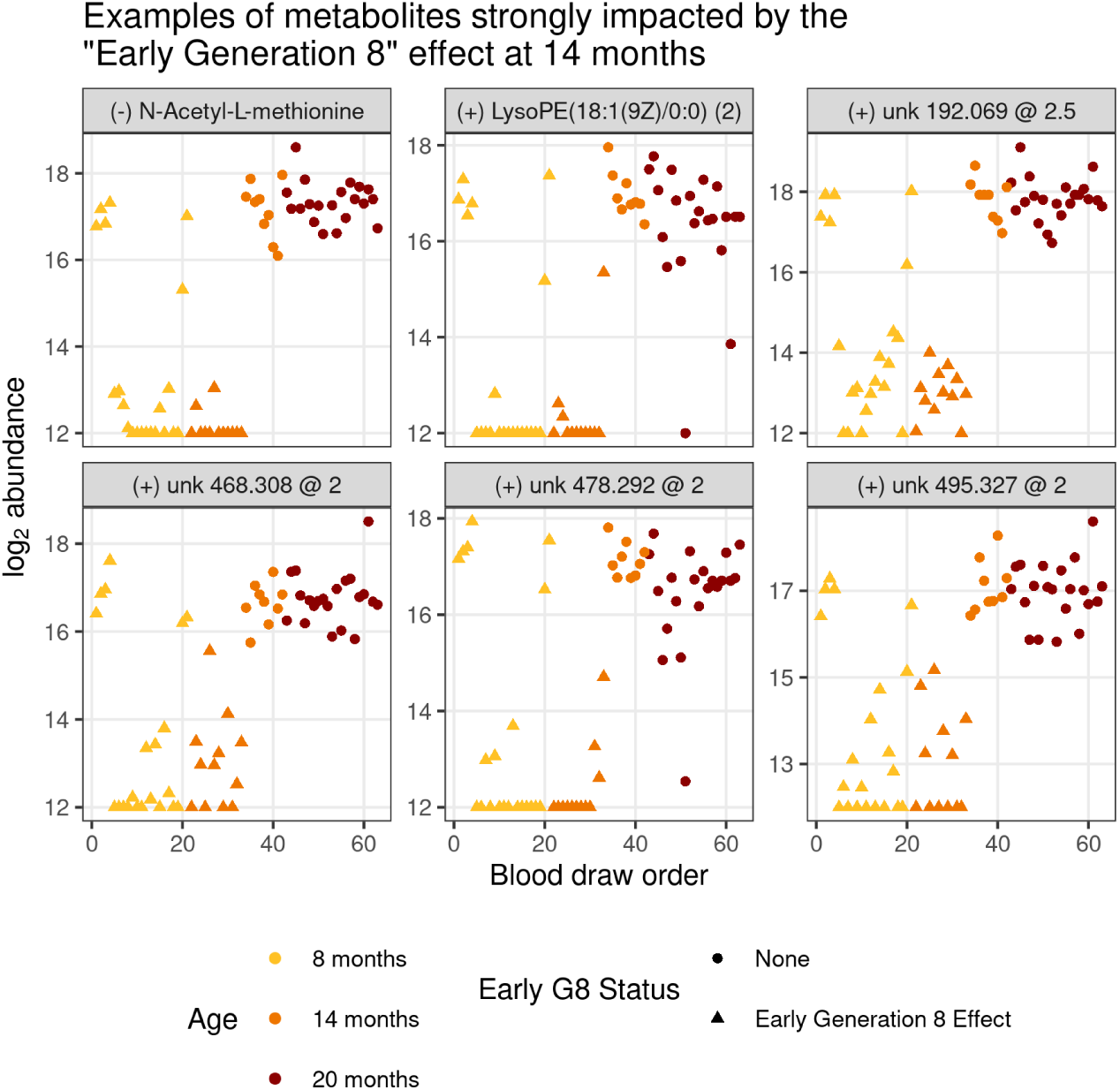
Metabolites with strong generation 8 effects are shown. The six metabolites displayed are those with the greatest fold-change between 20 month and 8 months draws of mice in generation 8 compared to the same fold-change in all other generations of mice.

**Figure SF7.**
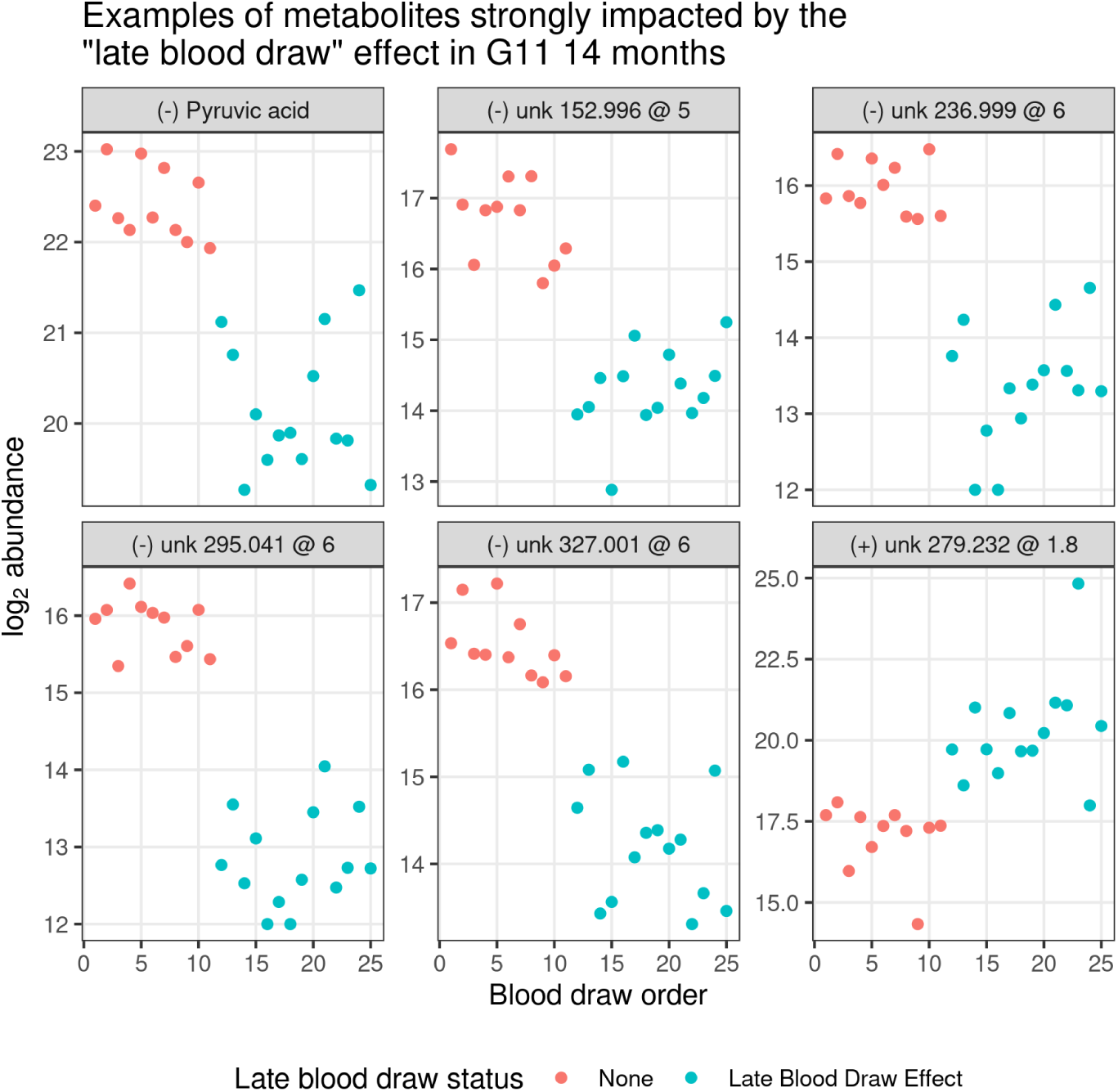
Metabolites with strong late blood draw effect are shown focusing on the 14 month blood draw of generation 11 mice. The six metabolites displayed were selected as those with the greatest difference between the average abundance at 20 months of generation 7 and 8 mice (which do not have the LBD effect) compared to the 20 month measurements of generations 9-11 mice. There is a clear breakpoint where mice before the cutoff date (red) don’t exhibit the LBD effect while those after this point reflect the same LBD effect seen in G9-11 20 month blood draws.

**Figure SF8.**
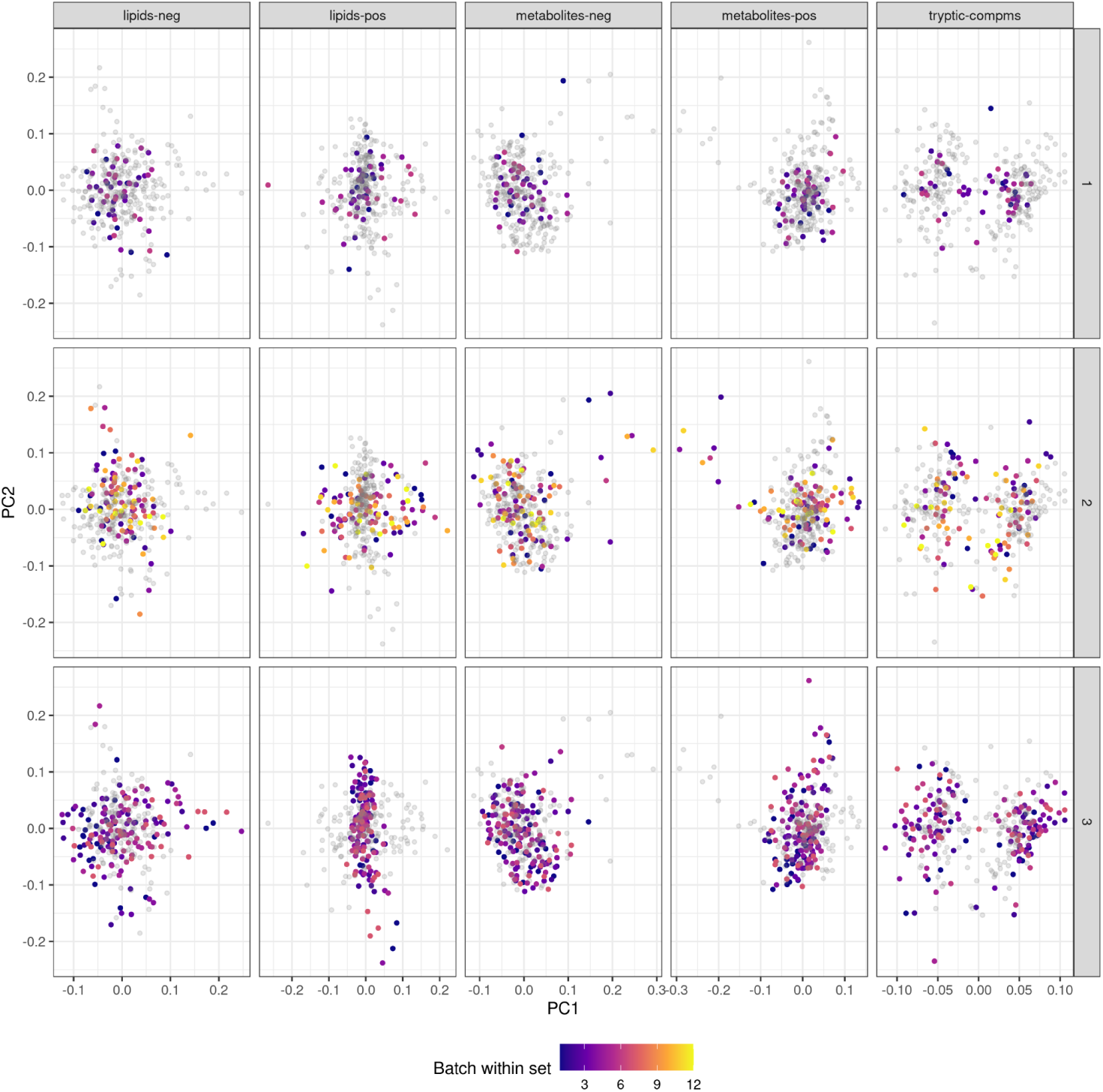
The leading components of each data type are separately calculated and overlaid by the experimental batch order. Samples’ leading principal components are relatively independent of set and batch. Principal components of different data types are not directly comparable.

**Figure SF9.**
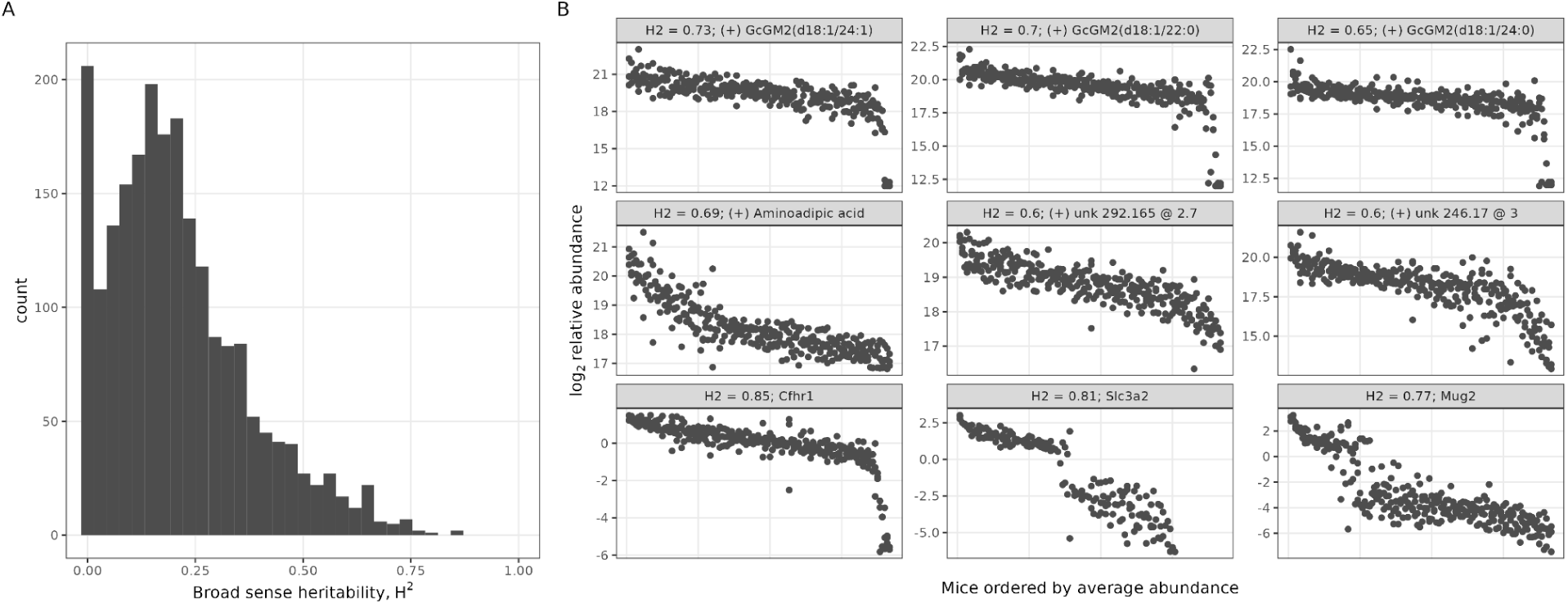
Heritability (H^2^) of proteins, metabolites, and lipids. **(A)** Distribution of H^2^ across all molecules. **(B)** Top lipids (top row), metabolites (middle row) and proteins (bottom row) by H^2^. Mice are ordered along the x-axis so that all ages for the same mouse have the same x-value, and mice are ordered by mean relative abundance.

**Figure SF10.**
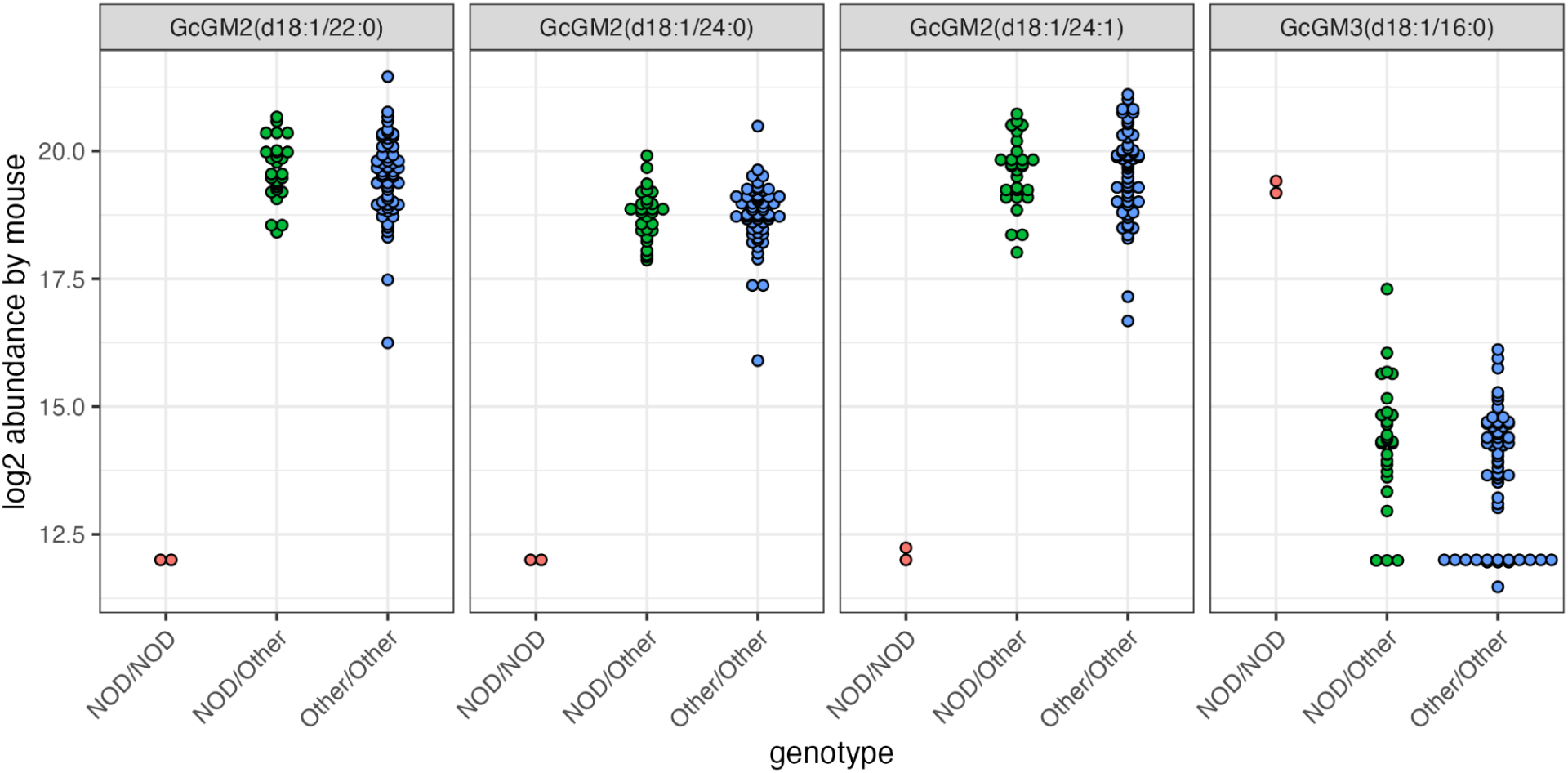
Reproducing an inborn error of lipid metabolism. The substrates of B4galnt1 (GM3 lipids) are elevated and its products (GM2 lipids) are depleted in mice which are nod/nod at the chr10:127Mb locus where the B4galnt1 enzyme is located.

**Figure SF11.**
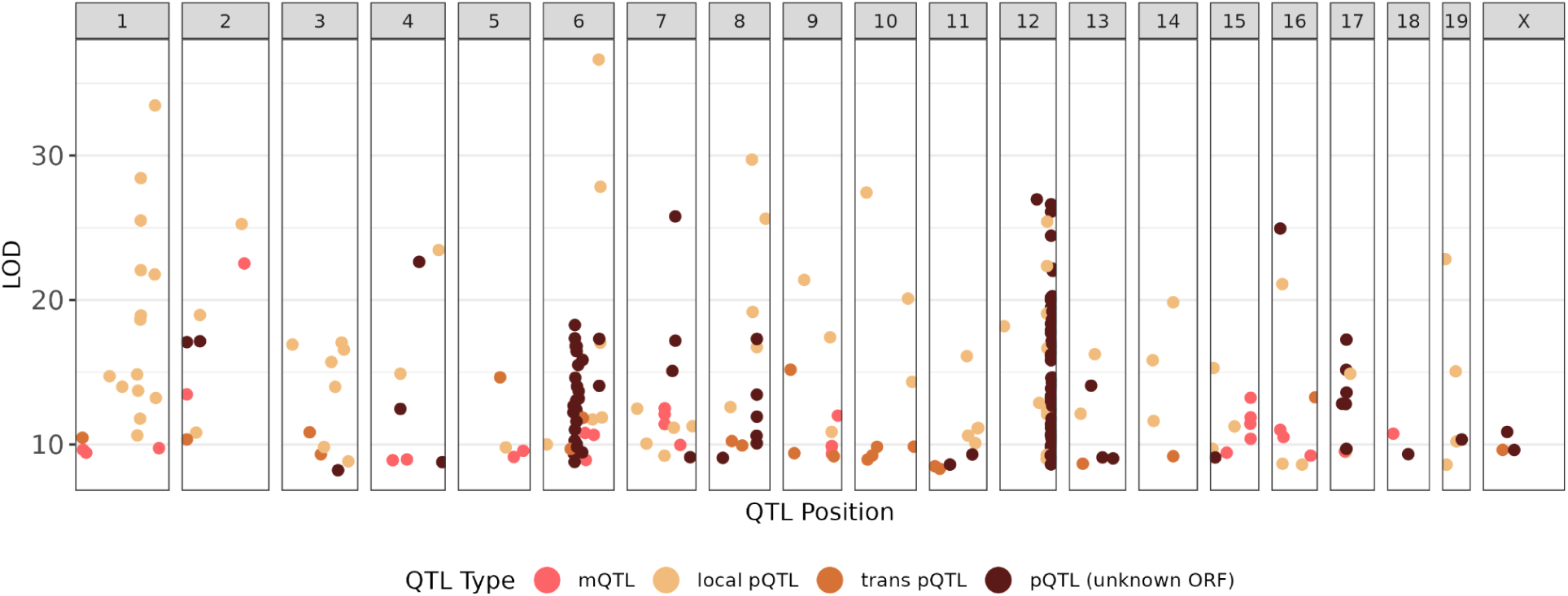
258 molecular QTLs. Protein QTLs were defined as local if they fell within 10 Mb of the gene’s starting position and trans otherwise. Some proteins, which are primarily immunoglobins were not assigned to specific genes though most of these proteins form clear QTL hotspots on chromosome 6 (kappa light chains) and chromosome 12 (heavy chains). Metabolite and lipid QTLs were jointly defined as mQTLs.

**Figure SF12.**
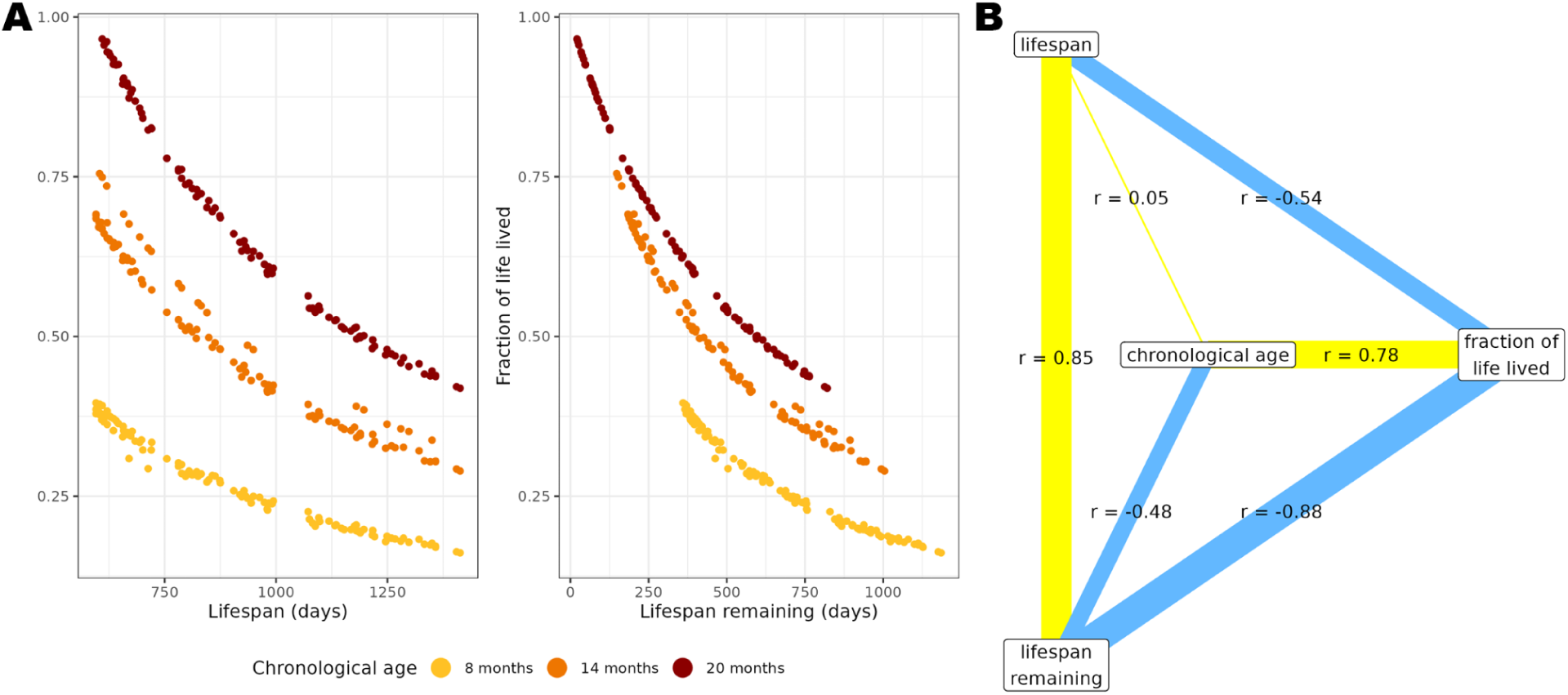
Aging archetypes are (anti)correlated by construction. Results are shown for all 320 non-DDM samples in the study. Age x lifespan interactions cannot be reduced to an informative univariate summary and unlike other archetypes they are not (anti)correlated by construction, hence they were excluded from this analysis. **(A)** Scatterplots contrasting aging archetypes. **(B)** Pearson correlations of aging archetypes across samples.

**Figure SF13.**
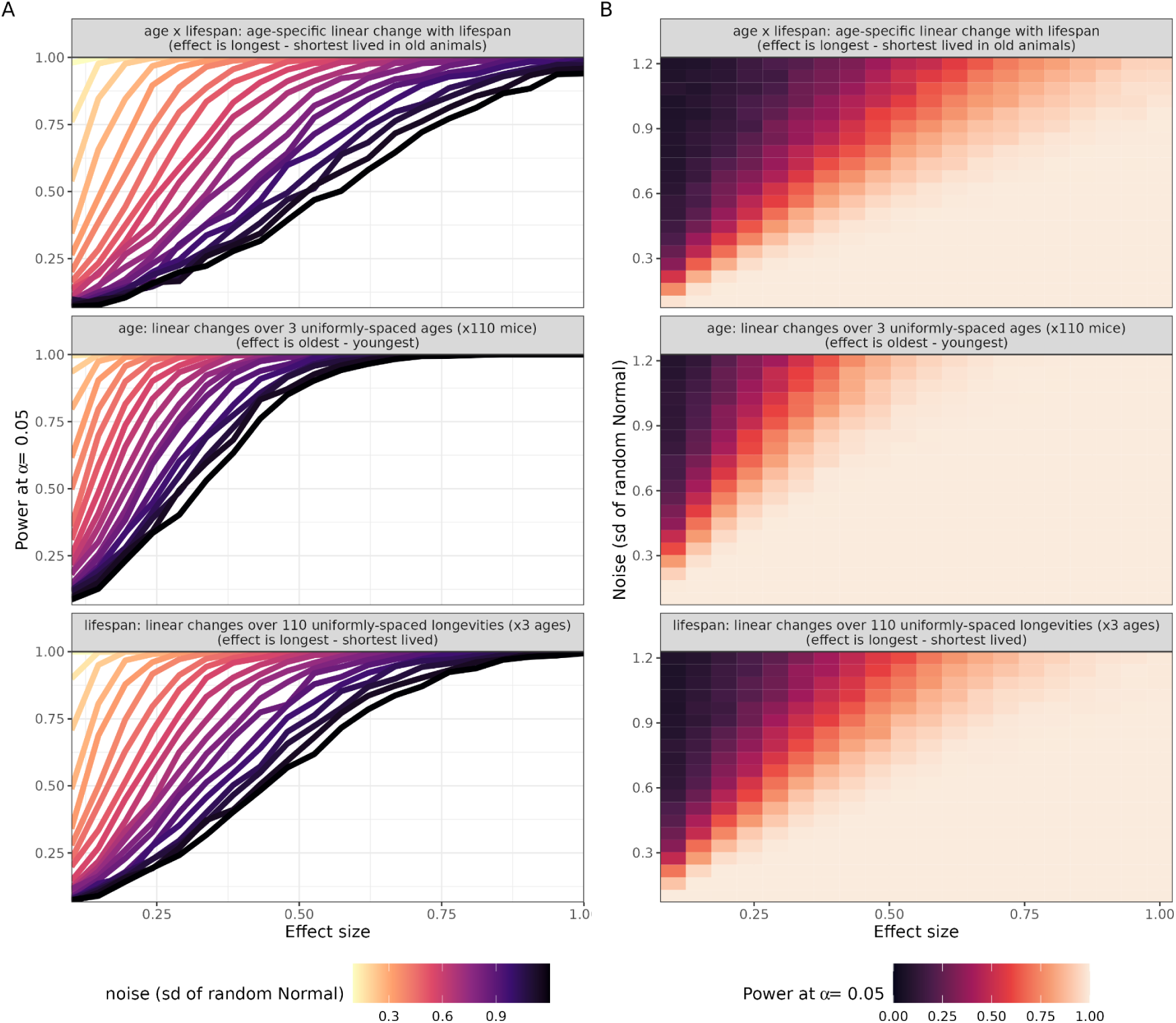
Power analysis for detecting differential abundances with age, lifespan, and age x lifespan. The relationship between effect size, noise, and power is shown two ways for age, lifespan, and age x lifespan effects. **(A)** The relationship between effect size (x-axis) and target power (y-axis) with different noise levels (curves) is shown. **(B)** The power to detect an effect size of interest (x-axis) given a noise level (y-axis) is shown.

**Figure SF14.**
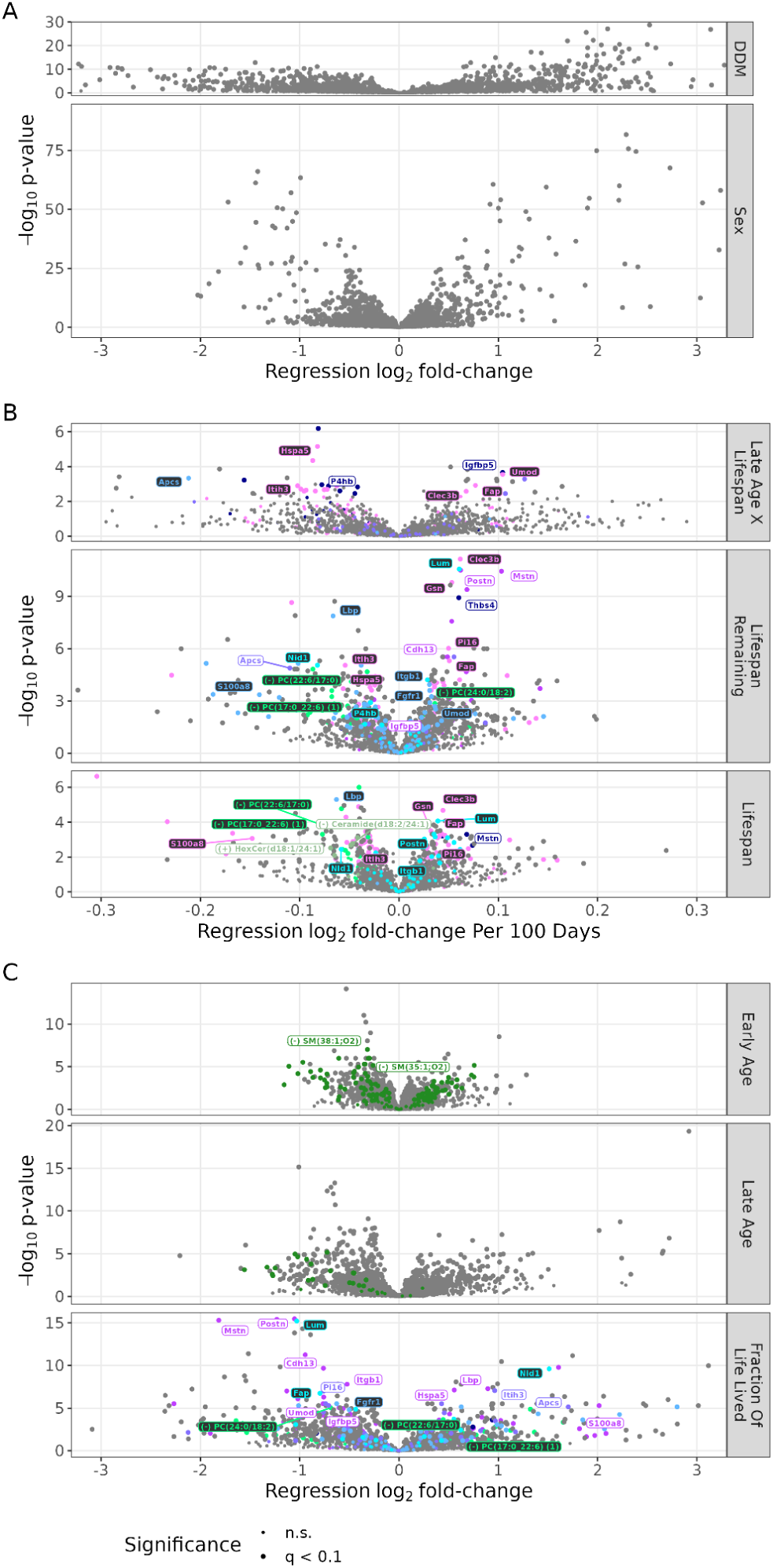
Volcano plot of molecules’ changing with sex, DDM, and aging archetypes. **(A)** Features changing with age and sex; the x-axis shows log_2_ effect sizes comparing two categories in a feature-level regression (i.e., male - female, ddm 20 month fold-changes vs. non-ddm 20 month fold-changes). **(B)** Features changing with aging archetypes with per-day units. Members of pathways which are significantly enriched for each term are colored irrespective of significance and key associations are labelled. **(C)** Features changing with non-per-day units. Early age contrasts 14 and 8 month draws from the same mouse; while late age contrasts 20 and 8 months. As for (B), members of enriched pathways are colored and representative molecules are labeled.

**Figure SF15.**
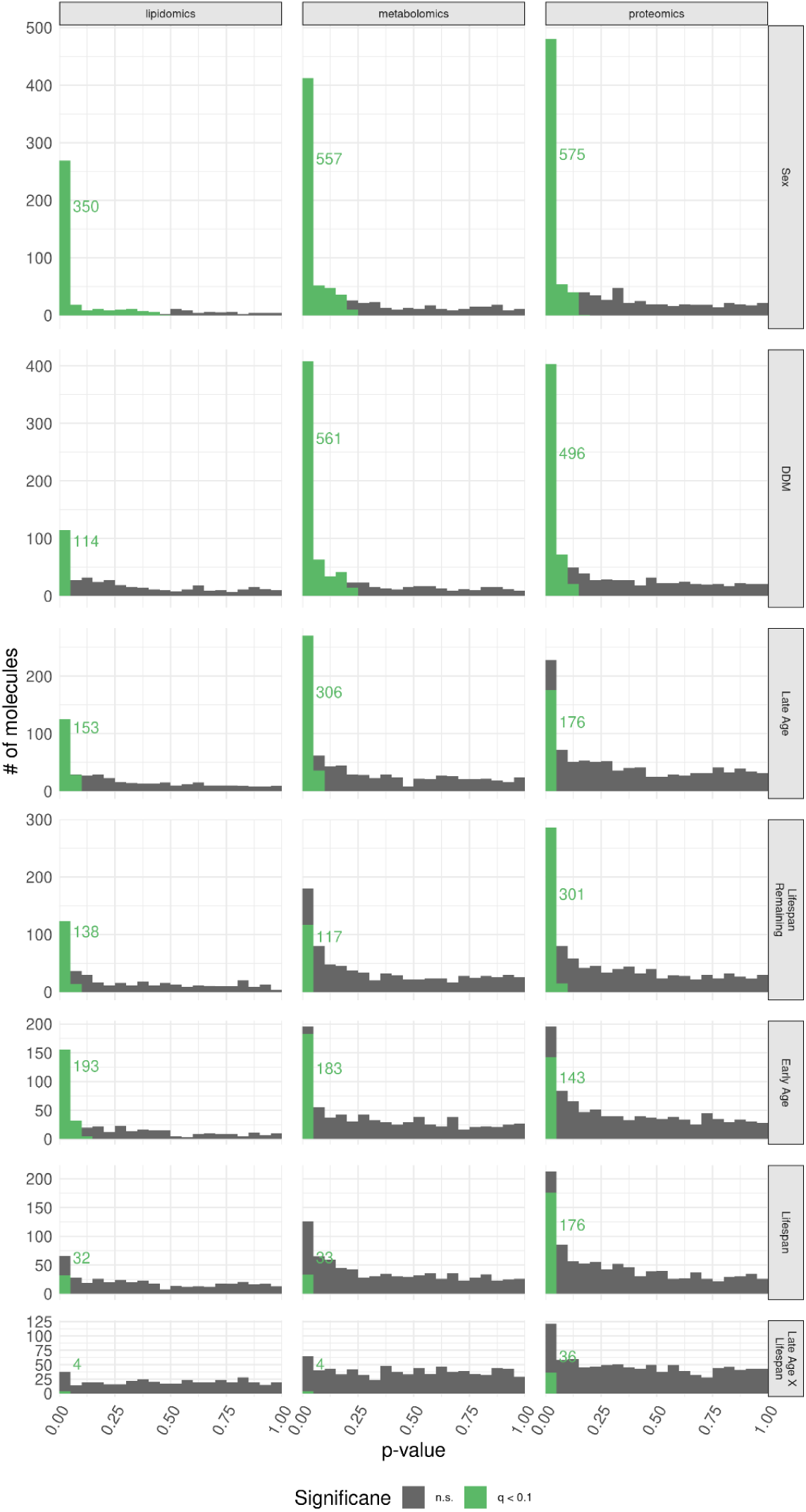
P-value histograms of each major effect split up by data modality.

**Figure SF16.**
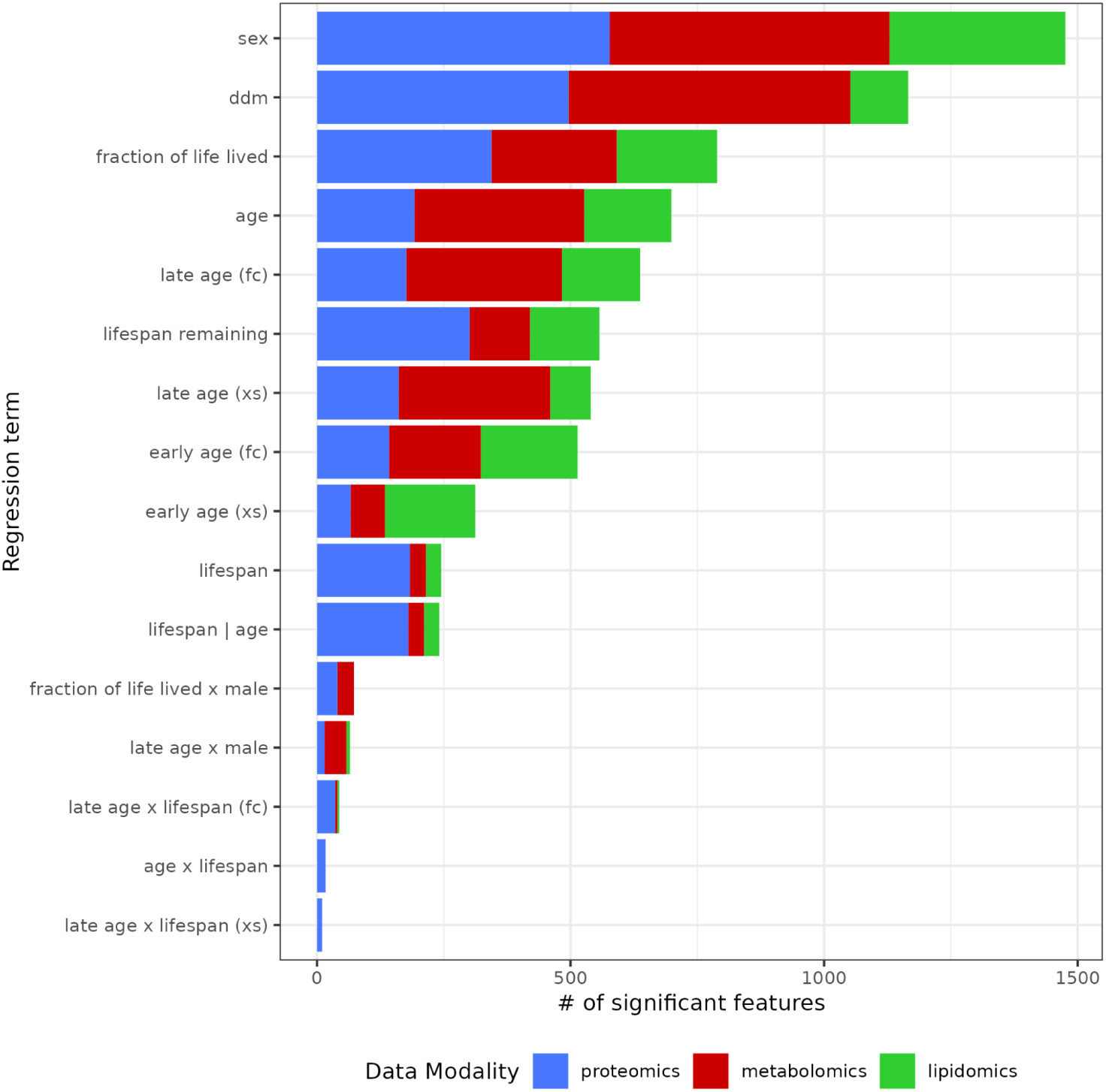
Counts of significant discoveries at q < 0.1 including alternative regression models. For descriptions of all alternative models see supplemental methods. Some supplemental models (e.g., cross-sectional models (xs)) were fit to enable model comparison of alternative models fit to identical data. Age and age x lifespan models that include age as a linear rather than a categorical variable were fit to enable the calculation of pathway delta age.

**Figure SF17.**
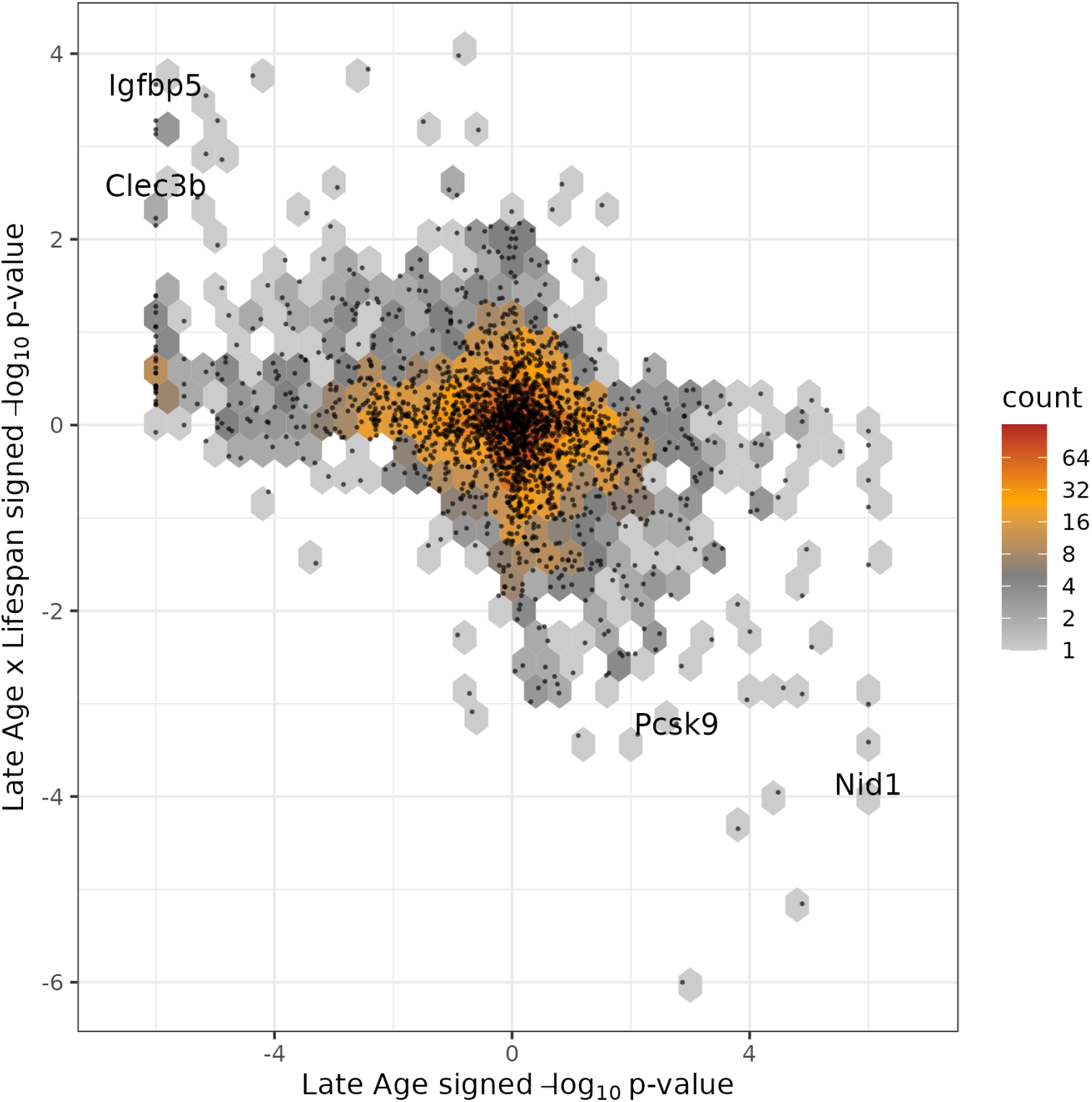
Hexbin plot demonstrating the anticorrelation of aging and age x lifespan interactions. Signed p-values comparing all ∼2,200 feature’s late age and late age x lifespan effect sizes are shown as a hexbin plot (i.e., a bivariate histogram) with a scatterplot of individual molecules overlaid.

**Figure SF18.**
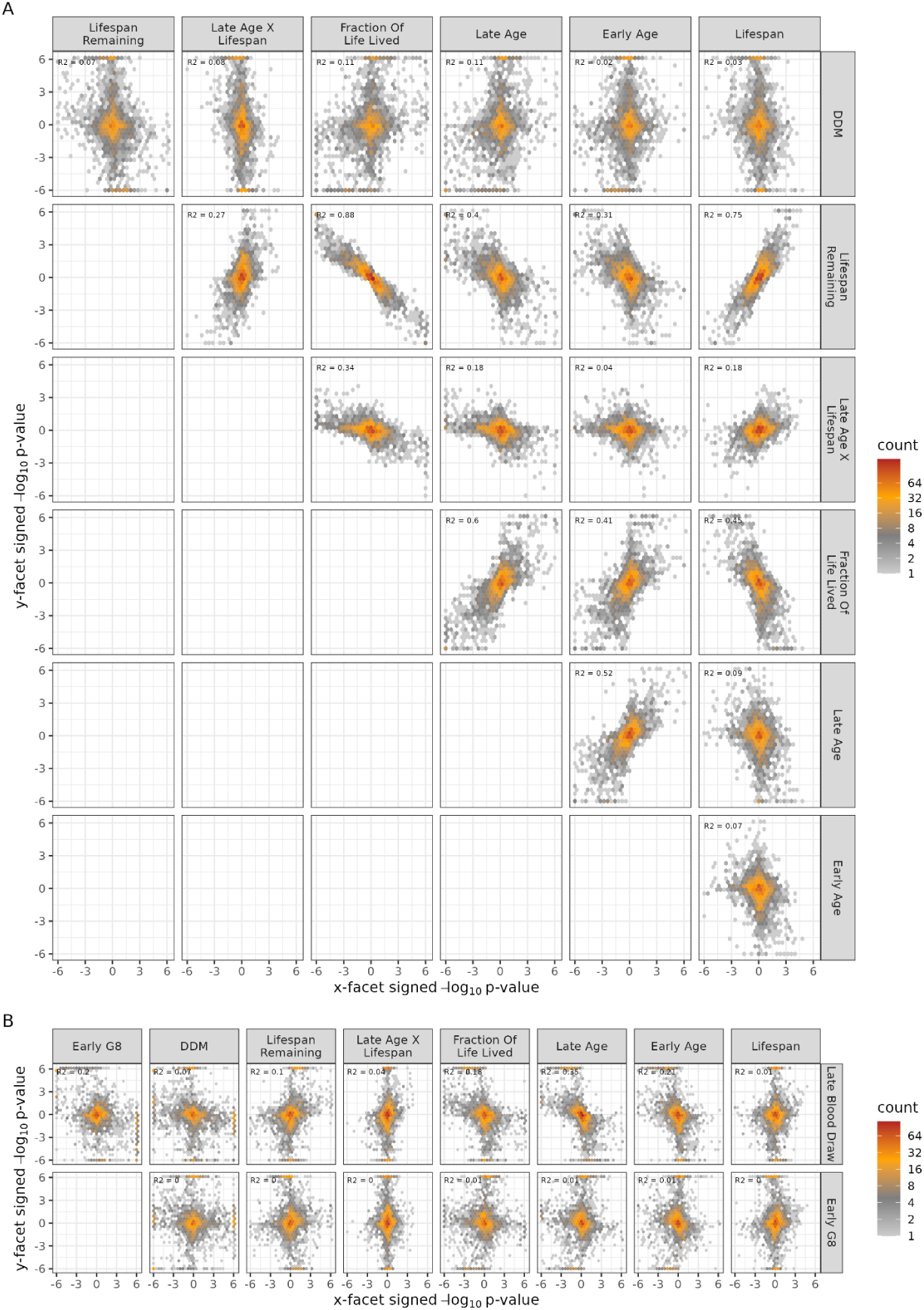
Hexbin plots comparing signed -log10 p-values between pairs of regression terms. Hexbin plots can be thought of as bivariate histograms where the number of observations falling within each hexagon is summarized by the shade of the hexagon itself. **(A)** Comparing effect estimates of all pairs of aging archetypes (and the DDM). Some aging archetypes are (anti)correlated by construction such as lifespan and lifespan remaining, while others such as late age effects compared to late age x lifespan effects show a non-trivial empirical (anti)correlation. **(B)** Comparing terms estimating in (A) to the late blood draw and early G8 batch effects. Batch effects overlap with strong chronological aging and FLL associations.

**Figure SF19.**
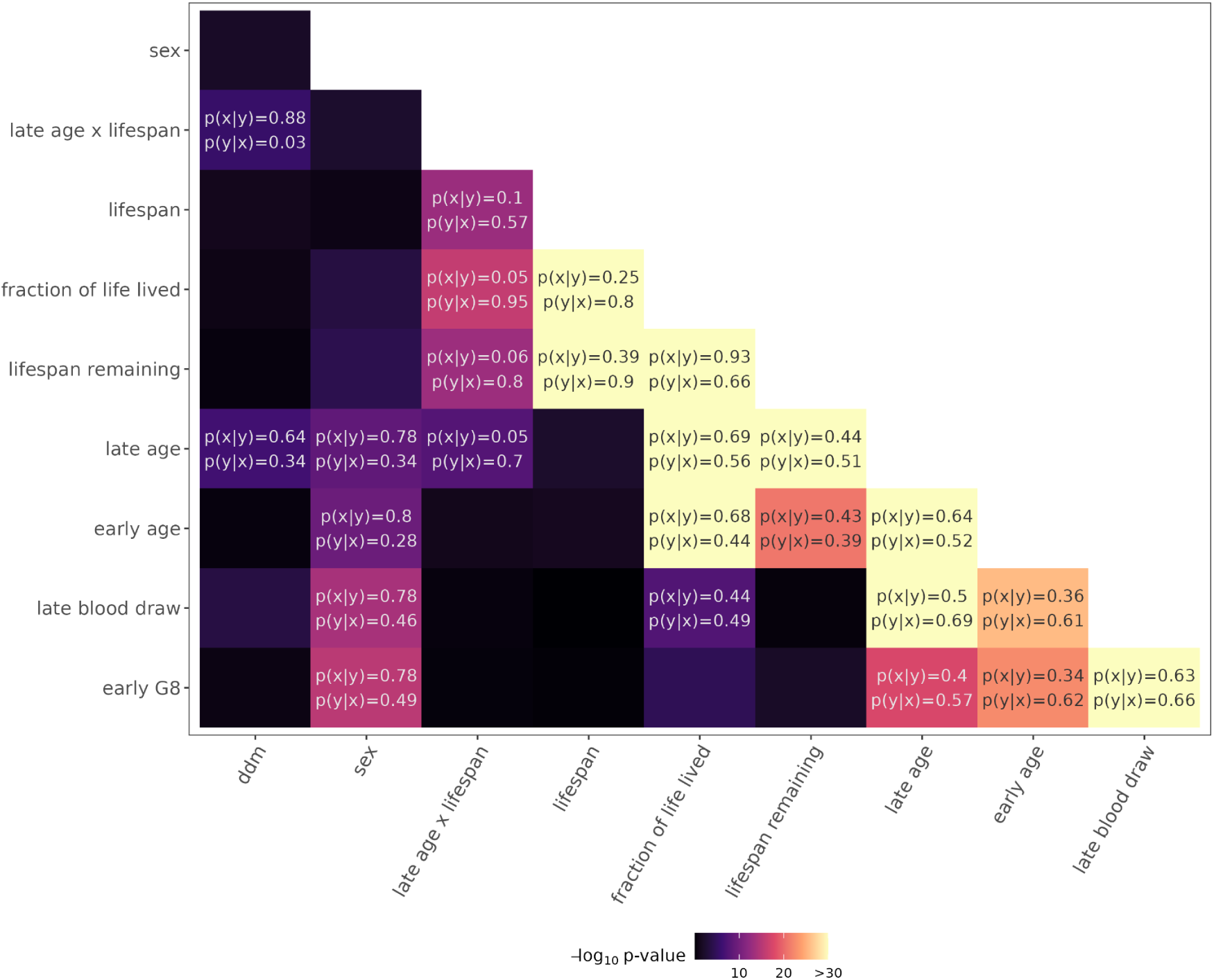
Identifying pairs of effects which tend to be shared by molecules. Pairs with strong enrichment based on a Fisher Exact test are labelled along with p(x|y) and p(y|x). p(y|x) reflects the fraction of molecules which are significantly associated with a row term given that they are significant for a column term. p(x|y) is the converse. As expected from its construction, aging hits and lifespan hits are both likely to influence lifespan remaining. There is also a moderate overlap of lifespan and age x lifespan interactions where ∼50% of the smaller set of significant age x lifespan interactions also show up as lifespan associations. Age is partially entwined with the early G8 and late blood draw batch effects, but such effects are noticeably absent among hits for most other effects.

**Figure SF20.**
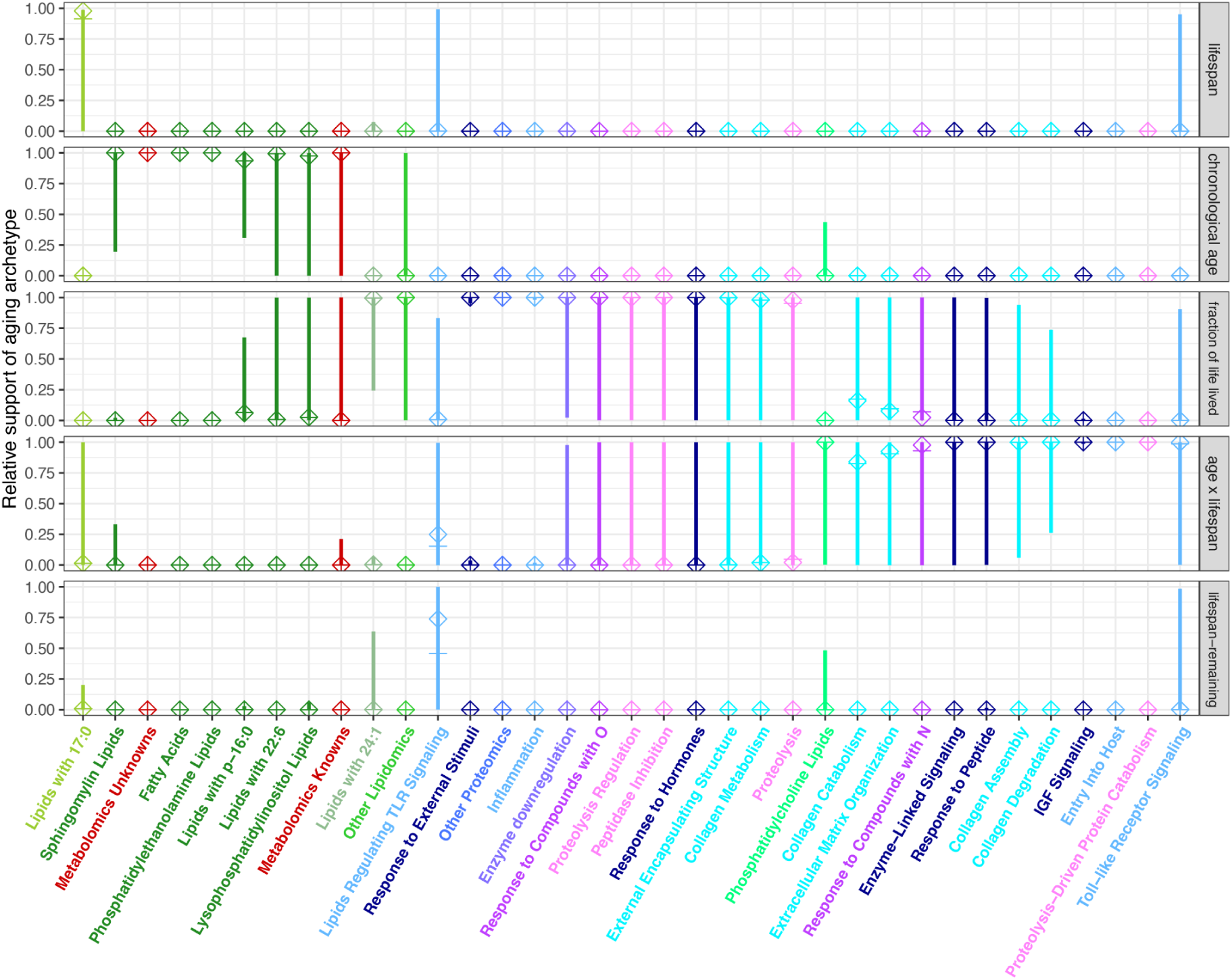
Model comparison of aging-associated pathways across aging archetypes. Each archetype’s total support was calculated by summing AICc over all pathway members, and model’s relative supports are shown as diamonds on the y-axis. To assess the uncertainty in these archetype assignments we resampled each pathway’s members with replacement 1,000 times and performed the same calculation of relative pathway support across archetypes. The lower 2.5% to 97.5% interval is shown as a range and the median value is a cross.

**Figure SF21.**
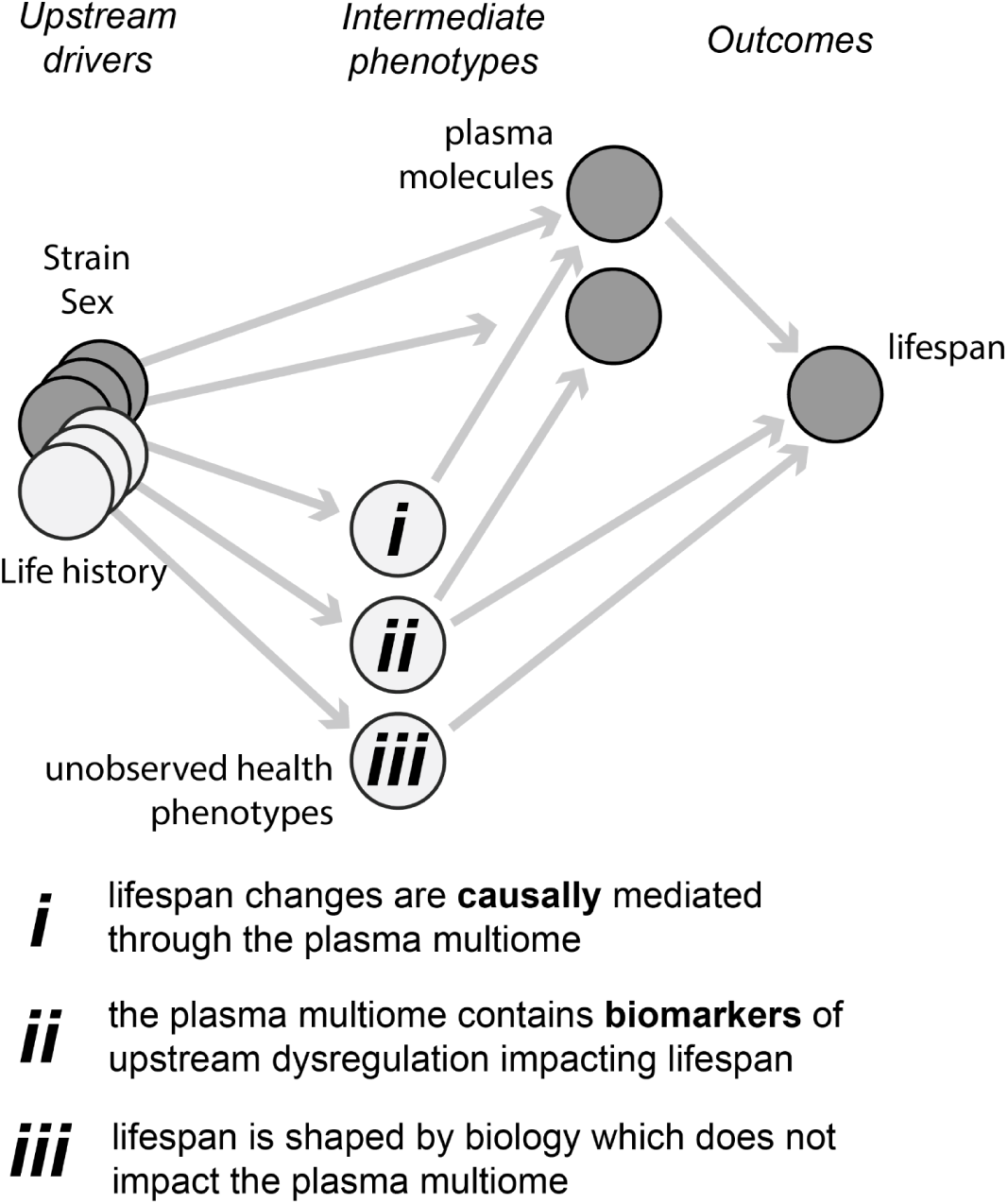
The plasma multiome is structured by mice’s genetic background and life history. Much of this heterogeneity impacts the multiome independent of health-relevant changes in physiology but some effects will be transmitted through unobserved health phenotypes. Most plasma longevity associations are likely biomarker readouts of these underlying differences in health but some may be responsible for causally mediating effects responsible for differential hazard.

**Figure SF22.**
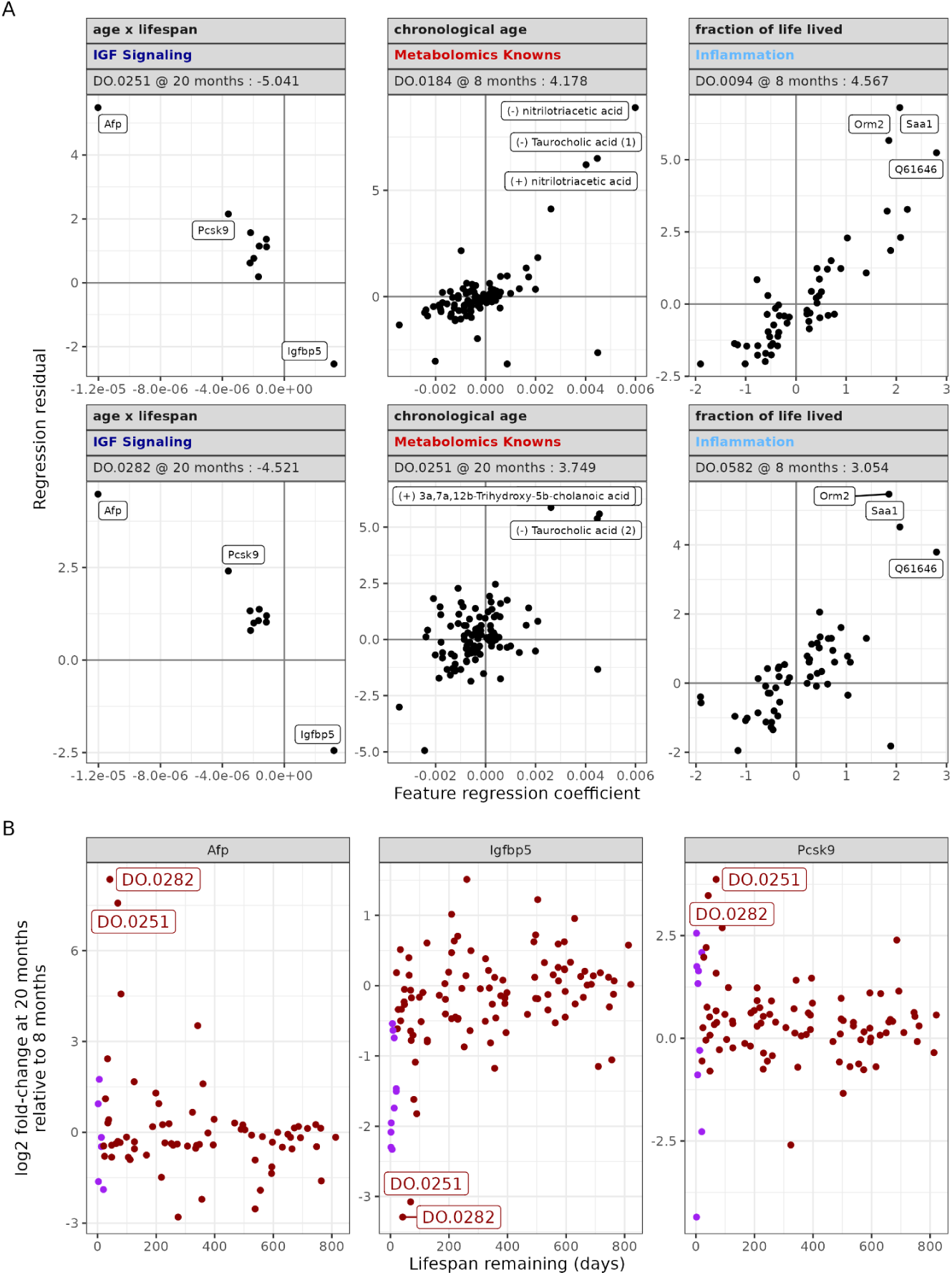
Extreme pathways based on delta age reflect (anti)correlation of pathway-associated aging metabolites. **(A)** Each panel compares a pathway to an aging archetype for a single mouse to relate measurements of pathway member’s regression residual (y-axis) to the corresponding molecule’s regression coefficient. An anticorrelation reflects that a mouse has a more negative value of the archetype than their known age and lifespan would suggest (for archetypes which are anticorrelated with FLL like age x lifespan this would be interpreted as accelerated aging). A correlation suggests a higher value which for chronological age and FLL can be directly interpreted as premature aging. **(B)** In panel (A), Afp, Pcsk9 and Igfbp5 have an outsized impact in estimating pathway delta age relative to some other pathway members which are near the scatterplot origin. These three molecules are shown compared to fraction of life lived labeling the two mice shown as extreme examples of IGF signaling dysregulation in (A). DDM samples are highlighted in purple.

**Figure SF23.**
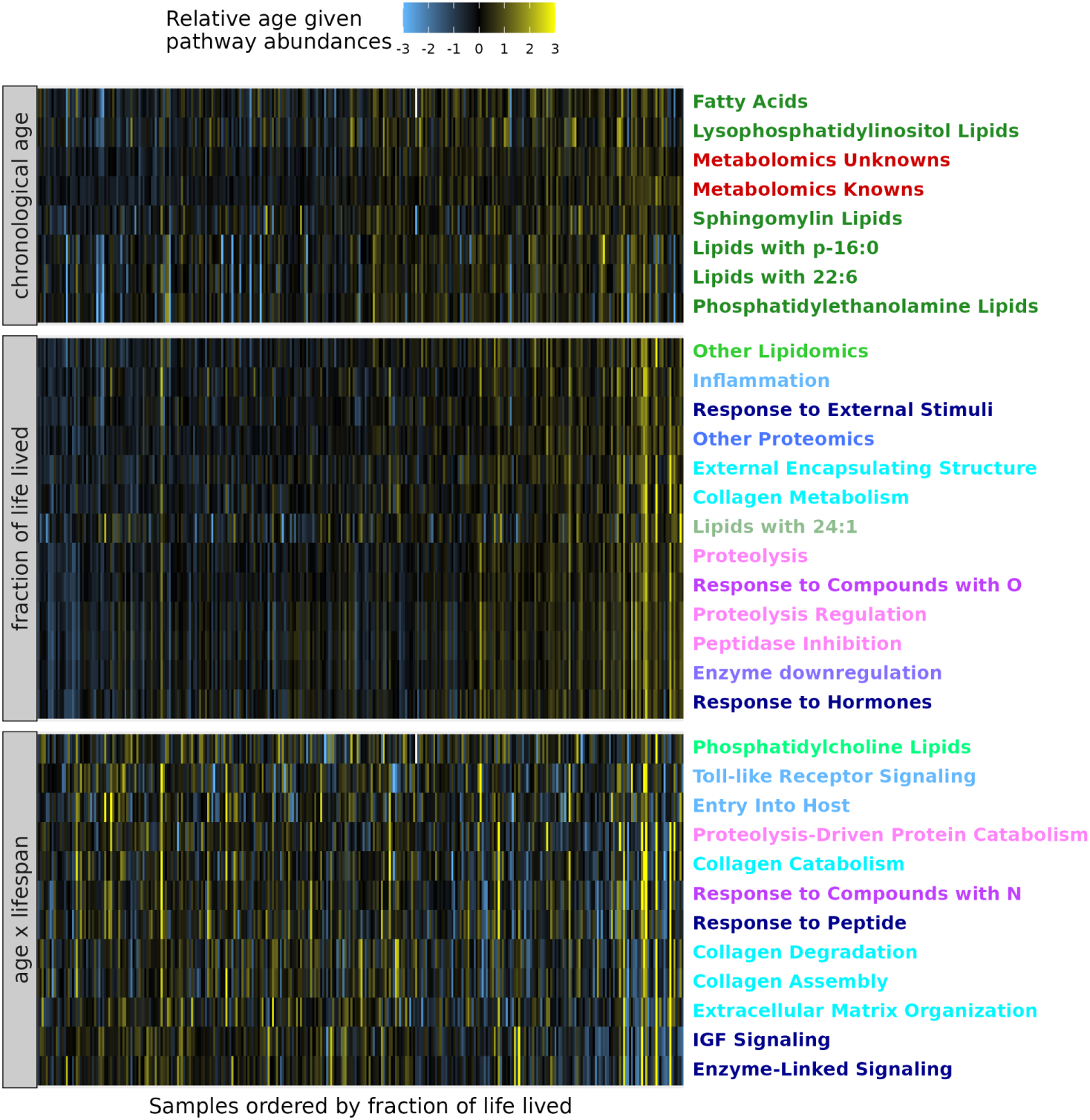
Sample-level biological age estimates based on individual pathways aging archetype. Pathway delta age was calculated for each pathway compared to its best supporting archetype (excluding the two pathways associated with lifespan and lifespan remaining). Pathway delta age was added to measured chronological age and FLL, while delta age was directly visualized for age x lifespan interactions. Rows were standardized and hierarchically clustered based on Pearson correlation.

**Figure SF24.**
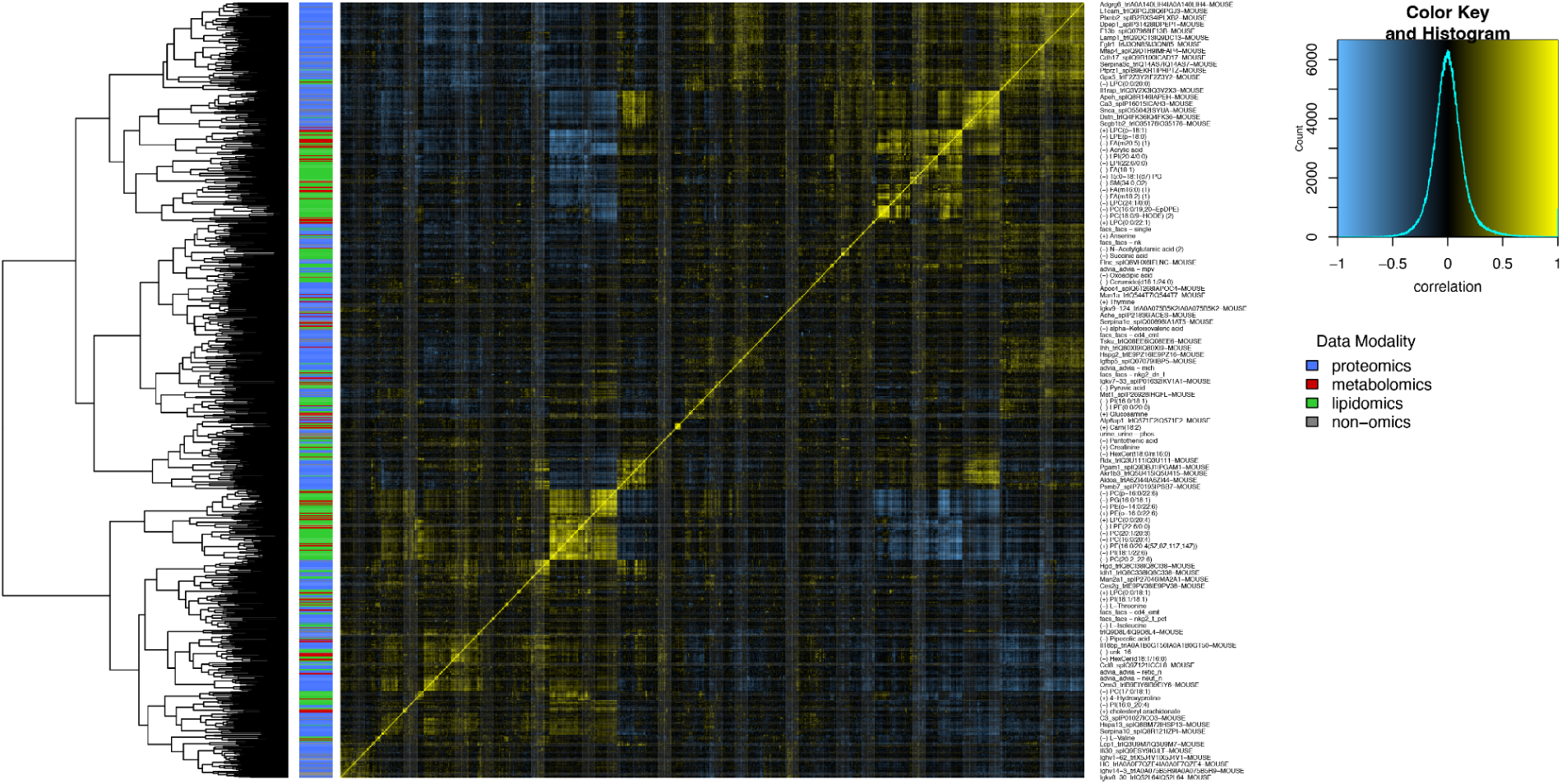
Hierarchically clustered heatmap of lifespan-associated molecules including non-molecular phenotypes.

**Figure SF25.**
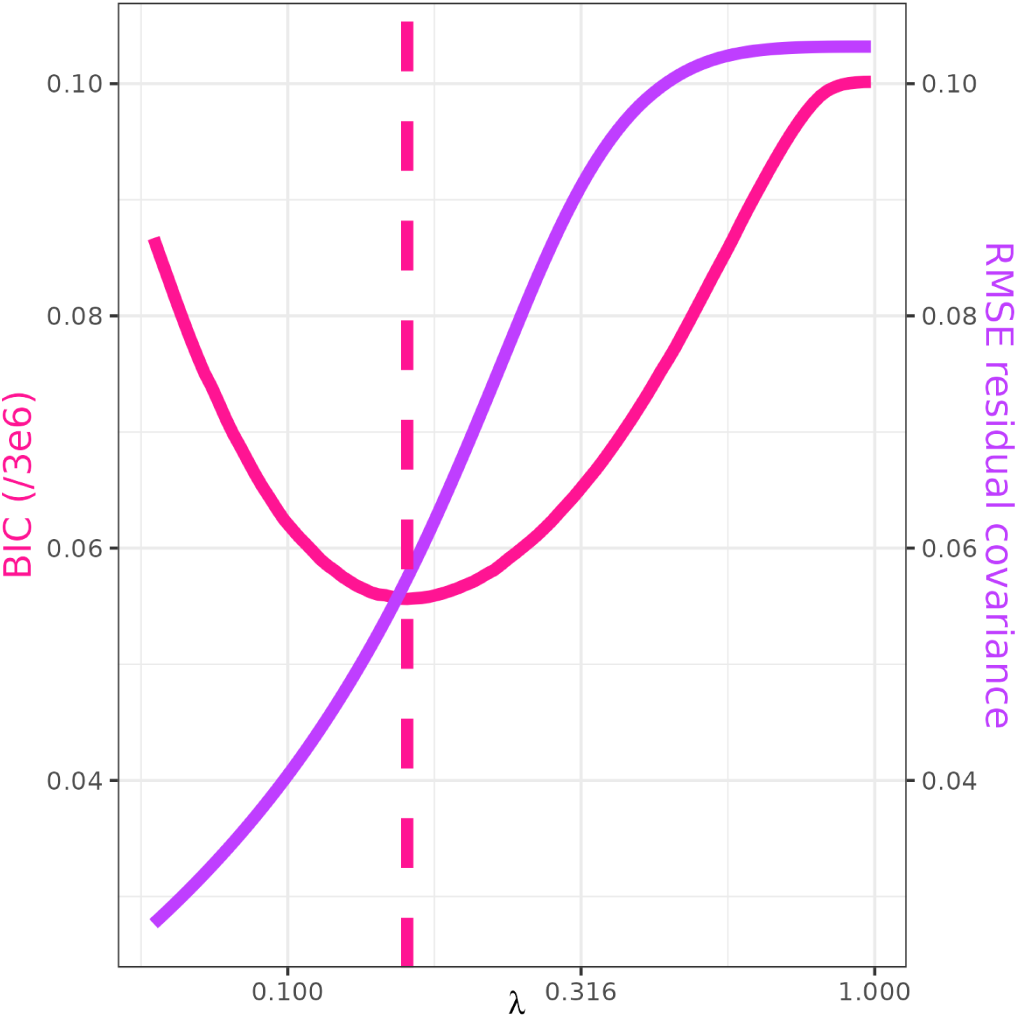
Estimating a sparse precision matrix using graphical LASSO. The model BIC and RMSE of residual covariance are shown as a function of regularization strength. The lambda which minimizes the BIC was selected as an optimal model and this value is observed to capture much of the structure in the underlying sample covariance matrix.

**Figure SF26.**
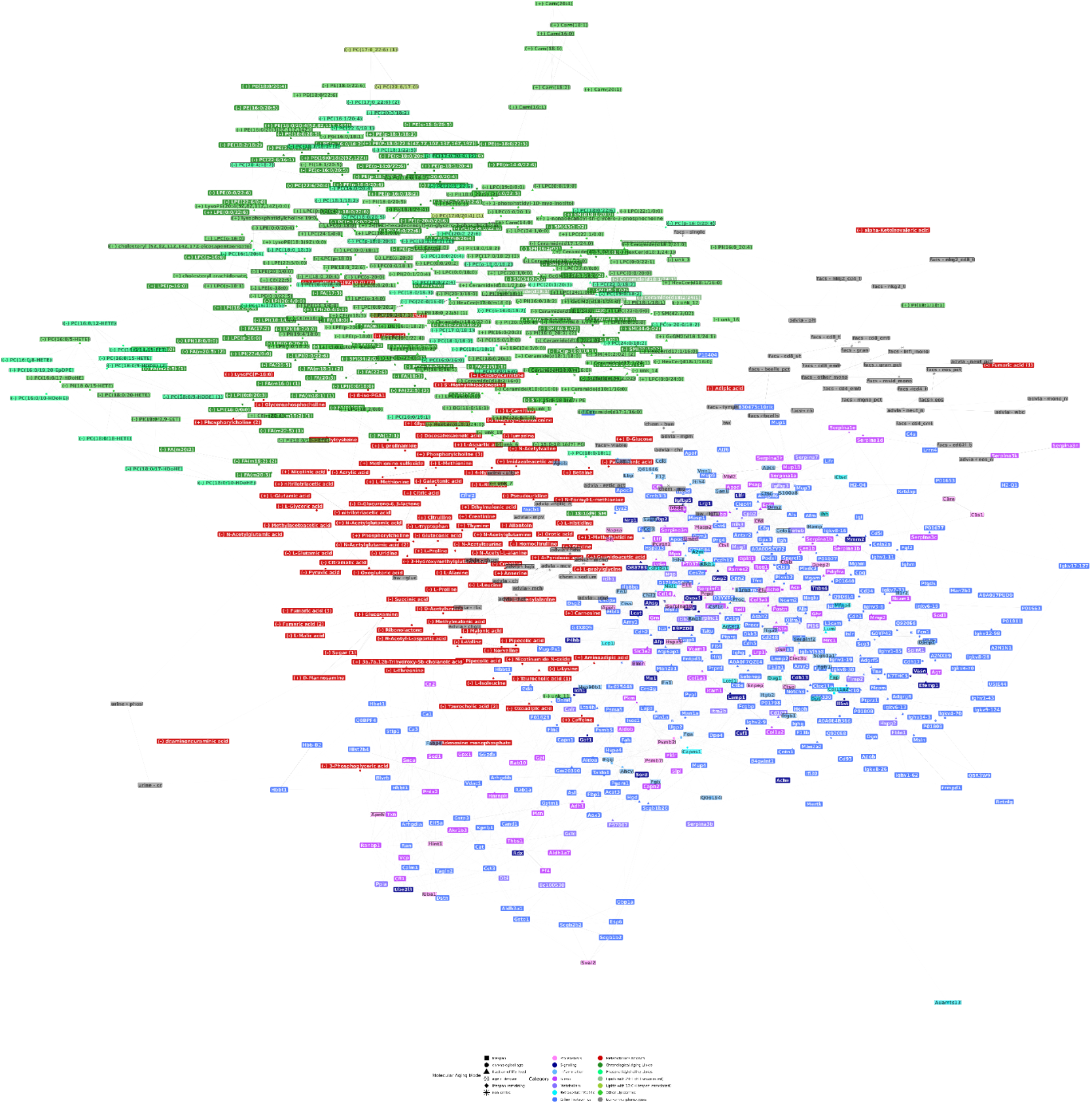
Figure 5B with all nodes labeled.

**Figure SF27.**
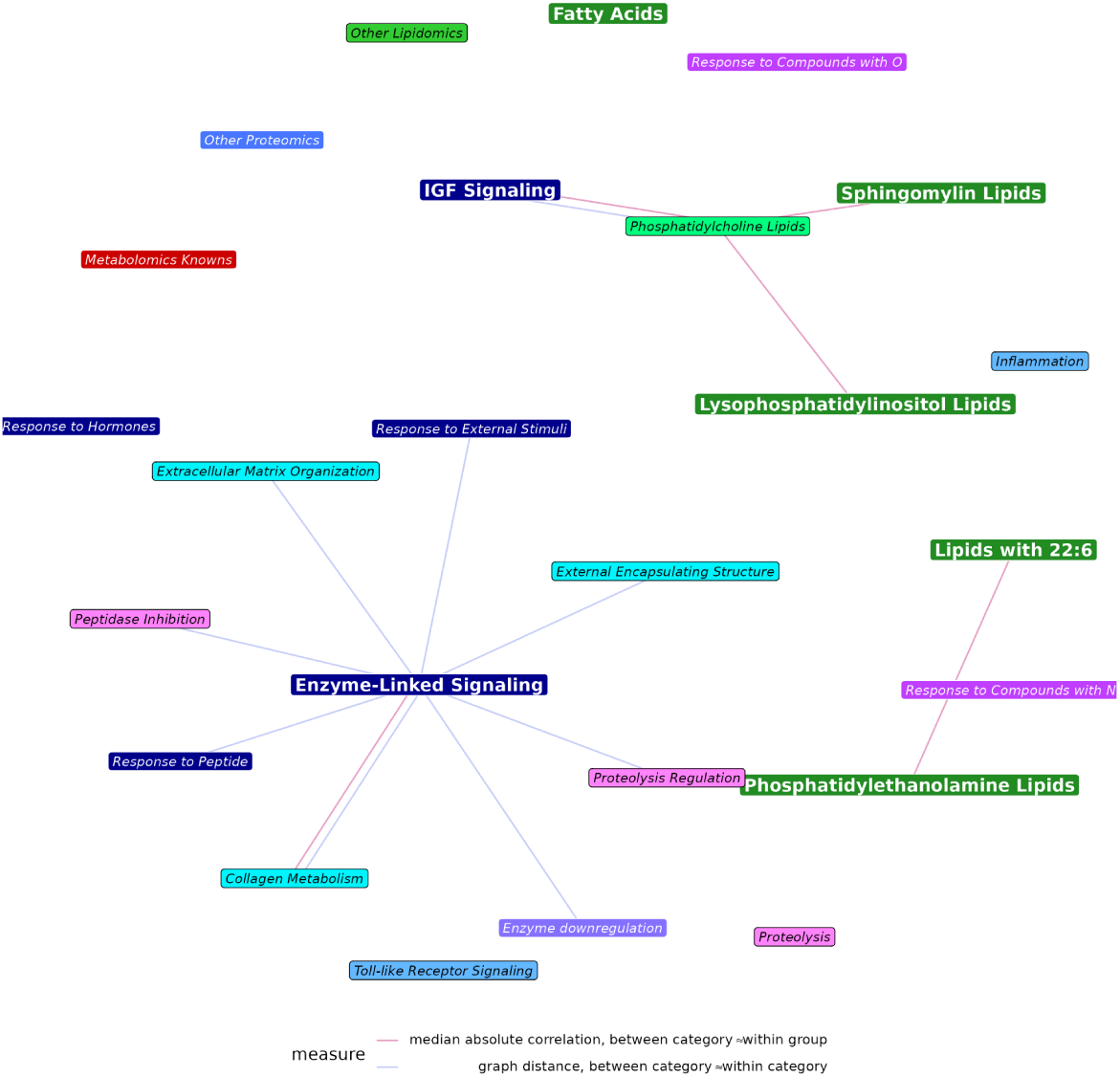
Comparing the similarity of molecules within versus between pairs of functional categories. To determine whether two categories are more coherent when separated or combined together each pairs’ within category median absolute correlation and median shortest paths (along weighted edges from the partial correlation network) was compared to 1,000 permutations where assignment of the molecules shared by the categories was shuffled to calculate p-values and pairs of categories with q-values less than 0.025 were considered as distinct. Categories were excluded from analysis if they contained fewer than 10 members and were then divided into those which were internally consistent (median absolute correlation > 0.2; **bold**) and those where pathway members were scarcely correlated (*italics*). Edges are only shown when one or more pathways is coherent (bold) because it wouldn’t make sense to look at loss of coherence between already incoherent categories (i.e., italics-italics links).

**Figure SF27.**
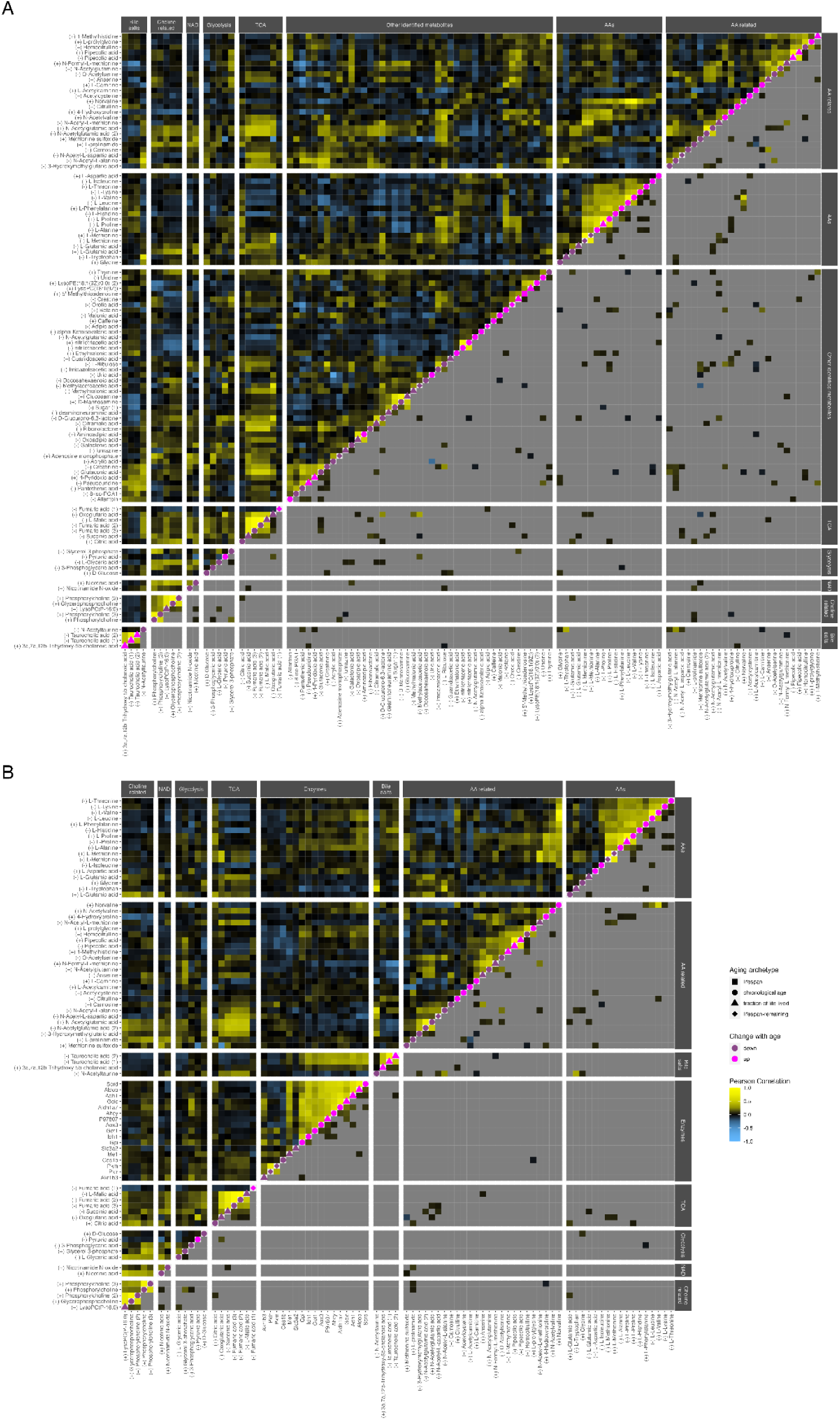
Extended metabolic changes with age. Dynamics (p)corr plots are described in Figure 6B. **(A)** Figure 6B including an additional “other” category including all aging-associated metabolomic changes falling outside of the emphasized categories. **(B)** Core metabolite changes from Figure 6B combined with an additional category of metabolic enzymes measured through proteomics.

**Figure SF28.**
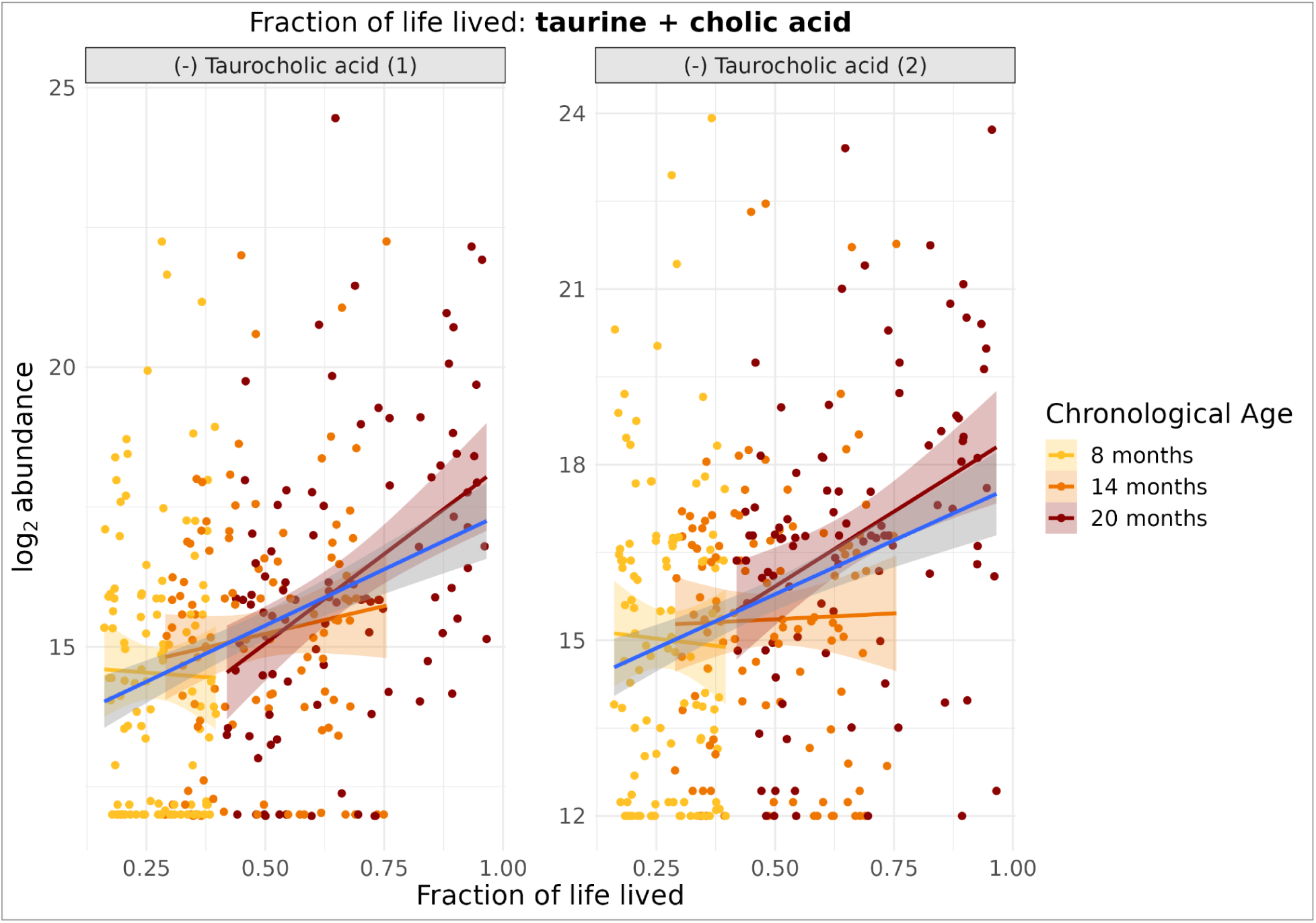
Taurocholic acid levels rise with fraction of life lived. Taurocholic acid (1) and (2) reflect separate peaks of the same compound.

**Figure SF29.**
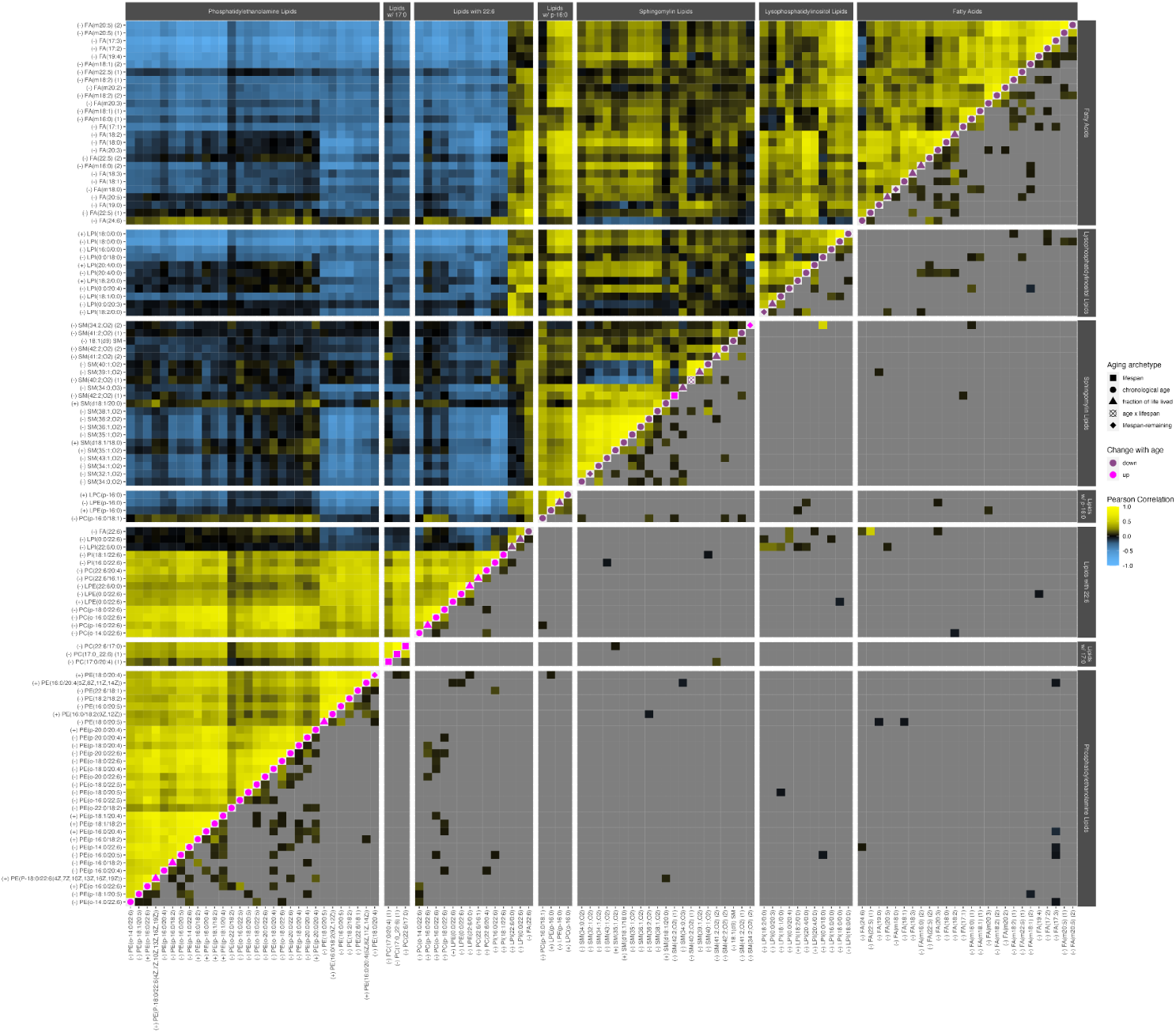
Chronological changes in the lipidome. Lipids which are part of a functional category enriched for chronological age are shown as a dynamics (p)corr plot as described in Figure 6B.

**Figure SF30.**
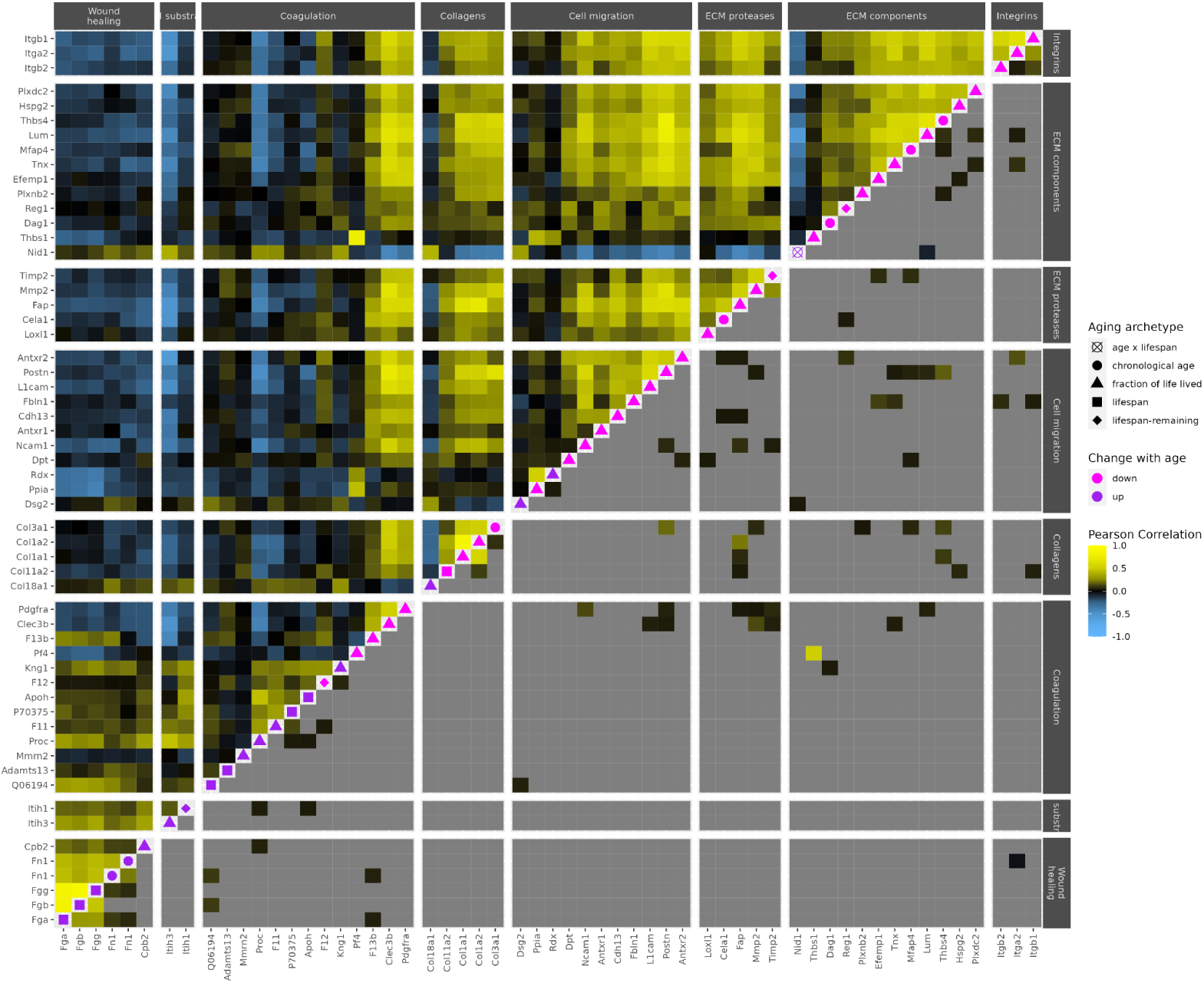
Matrisome proteins are depleted in circulation with age, while components promoting remodeling are elevated. The dynamics (p)corr plot is described in Figure 6B.

**Figure SF31.**
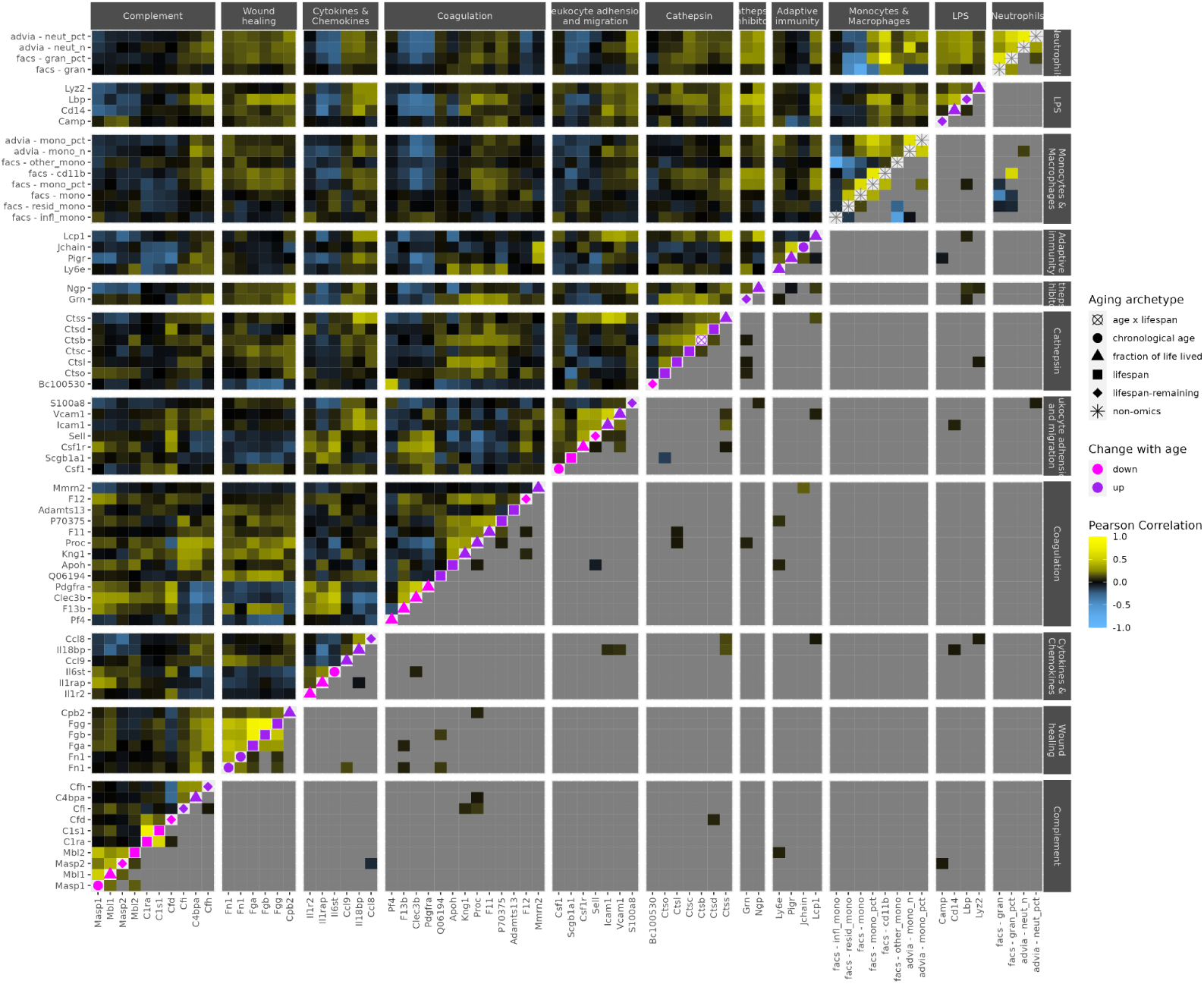
Toll-like receptor signaling, activated through Lbp and fibrinogens promotes monocyte recruitment and macrophage activation. The dynamics (p)corr plot is described in Figure 6B.

**Figure SF32.**
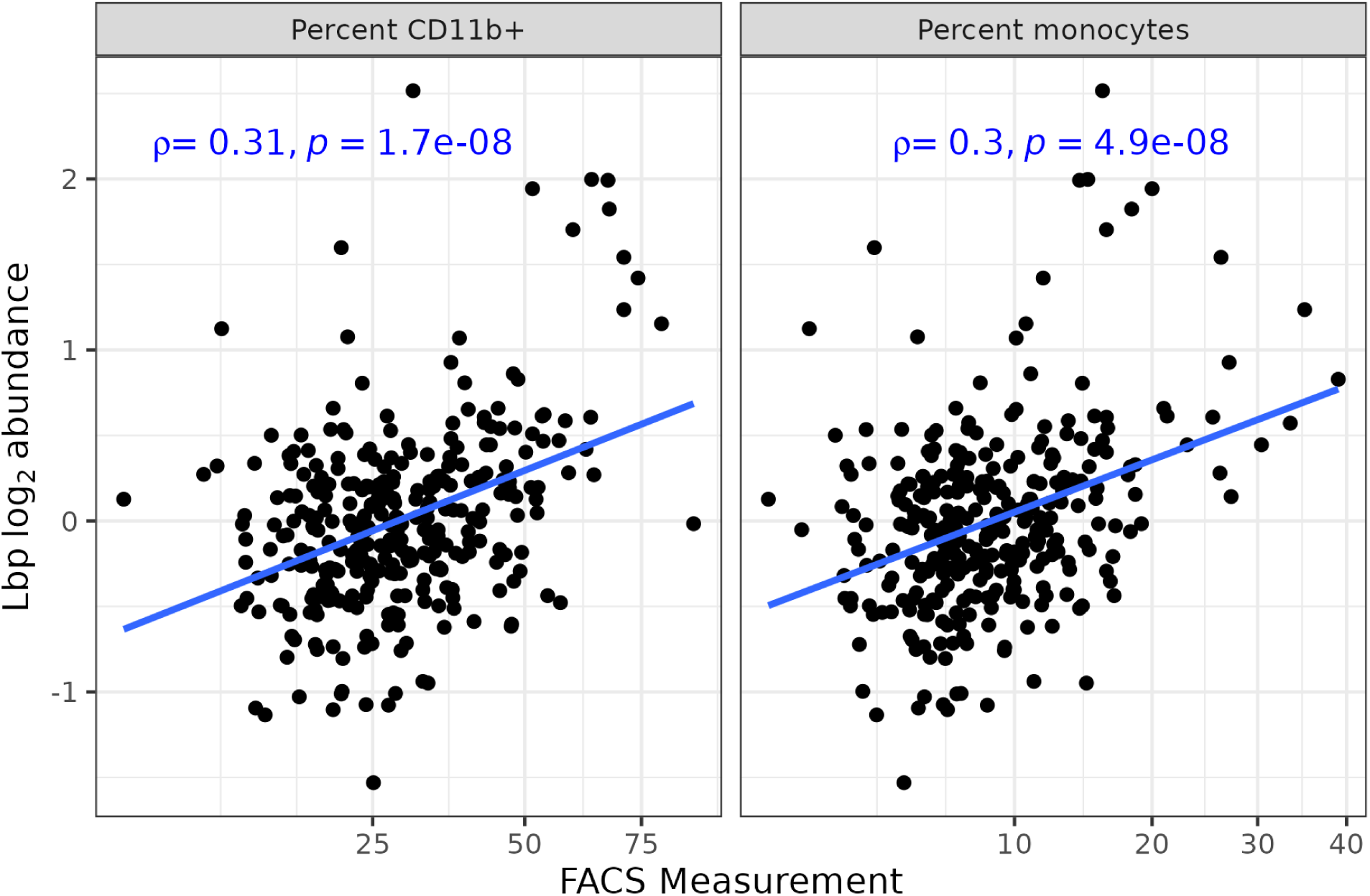
Comparing monocyte percentages and CD11b+ percentages based on FACS to LPS binding protein (Lbp) levels across samples.

**Figure SF33.**
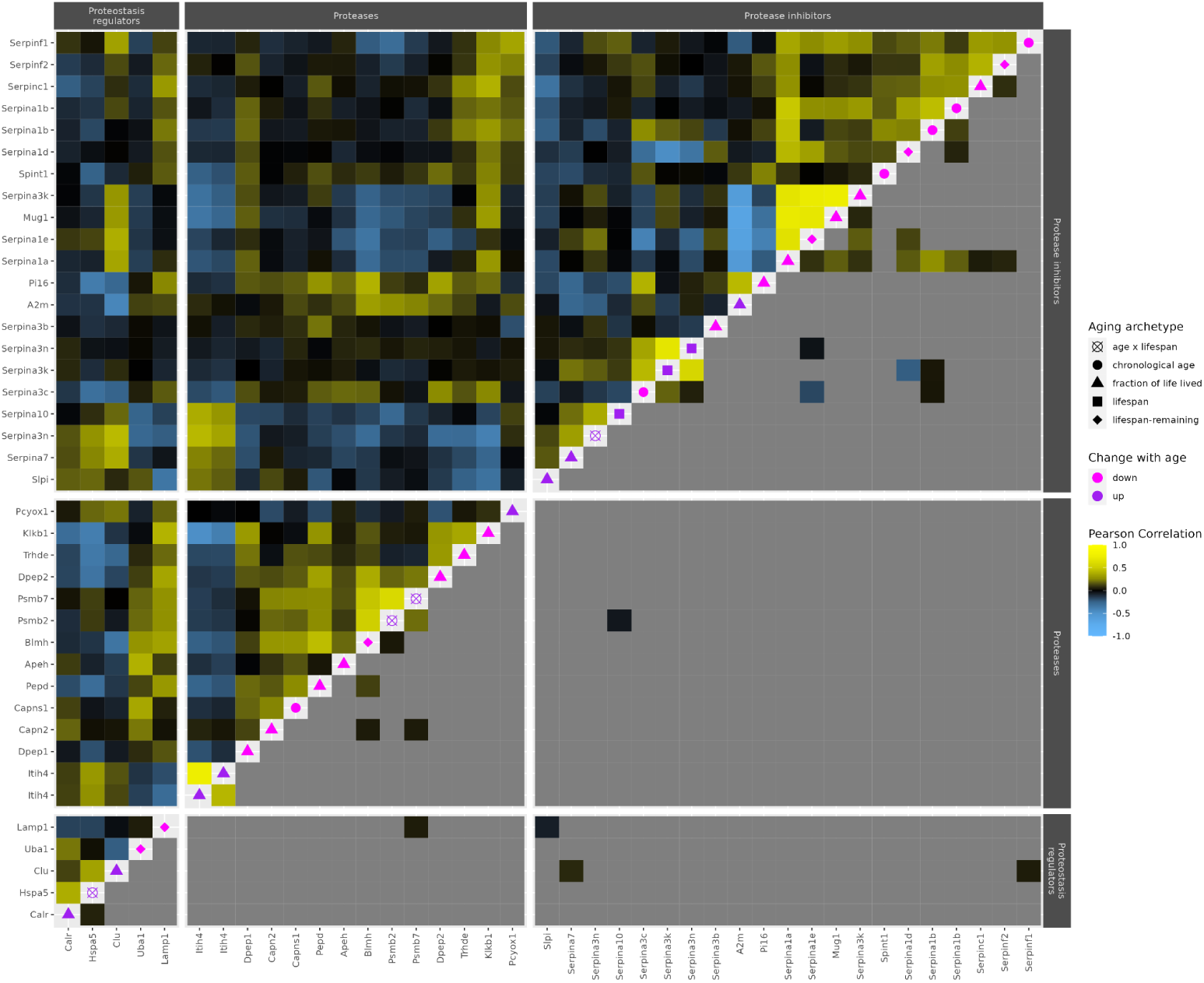
Circulating proteases and their regulators are broadly dysregulated during aging but with little directional coherence. The dynamics (p)corr plot is described in Figure 6B.

**Figure SF34.**
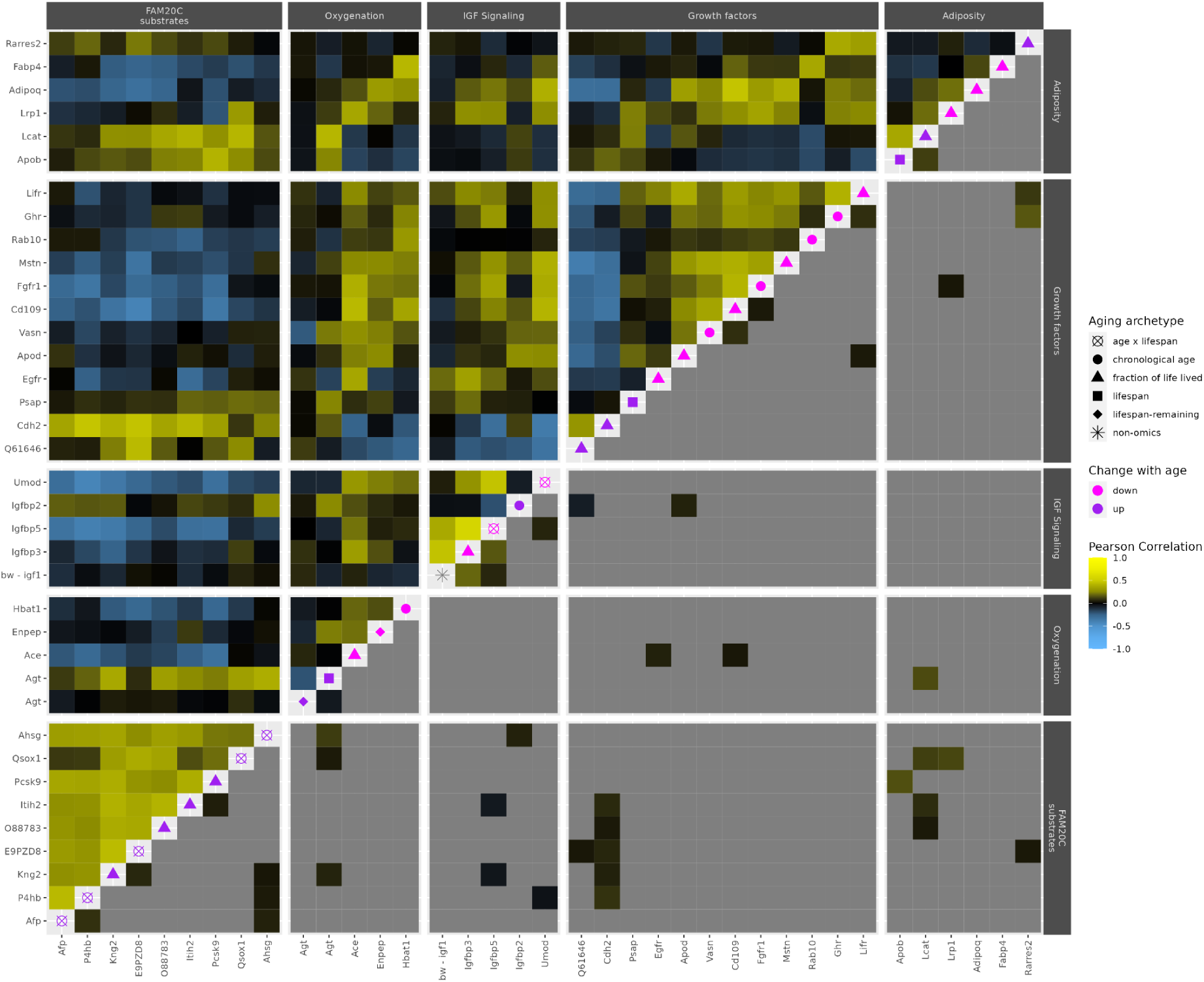
Aging affects mouse physiology spanning diverse homeostatic processes. The dynamics (p)corr plot is described in Figure 6B.

## Materials and Methods

### Sample generation

We enrolled 600 female DO mice in five waves, corresponding to birth cohorts of DO mice from generations 7 to 11. The first cohort entered the study in June 2011 and the study was fully populated in August 2012. This staggered enrollment allowed us to efficiently phenotype the cohort while simultaneously minimizing the confounding of seasonal and other time-specific batch effects with chronological age. The study included both male and female mice which were separately housed to avoid mating. Mice were housed 5 animals to cage in pressurized individually ventilated (PIV) polycarbonate cages supplied with high efficiency particulate air (HEPA) filtered air (Thoren Caging Systems Inc., Hazleton, PA, USA) maintained at temperatures ranging from 76 to 78°F. Mice were fed *ad libitum* with an autoclaved pelleted 6% fat diet (Lab diet 5K52, PMI Nutritional International, Bentwood, MO, USA). The mouse colony is maintained to avoid viral introduction and minimize exogenous stress as previously described ^21^. Environmental enrichments were provided including nestlets, biotubes, and gnawing blocks. Mice were maintained in these environments until they were either found dead, or euthanized following Jackson Laboratory’s standard practice for animal husbandry.

At 8, 14, and 20 months mice were weighted, urine was collected and 250 𝜇L whole blood was collected by retro-orbital bleed into heparin-coated micro-hematocrit tubes. Whole blood was immediately profiled. Plasma was extracted by centrifugation at 20,000 g for 10 min at 4°C and banked for future applications ^21^.

Blood glucose was profiled by OneTouch Ultra2 Blood Glucose Monitoring System Lot Number: Y15100068X and Igf1 levels were measured by ELISA (Mouse/Rat IGF-I Quantikine ELISA Kit, MG100, R&D Systems) ^21^. Blood urea nitrogen (BUN), magnesium, potassium, and sodium were measured using a Beckman Coulter AU680 Analyzer. Urine albumin, creatinine, magnesium, and phosphate were measured by Beckman Coulter AU680 Analyzer. Whole blood was profiled using two hematological analyzers, complete blood counting (CBC), and the ADVIA, Siemens ADVIA 2120 system to characterize the abundances and distributions of red, white blood cells, and platelets. A white cell differential as part of the CBC provided additional information about the distribution of white blood cells based on standard surface markers.

Gross phenotypes pertaining to the 110 mice in this study are included as part of this dataset, while phenotyping of the remaining mice will be presented in subsequent works outlining broader findings from the SHOCK study.

#### Selection for multiomics

Of the 600 mice enrolled in the SHOCK study, 455 lived past their third blood draw. We considered the mice who reached this threshold as eligible for our longitudinal profiling, and selected a subset of 110 of these mice enriching for both short- and long-lived mice for multiomics.

A selection of 54 mice were profiled in 2017, and a second tranche of 56 mice in 2019, to construct the full dataset of 110 mice profiled at three ages.

Tranche 1 was profiled as two sets of samples, set 1 and set 2. Because they were run near each other, possessed an identical experimental design, and were easily bioinformatically aggregated, leaving no evidence of systematic cross-set differences, we generally consider these two sets as a single tranche set 1/2. In contrast, set 3 was run at a later time than set1/2 with different sample preparation, a different blocking strategy and controls, and it required substantial effort to align its signals to those in set 1/2, as detailed below.

Within each set, samples were divided into a set of batches which were processed together, along with experimental controls. One of these controls was a positive-control reference sample, which was a common sample run in each batch to address batch-to-batch variability. This positive control served as an external control in metabolomics and lipidomics (i.e., it was run as a separate sample), and as an internal-control bridge channel in proteomics

Proteomics and metabolomics/lipidomics were profiled from separate aliquots of the same blood sample, with metabolites and lipids being separated from hydrophilic and hydrophobic fractions of the same sample. Proteins were tryptic digested and tagged with isobaric tandem mass tags (TMT) which allowed for pooling of 9-10 experimental samples with a standard bridge sample which facilitated cross plex comparison.

For set1/2, samples were split into 18 batches of 9 samples, each prepared on a separate day. Batches were created using a latin square, ensuring that no animal had more than 1 sample in a batch, and batches were balanced for age and survival. In this tranche, the positive control was young C57BL/6 blood, prepared fresh for each batch.

For set 3, samples were split into 7 batches of 24 samples, each prepared on a separate day. Batches were created, ensuring that no animal had more than 1 sample in a batch, and that the batches were balanced for age and survival. In this tranche, the positive control was a pooled sample with equal contributions of all 168 biological samples, prepared fresh for each batch.

#### Metabolomics and lipidomics sample preparation and LC-MS analysis

##### For the combined set 1/2

Separation of lipids and polar metabolites was performed by manual liquid-liquid extraction, a modification of the method published by Matyash et al. 2008 ^86^. Briefly, 1 mL of 1:1 MeOH:water was added to a 10 𝜇L aliquot of plasma in a 2 mL glass vial, followed by vortex mixing and incubation on ice for 10 minutes, addition of 1 mL of methyl,tert-butyl ether (MTBE), and further vortex mixing for 1 minute. Samples were then centrifuged at 3000 g for 5 minutes at 4°C to separate layers. Using a glass syringe, 750 𝜇L of the upper, organic layer was transferred to a destination vial, another 1 mL of MTBE was added, followed by vortex mixing and centrifugation as above, followed by removal of 1.4 mL of the organic phase, which was combined with the first organic phase layer. The lower, aqueous phase was transferred to a separate destination vial, leaving the protein pellet behind. The combined organic phase was evaporated to dryness overnight in a vacuum centrifuge maintained at 10°C, then resuspended in 1:1:0.3 chloroform:methanol:deionized water containing a 20-fold dilution of Splash Lipidomix (Avanti). The aqueous phase samples were evaporated to dryness overnight in a vacuum centrifuge maintained at 20°C and were then resuspended in 100 𝜇L of deionized water containing isotopically labeled internal standards (1 𝜇g/mL d4-lysine, 1 𝜇g/mL d5-phenylalanine and 0.25 𝜇g/mL d4 succinic acid). Aliquots of this solution were diluted 1:4 into deionized water or acetonitrile, for analysis by ion-pairing reverse-phase or HILIC LC-MS respectively.

##### For set 3

Separation of lipids and polar metabolites was performed by robotic liquid-liquid extraction, a modification of the method published by Matyash et al. 2008 ^86^, performed using a PAL DHR robot. Briefly, 20 𝜇L plasma aliquots were added to glass vials, and a 50% methanol solution containing containing 1 𝜇g/mL of deuterated tyrosine and deuterated alanine, 250 ng/mL of deuterated benzoate (Sigma Aldrich) and 20 𝜇L Splash Lipidomix mass spec standard (Avanti Polar Lipids) as internal standards was added to to a volume of 800 𝜇L, and the samples were vortexed for 70 seconds at 2000 rpm, followed by addition of 750 𝜇L of MTBE to each sample vial followed by vortex mixing for 30 seconds and centrifugation for 1 minute at 5000 rpm for complete phase separation. Next, 520 𝜇L of the upper (organic) phase was transferred to a lipidomics collection sample vial. The aqueous phase was re-extracted with addition of 600 𝜇L of MTBE to the source vial. After mixing and centrifugation as described above, 860 𝜇L of the top organic phase was transferred to the lipidomics collection vial. Then, 425 𝜇L of the lower aqueous phase was transferred to the metabolomics collection sample vials. The remaining solvent in the source sample vial, covering the protein pellet, had leftover volumes of around 100 𝜇L of the upper organic phase and 150 𝜇L of lower aqueous phase remaining that were not collected. Both organic and aqueous fractions were stored in blocks cooled to 4°C in the vial cabinet during the process. Both phases were dried under nitrogen gas, the aqueous phase was resuspended in 100 𝜇L of water containing 1 𝜇g/mL of deuterated lysine, deuterated phenylalanine and 250 ng/mL of deuterated succinate for the polar metabolites. Aliquots of this solution were diluted 1:4 into deionized water or acetonitrile, for analysis by ion-pairing reverse-phase or HILIC LC-MS respectively. The organic phase was resuspended in 2:1:1 butanol:methanol:deionized water for LC-MS analysis of lipids.

#### LC-MS analysis of polar metabolites

Metabolites were analyzed in positive ionization mode via HILIC chromatography using a SeQuant® ZIC®-pHILIC column, 5 μm particle size, 200 Å, 150 × 2.1 mm. Mobile phase A was 20 mM ammonium carbonate in water (pH 9.2); mobile phase B was acetonitrile. The flow rate was 150 μL/min and the gradient was t = −6, 80% B; t = 0, 80% B; t = 2.5, 73% B; t = 5, 65% B, t = 7.5, 57% B; t = 10, 50% B; t = 15, 35% B; t = 20; 20% B; t = 22, 15% B; t = 22.5, 80% B; t = 24; 80% B. The mass spectrometer was operated in positive ion mode using data-dependent acquisition (DDA) mode with the following parameters: resolution = 70,000, AGC target = 3.00E + 06, maximum IT (ms) = 100, scan range = 70–1050. The MS2 parameters were as follows: resolution = 17,500, AGC target = 1.00E + 05, maximum IT (ms) = 50, loop count = 6, isolation window (m/z) = 1, (N)CE = 20, 40, 80; underfill ratio = 1.00%, Apex trigger(s) = 3–10, dynamic exclusion(s) = 25.

Metabolites analyzed in negative ionization mode using a reverse phase ion-pairing chromatographic method with an Agilent Extend C18 RRHD column, 1.8 μm particle size, 80 Å, 2.1 × 150 mm. Mobile phase A was 10 mM tributylamine, 15 mM acetic acid in 97:3 water:methanol pH 4.95; mobile phase B was methanol. The flow rate was 200 μL/min and the gradient was t = −4, 0% B; t = 0, 0% B; t = 5; 20% B; t = 7.5, 20% B; t = 13, 55% B; t = 15, 95% B; t = 18.5, 95% B; t = 19, 0% B; t = 22, 0% B. The mass spectrometer was operated in negative ion mode using data-dependent acquisition (DDA) mode with the following parameters: resolution = 70,000, AGC target = 1.00E + 06, maximum IT (ms) = 100, scan range = 70–1050. The MS2 parameters were as follows: resolution = 17,500, AGC target = 1.00E + 05, maximum IT (ms) = 50, loop count = 6, isolation window (m/z) = 1, (N)CE = 20, 50, 100; underfill ratio = 1.0 0%, Apex trigger(s) = 3–12, dynamic exclusion(s) = 20.

#### LC-MS analysis of lipids

Two injections of each sample were performed, one for analysis is positive ion mode, and one in negative ion mode. Separation of lipid compounds in both positive and negative ion mode was achieved by reverse-phase liquid chromatography on an Accucore C30 column (250 x 2.1 mm, 2.6 µm particle size, Thermo Scientific) at 35°C column temperature with a 200 µL/min flow rate. Mobile phase A was 20 mM ammonium formate in 60:40 meCN:H_2_O, with 0.25 µM medronic acid and mobile phase B was 20 mM ammonium formate in 90:10 IPA:meCN, with 0.25 µM medronic acid. The gradient was: t= -7, 30% B; t=7, 43% B; t=12, 65% B; t = 30, 70% B; t=31, 88% B; t=51, 95% B; t=53, 100% B; t=55, 100% B; t=55.1, 30% B; t=60, 30% B for a total run time of 67 min per injection. The injection volume was 5 uL.

The mass spectrometer was a Q-Exactive Plus (Thermo) operated using data-dependent acquisition. The MS^1^ settings were: 140,000 resolution; AGC target of 3e6; max. IT of 100 ms; scan range of 200 to 2000 m/z. The MS^2^ settings were: 17,500 resolution; loop count 8; AGC target of 3e6; max. IT of 150 ms; isolation window of 1 m/z; and underfill ratio at 1%. Dynamic exclusion was set at 15 sec with an apex trigger from 5 to 30 sec. Stepped collision energies were set to 20, 30 and 40% NCE.

#### Metabolomics and lipidomics informatics

Metabolomics and lipidomics datasets were analyzed using OpenClam (https://github.com/calico/open_clam). The quahog R/python pipeline in OpenClam was used to process each dataset using a standardized workflow encompassing peak detection, alignment, and peak re-detection. Alignment was facilitated by detecting ∼20 unambiguous masses in each method where a single peak was found (see clamr::find_candidate_anchors()). The shift of such “anchor point” peaks relative to the peak group’s average retention time provided a measure of the retention time shift of each sample as a function of retention time. This measure was linearly interpolated to adjust each peak’s retention time. After adjusting peaks’ retention time using anchor points a second retention time aligner was used to further correct for cross sample retention time drift. This aligner groups fragmentation patterns across samples and then tries to minimize the cross-sample retention time disagreement of matched fragmentation patterns using a generalized additive model (see clamr::ms2_driven_aligner). Metabolomics results could be aligned across set12 and set3 using a combination of MS2 based alignment and anchor points. This allowed us to explore associations with unknown metabolites and cohort age and lifespan. Lipidomics results were alignable within set12 and set3 but non-monotonic shifts in retention time required us to separately annotate the knowns in each dataset and then merge results based on these labels. This prevented us from studying unknown lipids.

Each of 6 metabolomic/lipidomic datasets (positive and negative mode metabolites, and positive and negative mode lipids processed separately for set1/2 and set3) was manually curated using MAVEN ^31^. Metabolites were annotated based on the retention time of pure standards running on a matching chromatographic method, and based on a fragmentation pattern match to a pure standard fragmented at the collision energy of the experimental method. Lipids were matched to the published CalicoLipids *in silico* fragmentation library encompassing common classes of lipids ^31^. Due to the size of these complete datasets, a subset of samples covering each set and batch (nested within set) was curated and manual identities were then propagated to the complete dataset by matching the mass and retention time of curated peakgroups to those that were automatically detected. Curated peakgroups which were missed by automated peak picking were re-extracted as extracted ion chromatographs (EICs) in the complete dataset.

Automated peak picking and manual curation ensured that features could be consistently identified across all experimental samples by virtue of either shared mass and retention time in metabolomics, or shared identity in lipidomics. To interpret changes in abundance of each feature further requires us to (1) filter non-biological analytes, and (2) to minimize the impact of technical variability on variance and bias. These effects were addressed separately in each of 8 datasets (treating each of metabolites/lipids, positive-/negative-mode, set12/3 separately) to ensure that within-dataset batch effects were addressed before comparing across sets, modes, and data modalities. Each dataset was normalized and QC’d using a standard workflow (wdl_processes/featurization_small_molecules.Rmd) scheduled using [Cromwell](https://github.com/broadinstitute/cromwell). Results for all datasets can be found at [GitHub](https://github.com/calico/hackett-doomics).

First, non-biological unknowns (i.e., non-manually curated) were detected and removed if the median signal in biological samples was less than 4-fold greater than the median signal in negative controls (processed like biological samples but with no plasma) or if a peak was missing in more than 90% of biological samples. This identified a subset of features that ostensibly came from plasma. Before correcting for addressable batch effects, unknowns with egregious batch effects were discarded based on whether the standard deviation of batch mean(log2(abundance)) was more than 1.5.

To correct for batch-to-batch variability, we explored several approaches to normalization which performed a loading adjustment to samples ^40^ and/or features. The feature normalization strategies we explored were enforcing a constant batch median for each feature, subtracting each batch’s median positive control (the same biological sample run multiple times in each batch), and fitting and then subtracting a weighted LOESS curve over time (which upweighted positive controls). An ideal normalization approach would reduce the variability of positive controls (which are biologically identically, and hence variability is entirely technical) while maintaining much of the variability in biological samples (which vary due to both technical and biological variability). Based on these criteria loading adjustment was uniformly beneficial and the feature normalization strategies performed similarly (Figure <u>SF2</u>). Because the positive controls of the set1/2 were aged BL6 mouse plasma which was systematically different from the experimental samples, we opted to correct for feature-level batch effects using batch centering and we applied this uniformly across all datasets for simplicity. Because the major biological effects (age, sex, lifespan quartile) could be carefully balanced within each batch the batch median should be relatively independent of the specific biological samples in each batch and hence this should primarily remove technical variation. After normalization, we excluded additional unknowns where we would be underpowered to detect biological trends - those where either the standard deviation of positive controls was > 1.2 or the standard deviation of samples was less than 10% greater than the standard deviation of positive controls. To merge across set1/2 and set3, only unknowns which were separately retained in both sets were used, and features were median centered within each set; As for centering within batches this is broadly enabled by the similar composition of samples within each set. One caveat is that there were no DDM samples in set3 - this difference could affect mean centering but would have a minimal effect when centering by median.

#### Proteomics sample preparation and LC-MS analysis

10 µL of plasma were diluted with 290 µL 50 mM HEPES buffer (pH = 8.5). The disulfide bonds were reduced by adding dithiothreitol (DTT) to a final concentration of 5 mM and incubation at 56 °C for 30 min. Followed by adding iodoacetamide to adjust a final concentration of 15 mM and an incubation in the dark at room temperature for 20 min. The reaction was stopped by adding DTT to a final concentration of 5 mM and an incubation in the dark at room temperature for 15 minutes. 37.5 µL of 8 M Urea and digested overnight at room temperature with 1 µg/µL endoproteinase Lys-C (Wako) followed by digestion with sequencing-grade trypsin (Promega) at a final concentration of 1 ng/μL 6 h at 37 °C. The digestion was quenched with 1% trifluoroacetic acid (TFA), and peptides were desalted using Sep-Pak C18 solid-phase extraction (SPE) cartridges (Waters). The peptide concentration of each sample was determined using a BCA assay (Thermo Scientific).

For labeling with TMT-10plex reagents (Thermo Scientific), 50 µg of peptides were dried and resuspended in 50 µL of 200 mM HEPES (pH 8.5), 30% acetonitrile (ACN). Labeling was performed by adding 150 μg TMT reagent in anhydrous ACN and incubating at room temperature for 1 h. The reaction was stopped by addition of 5% (w/v) hydroxylamine in 200 mM HEPES (pH 8.5) to a final concentration of 0.5% hydroxylamine and incubation at room temperature for 15 min. Samples were acidified with 1% TFA, and samples were combined. The pooled samples were desalted using Sep- Pak C18 SPE cartridges. The combined multiplexed samples underwent a prefractionation by basic pH reversed chromatography (bRPLC) as previously described, collecting 96 fractions that were combined into 24 samples of which 12 were analyzed by mass spectrometry ^87^. The samples were dried and resuspended in 5 % ACN/5 % formic acid to be analyzed in 180-min runs via reversed-phase LC-M2/MS3 on an Orbitrap Lumos mass spectrometer (Thermo Fisher Scientific) using the Simultaneous Precursor Selection (SPS) supported MS3 method ^88,89^. The mass spectrometer was coupled to an Easy-nLC 1000 (Thermo Fisher Scientific) with a chilled autosampler. Peptides were separated on an in-house pulled, in-house packed microcapillary column (inner diameter, 100 μm). Columns were packed to a final length of 30 cm with GP-C18 (1.8 μm, 120 Å, Sepax Technologies). Peptides were eluted with a linear gradient from 11 to 30 % ACN in 0.125 % formic acid over 165 minutes at a flow rate of 300 nL/minute while the column was heated to 60 C. Electrospray ionization was achieved by applying 1800 V through a stainless steel T-junction at the inlet of the microcapillary column.

The Orbitrap Lumos was operated in data-dependent mode, with a survey scan performed over an m/z range of 500-1,200 in the Orbitrap with a resolution of 12×10^4^, automatic gain control (AGC) of 4 x 10^5^, and a maximum injection time of 50 ms. The most abundant ions detected in the survey scan were subjected to MS2 and MS3 experiments to be acquired in a 5 seconds experimental cycle. For MS2 analysis, doubly charged ions were selected from an m/z range of 600-1200, and triply and quadruply charged ions from an m/z range of 500-1200. The ion intensity threshold was set to 5×10^5^ and the isolation window to 0.5 m/z. Peptides were isolated using the quadrupole and fragmented using CID at 30 % normalized collision energy at the rapid scan rate using an AGC target of 2 x 10^4^ and a maximum ion injection time of 35 ms. MS3 analysis was performed using synchronous precursor selection (SPS) ^88,89^. Up to 5 MS2 precursors were simultaneously isolated and fragmented for MS3 analysis with an isolation window of 2 m/z and HCD fragmentation at 55 % normalized collision energy. MS3 spectra were acquired at a resolution of 5×10^3^ with an AGC target of 4.5 x 10^5^ and a maximum ion injection time of 150 ms.

#### Proteomics informatics

Mass spectrometry data were processed using an in-house software pipeline ^90^. Raw files were converted to mzXML files and searched against either a mouse Uniprot database using the Sequest algorithm. Database searching matched MS/MS spectra with fully tryptic peptides with a precursor ion tolerance of 20 p.p.m. and a product ion tolerance of 0.6 Da. Carbamidomethylation of cysteine residues (+57.02 Da) and TMT tags (+229.16 Da) on lysines and peptide N-termini were set as static modifications. Oxidation of methionine (+15.99 Da) was set as a variable modification. Linear discriminant analysis was used to filter peptide spectral matches to a 1 % FDR (false discovery rate) at the peptide level as previously described ^90^. Non-unique peptides that matched to multiple proteins were assigned to proteins that contained the largest number of matched redundant peptide sequences using the principle of Occam’s razor ^90^. Quantification of TMT reporter ion intensities was performed by extracting the most intense ion within a 0.003 m/z window at the predicted m/z value for each reporter ion. TMT spectra were used for quantification when the sum of the signal-to-noise for all the reporter ions was greater than 200 ^88^.

The fold-change of an experimental samples’ reporter ions to the bridge sample report ion was used as a measure of relative abundance. These relative abundances were centered so all sets possessed the same mean for every protein. Proteins which were present in more than 50% of the 330 experimental samples were considered further for statistical analysis.

### Genetics

We collected tail clippings and extracted DNA using DNeasy Blood and Tissue Kit (Qiagen) from 465 of the 600 animals enrolled in the SHOCK study. Samples were genotyped using the 77,808-probe MegaMUGA array from the Illumina Infinium platform ^91^. Genotype data is available at https://dodb.jax.org/, under project title: Shock Center Longitudinal Study.

We evaluated genotype quality using the R package: *qtl2* ^92^. We processed all raw genotype data with a corrected physical map of the MegaMUGA array probes (https://kbroman.org/MUGAarrays/muga_annotations.html, last accessed September 12, 2023). After filtering genetic markers for uniquely mapped probes, genotype quality and a 20% genotype missingness threshold, our dataset contained 72,370 markers. Two pairs of mice were found to have nearly identical genotypes, which suggested that one sample was mislabeled, and were removed. One additional mouse with more than 20% missing genotype data was also removed leaving a total of 460 mice with high quality genotype information. Of the 110 mice included in the multi-omic profiling, 86 had high quality genotype data.

Genetic mapping analyses were implemented in qtl2 ^92^ and followed the procedure described in Zhang et al. 2022 ^26^. Briefly, we calculated the probability that a given founder contributed a given marker allele by comparing each mouse’s genotype at each marker to that of the eight founder strains. Using these eight-state genotype probabilities, we calculated the pairwise relatedness (e.g. kinship matrix) between all mice and the narrow-sense heritability of all multi-omic traits. Additionally, we used the eight-state genotype probabilities to test for an association between the founder-of-origin probability and phenotype at all genotyped markers. We explored three types of mouse-level traits, an average effect which was simply the mean of a mouse’s blood draws, an early age effect (14 - 8 months), and a late age effect (20 - 8 months). In these QTL mapping analyses, the kinship matrix was treated as a random effect and sex and generation were included as fixed effects. Statistical significance for each trait was assessed via permutation analysis. Here, a trait’s p-value would be 1 minus the quantile of the observed maximum LOD score compared to the distribution of maximum LOD scores of individual null permutations.

To correct for multiple comparisons, we applied FDR-control to all traits p-value using the qvalue methodology, separately exploring association with each type of trait (i.e., average mouse abundances, and early- and late-age effects) ^93^. At FDR rates as high as 20% zero genetic associations with aging effects were retained, while we were able to detect hundreds of associations with average abundances at a 10% FDR. To interpret these associations for protein QTLs, we compared the genomic coordinate maximizing the LOD score in each QTL to the cognate protein’s start site, and defined all QTLs falling within 1 Mb of this start site as local QTLs.

### Statistical analysis

A set of 14 linear regression was fitted to each feature treating the features’ log_2_ fold-changes or log_2_ abundances as the dependent variable (see below). When identifying associations with age (early age, late age, age interactions, DDM), fold-changes of 14 month and 20 month timepoints relative to the corresponding 8 month observation from the same mouse were used to correct for cross-strain variability in baseline abundance. The 8 months observations were then excluded from regression (as their fold-change would be zero by construction). When identifying associations to mouse-level traits such as lifespan and sex, abundances were predicted since working with fold-changes would subtract out the signal of interest which would be shared across all of a mouses’ measurements.

For each model, major known biological and non-biological covariates were controlled for. Aside from the “fold-change lm (DDM)” model which aimed to identify molecules associated with imminent death, the ten 20 month observations from mice that would live less than 21 more days following their final blood draw were excluded from all models. This was because the DDM signature varied across the 10 DDM samples with some mice (notably a mouse who died two days after the blood draw) exhibiting profound changes in the multiome and other mice looking relatively normal despite their impending death. Filtering these samples was the safe option since failing to account for this variability could drive aging and lifespan associations which would not generalize across the cohort.

The other major biological effects identified through exploratory data analysis, age and sex, were included as covariates unless the covariates effect had already been addressed (i.e., sex’s effect was removed when working with fold-changes), or when a covariates inclusion would prevent the estimation of the effect of interest (i.e., we couldn’t correct for age when assessing associations with lifespan remaining). Besides biological covariates, the two generation x blood draw date batch effects, the early generation 8 effect and late blood draw effect were included as categorical variables. The formulation of these effects was slightly different based on whether a regression used fold-changes or abundances. For example, because the 8 and 14 month measurements of early generation 8 mice were similarly affected by the EG8 batch effect this effect was removed when taking their fold-change difference.

The 14 models accounting for these considerations are:

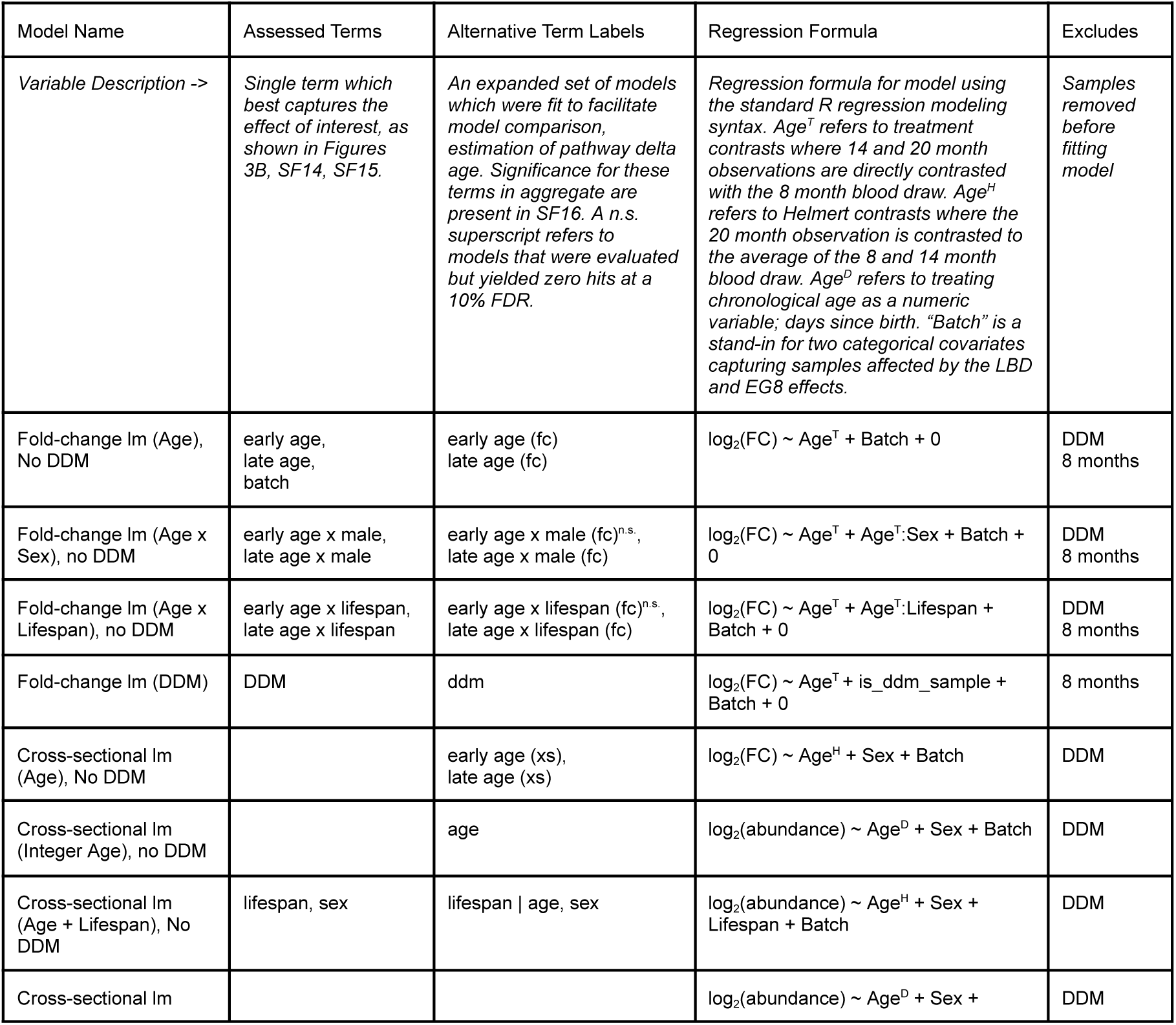
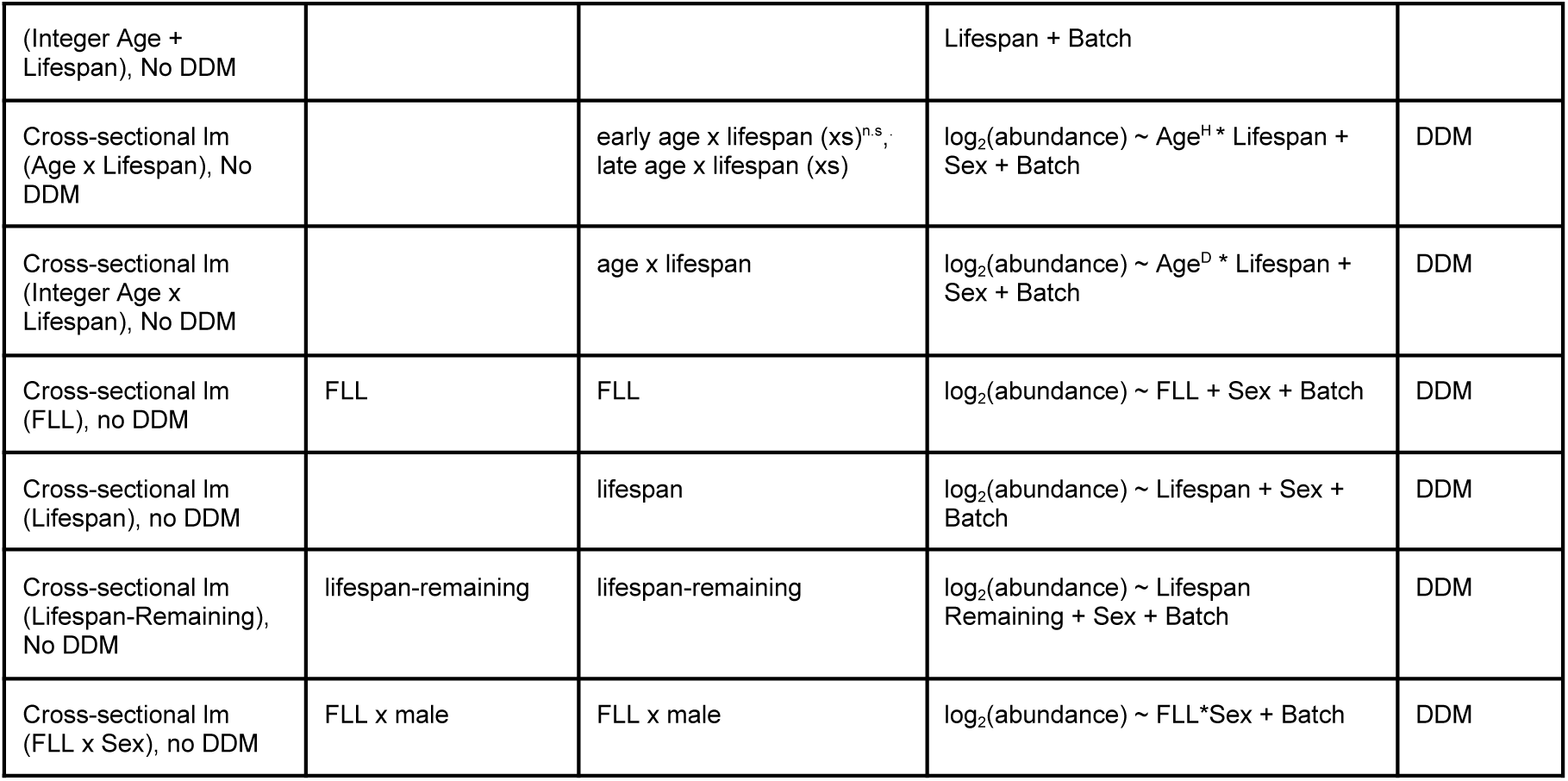

Each of the above models was separately fit to every molecule and significance was evaluated by bootstrapping residuals using a non-parametric bootstrapping procedure ^39^. Briefly, an initial regression was fit and its residuals were inflated to adjust for the fitted degrees of freedom (ε*^bootstrap^* = ε(𝑁/(𝑁 − 𝑑. 𝑜. 𝑓)). Then for each of 1e6 bootstraps, residuals were samples with replacement and added to the 𝑦 values from the initial regression, and an identical regression was fit to each bootstrap to estimate to generate a bootstrapped distribution of effect sizes.

Two-tailed p-values can then calculated as 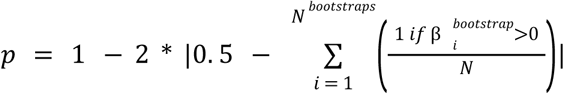. We then added 1/𝑁^𝑏𝑜𝑜𝑡𝑠𝑡𝑟𝑎𝑝𝑠^ to each p-value and p-values greater than one were set to one. This lower bounds each term’s nominal significance to 1/𝑁^𝑏𝑜𝑜𝑡𝑠𝑡𝑟𝑎𝑝𝑠^ with such values reflecting that zero conformations of the bootstrapped residuals were able change the sign of β’s relation to ŷ. Missing values, which are common in TMT proteomics, were addressed by excluding the relevant sample from statistical analysis.

To correct for multiple comparisons, we estimated a single value of π_0_ (an estimate of the fraction of molecules which do not exhibit differential abundance) for each term irrespective of data modality ^93^. We then applied the q-value FDR procedure to each data modality separately to identify discoveries at a 10% FDR using a π_0_estimate that was shared across data modalities. This reflects that π_0_ estimates are similar across data modalities but our power nonetheless differs by data modality due to the higher accuracy of the proteomics data. FDR controlling all data modalities together using a single π_0_would decrease the number of proteomics hits (adding false negatives) while simultaneously selecting more metabolites and lipids (increasing false positives).

As an alternative approach we explored the use of linear mixed-effects models for accounting for repeated measures of the same mouse in cross-sectional data but ultimately decided against treating mouse as a random effect. This was primarily because linear mixed-effects models generated pathological results when applying nested model comparison to assess the significance of variables which are a property of individuals (e.g., lifespan and sex). Taking lifespan as an example, comparing a model with lifespan + (1|mouse) to a model with just (1|mouse), the poorer prediction of lifespan is largely absorbed by an elevated mouse-effect random effect standard deviation. This results in a pathological p-value distribution with an excess of p-values near 1. This property would prevent us from uniformly treating all aging archetypes, and beyond this, we don’t believe a random effect is generally needed because mouse-effects are small as can be seen from the low H^2^ of most molecular features.

To identify functional enrichments, we explored whether significant hits were enriched among common biological processes for proteins and whether lipids possessed common headgroups and/or acyl chains. For proteomics results we tested the biological processes and Reactome gene ontologies drawn from MSigDB ^94^ and filtered each category to proteins measured in our study. We focused on these ontologies over alternatives such as those focused on cellular compartments or molecular functions because plasma is acellular and most molecular functions are poorly represented in the plasma proteome. Lipid headgroups and tails were extracted based on the lipid naming scheme using the claman R package (https://github.com/calico/claman). To identify functional enrichments we applied a Fisher Exact test to each pair of terms (from regression) and category to identify significant overlaps between category membership and FDR-controlled discoveries (q < 0.1). P-values for each term were FDR controlled using the Benjamini-Hochberg procedure ^95^ to identify functional enrichments at a 10% FDR. Significant protein sets were further refined by removing genesets which are defined by similar membership. This was done by retaining a set of genesets with pairwise Jaccard similarity (intersection / overlap) of less than 0.7 favoring categories with higher Fisher Exact significance over categories with weaker functional enrichment.

A small number of lipid unknowns were identified in a preliminary analysis of set1/2 and were manually identified in the final set1/2 and set3 dataset. Since these unknowns were selected based on their strong association with age and/or lifespan in set1/2 we excluded them from FDR control to avoid contaminating the remaining features which were curated independently of their aging trends. We report a small number of these unknown’s associations based on their strong nominal significance (p < 0.0001) for exploratory purposes.

#### Power analysis

To determine the effect sizes that we are powered to detect for several of the aging patterns that we expect a priori in our study, changes with chronological age, lifespan, and age x lifespan interactions, we applied a simulation of our power to detect effects as a function of varying levels of noise.

When exploring changes with age, the comparison of a 14 or 20 month measurement to its 8 month baseline increases noise (but decreases bias) because technical variance excessed the biological variance (i.e., the variability among biological samples 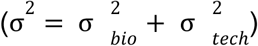 is less than twice that of positive control technical replicates 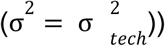

### Data interpretation

#### Model comparison

To identify the relative support of individual features and pathways for each aging archetype we used Akaike Information Criteria with a correction for finite sample sizes (AICc) to compare alternative models fit to the same data. Each model was fit to the 320 non-DDM cross-sectional samples, providing that it contains an aging archetype term among the “alternative term labels”. The one exception was the “Cross-sectional lm (Lifespan), no DDM” which we dropped because the “lifespan” effects were correlated with “lifespan | age”, but age was deemed to be an important covariate in capturing lifespan in a non-pathological manner.

For a single feature, the relative support for alternative models is proportional to *e*^𝐴𝐼𝐶/2^. This quantity can be rescaled by the sum of relative supports which allows us to treat the supports as a simplex bounded on [0, 1] and summing to a 1 over all models.

AIC is a combination of a Gaussian log-likelihood summed across samples, 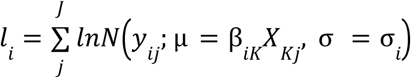, and a penalization for model complexity (2|K|). Just as log-likelihoods can be summed across samples, they can be summed across features, and similarly the total number of parameters used to fit all features is the sum of parameters for each feature. Following this logic that we are combining features and samples by aggregating their log-likelihoods and number of parameters we could aggregate AIC measures and then apply the correction for finite sample size (i.e., adjusting AIC to calculate AICc). But, this correction would dissipate as a larger number of observations are utilized in aggregate, and we felt that this feature was inappropriate because feature level regressions are independent hence the aggregate measure should be independent combinations of marginal probabilities. We retain this feature by summing AICc over features.

While this approach could be applied to any set of features assessed by a common set of regressions to determine which model is best supported overall, we chose to aggregate over pathways with functional enrichments. For each pathway we took all members, irrespective of their significance, and summed model AICc over all features.

#### Pathway delta age

To identify samples featuring premature or delayed changes in age-associated pathways, we explored the degree to which we could estimate an individual’s age based on biochemical pathways. Here, we are making the assumption that for a sample *j*, the biological age for a pathway is constant (𝑎_𝑗_ + δ_𝑗_) but this age may deviate from a sample’s known age (𝑎_𝑗_). For intuition, if a pathway was characterized by a constant increase in a set of molecules proportional to chronological age, then a sample with a higher level of these molecules than their age could be considered as having an elevated delta age (i.e., δ > 0). 𝑎_𝑗_ could be the chronological age of an individual at the time of sampling but could also be any other aging archetype. In these cases, pathway delta age may indicate the degree to which a sample either does or does not exhibit an aging pattern which is observed at a population level but may not be observed in some mice.

To estimate δ for a sample across all members of a pathway 𝑖 ⊂ 𝐼, we can write an extended form of the true model that we posit for describing a molecule’s dynamics.

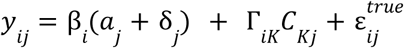

This model includes a sample’s measured age (𝑎_𝑗_), a to-be inferred delta age (δ_𝑗_) both of which are scaled by a feature-specific aging slope (β_𝑖_). Because age needs to be a continuous variable, we used aging regression models which treated age and age x lifespan as numeric rather than categorical variables for this analysis. 𝑦_𝑖𝑗_ is the measured level of molecule *i*, while Γ_𝑖𝐾_ is all other covariates’ regression coefficients which are scaled by covariates 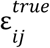 is the irreducible error which we could achieve after correcting for d.

This formula can be rearranged to relate previously determined regression residuals to feature slope (β_𝑖_), delta age, and irreducible error.

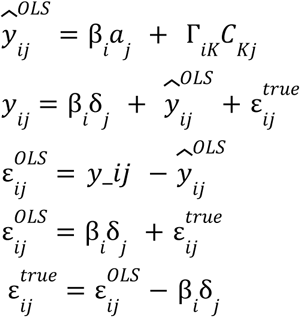

For a single feature we could solve for δ directly but the result would be meaningless because any feature could take on a different age which set 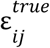 to zero. To avoid this conundrum constrain delta age (δ_𝑗_) to remain fixed across a set of features (𝑖 ⊂ 𝐼). Here, we considered the notion that each pathway could have a separate biological age and hence we estimated a delta age (δ_𝑗_), and by extension biological age (𝑎_𝑗_ + δ_𝑗_), which was constant across a pathway.

This can be solved as a least-squares problem where we minimize the total sum of squares of the irreducible error:

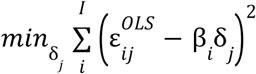

This is just simple least squares regression and hence we can identify δ by regressing residuals 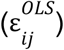 on aging slopes β_𝑖_.

#### Estimating partial correlations

To estimate sparse partial correlations we applied graphical LASSO (GLASSO) using the EBICglasso function from the R qgraph package. GLASSO approaches estimating the sample precision matrix (the inverse of the sample covariance matrix) by balancing predictive accuracy (of observed Multivariate Normal observations based on the estimated precision matrix), while shrinking individual coefficients towards zero with an L1 penalty (i.e., sum(abs(coef))) weighted by a regularization parameter lambda. While GLASSO requires a positive semi-definite (PSD) covariance matrix, due to missing measurements, the sample covariance matrix possessed negative eigenvalues. To address this issue we imputed missing values using k-nearest neighbor imputation (with K set as 10). Having done this, the sample covariance was PSD enabling us to apply GLASSO. We explored a range of lambda values with large values of lambda resulting in few non-zero coefficients and hence a nearly diagonal reconstructed covariance matrix. In contrast, small values of lambda will generate a near-perfect reconstruction of the sample covariance matrix. Neither extreme is biologically informative, however, intermediate values of lambda will select individual coefficients which are necessary to reconstruct sample correlations. We identified an appropriate intermediate value of lambda by minimizing the Bayesian Information Criteria (BIC) over values of lambda (i.e, *k*ln(*n*) - 2*logLik). BIC penalizes model complexity less heavily than AIC but we found that it was appropriate for this problem. The fitted precision matrix based on the value of lambda minimizing the BIC can be inverted to generate a covariance matrix which describes much of the covariance across molecules. The top 10 principle components of the sample covariance matrix capture 80% of the matrix’s variance, while the reconstruction error of the covariance matrix based on GLASSO equates to around 75% of variance explained as unexplained. While the precision matrix is directly estimated using GGLASSO we found partial correlations were easier to intuit since they are bounded on [-1,1] and capture the dependence between variables irrespective of the variance of these features. The formula to convert from the precisions to partial correlations is 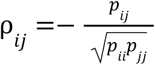 where *i* and *j* index features in the precision matrix.

